# Reconstructing the lineage histories and differentiation trajectories of individual cancer cells in *JAK2*-mutant myeloproliferative neoplasms

**DOI:** 10.1101/2020.08.24.265058

**Authors:** Debra Van Egeren, Javier Escabi, Maximilian Nguyen, Shichen Liu, Christopher R. Reilly, Sachin Patel, Baransel Kamaz, Maria Kalyva, Daniel J. DeAngelo, Ilene Galinsky, Martha Wadleigh, Eric S. Winer, Marlise R. Luskin, Richard M. Stone, Jacqueline S. Garcia, Gabriela S. Hobbs, Fernando D. Camargo, Franziska Michor, Ann Mullally, Isidro Cortes-Ciriano, Sahand Hormoz

## Abstract

Some cancers originate from a single mutation event in a single cell. For example, blood cancers known as myeloproliferative neoplasms (MPN) are thought to originate through the acquisition of a driver mutation (most commonly *JAK2*-V617F) in a hematopoietic stem cell (HSC). However, when the mutation first occurs in individual patients and how it impacts the behavior of HSCs in their native context is not known. Here we quantified the impact of the *JAK2-*V617F mutation on the proliferation dynamics of HSCs and the differentiation trajectories of their progenies in individual MPN patients. We reconstructed the lineage history of individual HSCs obtained from MPN patients using the patterns of spontaneous somatic mutations accrued in their genomes over time. Strikingly, we found that the *JAK2-*V617F mutation occurred in a single HSC several decades before MPN diagnosis — at age 9±2 years in a 34-year-old patient, and at age 19±3 years in a 63-year-old patient. For each patient, we inferred the number of mutated HSCs over time and computed their fitness. The population of *JAK2*-mutated HSCs grew exponentially by 63±15% and 44±13% every year in the two patients, respectively. To contrast the differentiation trajectories of the *JAK2*-mutated HSCs with those of healthy HSCs, we simultaneously measured the full transcriptome and somatic mutations in single hematopoietic stem and progenitor cells (HSPCs). We found that the fraction of *JAK2*-mutant HSPCs varied significantly across different myeloid cell types within the same patient. The erythroid progenitor cells were often entirely *JAK2*-mutant, even when the peripheral blood *JAK2*-V617F allele burden was low. The novel biological insights uncovered by this work have implications for the prevention and treatment of MPN, as well as the accurate assessment of disease burden in patients. The technology platforms and computational frameworks developed here are broadly applicable to other types of hematological malignancies and cancers.

## INTRODUCTION

Philadelphia chromosome-negative myeloproliferative neoplasms (MPN) are blood cancers often caused by a single nucleotide change in the *JAK2* gene (*JAK2*-V617F mutation) that disrupts normal blood production by activating JAK2 signalling^1–4^. Constitutive activation of JAK2 results in increased production of mature blood cells of the myeloid lineage, with some MPN patients presenting primarily with increased numbers of red blood cells (polycythemia vera, PV), others with increased numbers of platelets (essential thrombocythemia, ET), and more rarely some with scarring (“fibrosis”) of the bone marrow (primary myelofibrosis, PMF). While it has been shown that the *JAK2-*V617F mutation is detectable in hematopoietic stem cells (HSCs)^5^ and in all mature cell lineages^6,7^, it is unclear how the mutation impacts HSC differentiation and proliferation *in vivo* in individual patients. Therefore, to understand the natural history of *JAK2-*V617F mutated HSCs and progression to clinical disease, we studied unrelated patients with newly diagnosed *JAK2*-V617F+ ET and PV and asked when the *JAK2*-V617F mutation first occurred in each patient, how the number of *JAK2*-mutated cells expanded over time, and the extent to which the differentiation trajectories of the *JAK2*-mutated cells deviated from those of healthy cells.

The average human generates hundreds of billions of new blood cells every day to replace dying cells^8^. To sustain this flux of new cells, HSCs differentiate into hematopoietic progenitor cells that proliferate and produce terminally-differentiated blood cells^9^. To date, assessment of the functional consequences of *JAK2*-V617F on human HSCs has been limited to *in vitro* assays and xenotransplantation studies in immunocompromised mice. Studies *in vitro* have found that *JAK2*-mutated progenitor cells are hypersensitive to erythropoietin, exhibit a proliferative advantage when cultured^4,10^ and undergo a larger number of cell divisions compared with wild-type (WT) progenitor cells^11^. However, these observations may not represent the *in vivo* behavior of mutated progenitor cells, for example, the cytokine concentrations levels may not match the physiological levels. Xenotransplantation studies^12,13^ have also been limited because many human cytokine receptors that are expressed on transplanted CD34+ cells are not compatible with murine cytokines that are produced by the recipient murine bone marrow. Therefore, the experimental systems used in these studies do not accurately recapitulate the microenvironment or the behavior of *JAK2*-mutant hematopoietic stem and progenitor cells (HSPCs) in patients.

In addition to differentiating into lineage-restricted progenitors, an HSC can also divide symmetrically into two HSCs or be removed from the stem cell pool by death or by differentiating to a progenitor cell^14–16^. Over time, the number of HSCs descended from a particular HSC (including the *JAK2*-mutated HSC) fluctuates through random cell division and cell death events; descendants of some HSCs might take over the population whereas those of others dwindle and eventually disappear^17–19^. Do *JAK2*-mutated HSCs have a selective advantage over wild-type HSCs in human bone marrow? Does the number of *JAK2*-mutant HSPCs increase in MPN patients due to a series of random cell division and death events driven by neutral drift or as a result of a deterministic expansion process driven by positive selection?

To answer these questions, we reconstructed lineage trees of *JAK2*-mutated and healthy HSCs obtained from individual MPN patients and inferred the history of disease progression in 2 patients with ET. In addition to identifying how the *JAK2* mutation impacts the differentiation trajectories of the progenies of HSCs, we profiled the transcriptomes of individual cells obtained from bone marrow aspirates of 7 MPN patients. We used single cell profiling because bulk measurements average the transcriptomes of mutated cells and healthy cells and obscure different cell states along the differentiation trajectory. Importantly, our approach does not utilize *in vitro* cell culture or xenograft models of *JAK2* mutated HSPCs but instead directly measures the behavior of the mutated hematopoietic cells in their native context in patients.

## RESULTS

To investigate the effect of *JAK2* mutations in ET and PV patients, we performed single-cell transcriptomic and whole-genome sequencing of HSPCs from 7 newly diagnosed, untreated PV (*n*=3) and ET (*n*=4) patients, as well as healthy controls (*n*=2) (Fig. 1). The *JAK2*-V617F mutation was detected in 6 patients, while the remaining ET patient had a *JAK2* variant previously unreported in humans (*JAK2*-V617L)^20^, with fibroblast germline testing confirming a somatic origin for the mutation. The ET patients did not harbor additional myeloid-malignancy-associated mutations in their peripheral blood as measured by a clinical next-generation sequencing (NGS) assay (i.e. Rapid Heme Panel^21^), whereas somatic truncating mutations in *TET2* (2 patients) and *EZH2* (1 patient) were identified in the PV patients (Fig. 1b). From each MPN patient and healthy donor, we collected a bone marrow aspirate, isolated mononuclear cells, and then enriched for CD34 expression to isolate HSPCs (Methods).

**Figure 1.**
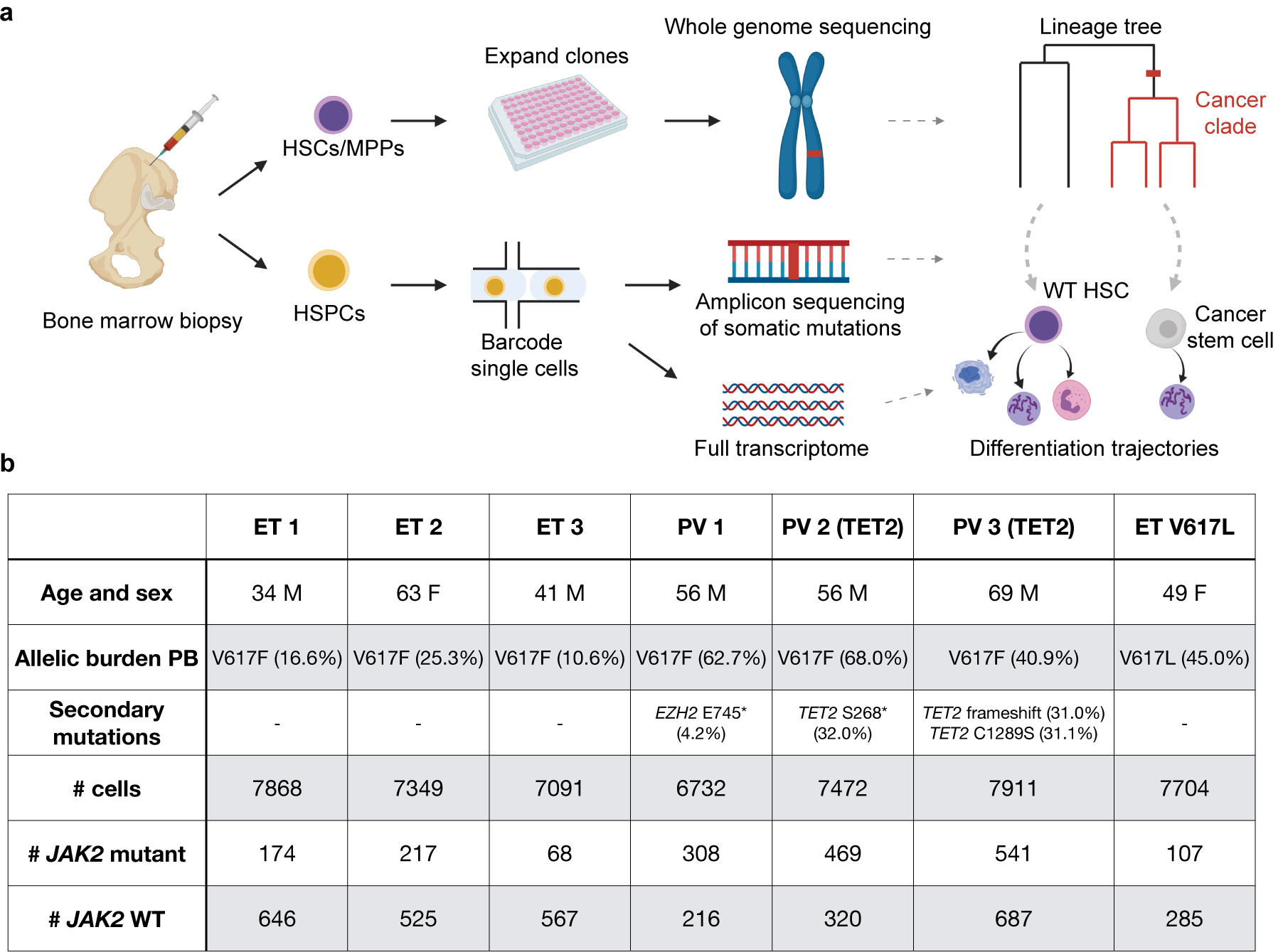
Experimental design. **a**. Individual hematopoietic stem and progenitor cells (HSPCs) from bone marrow aspirates of MPN patients were analyzed in two ways. First, hematopoietic stem cells (HSCs) and multipotent progenitors (MPPs) were expanded *in vitro* and characterized using whole-genome sequencing. Second, we simultaneously read out the transcriptional profiles and somatic mutations in single HSPCs. **b**. Information on MPN patients sampled in this study. “Allelic burden PB” and “secondary mutations” refer to variant allele fractions of *JAK2* mutations and other hematopoiesis-associated mutations in peripheral blood, respectively. The numbers of *JAK2* WT and mutant cells identified in the HSPCs using scRNA-seq are given in the last two rows.

### *JAK2-*mutant HSPCs exhibit fate bias

To determine how *JAK2* mutations affect HSPC differentiation dynamics in MPN patients, we simultaneously measured the full transcriptome and genotyped the *JAK2* mutation in individual CD34+ cells obtained from each bone marrow aspirate (Fig. 1a, Methods). To do so, we developed a protocol for amplifying specific transcripts from single-cell RNAseq libraries. Briefly, we used the 10X platform to generate barcoded single-cell cDNA libraries. Before fragmenting the libraries for sequencing, we generated amplicon libraries of the target loci for the somatic mutations of interest, by performing three rounds of nested PCR with locus-specific reverse primers and generic forward primers (SI Fig. 1, Methods). Next, we sequenced each of the single-cell libraries to obtain the transcriptomes and each of the amplicon libraries to obtain the somatic mutations. The somatic mutations were mapped to the transcriptional profiles using the shared single-cell barcodes across the two libraries (SI Fig. 2, Methods).

**Figure 2.**
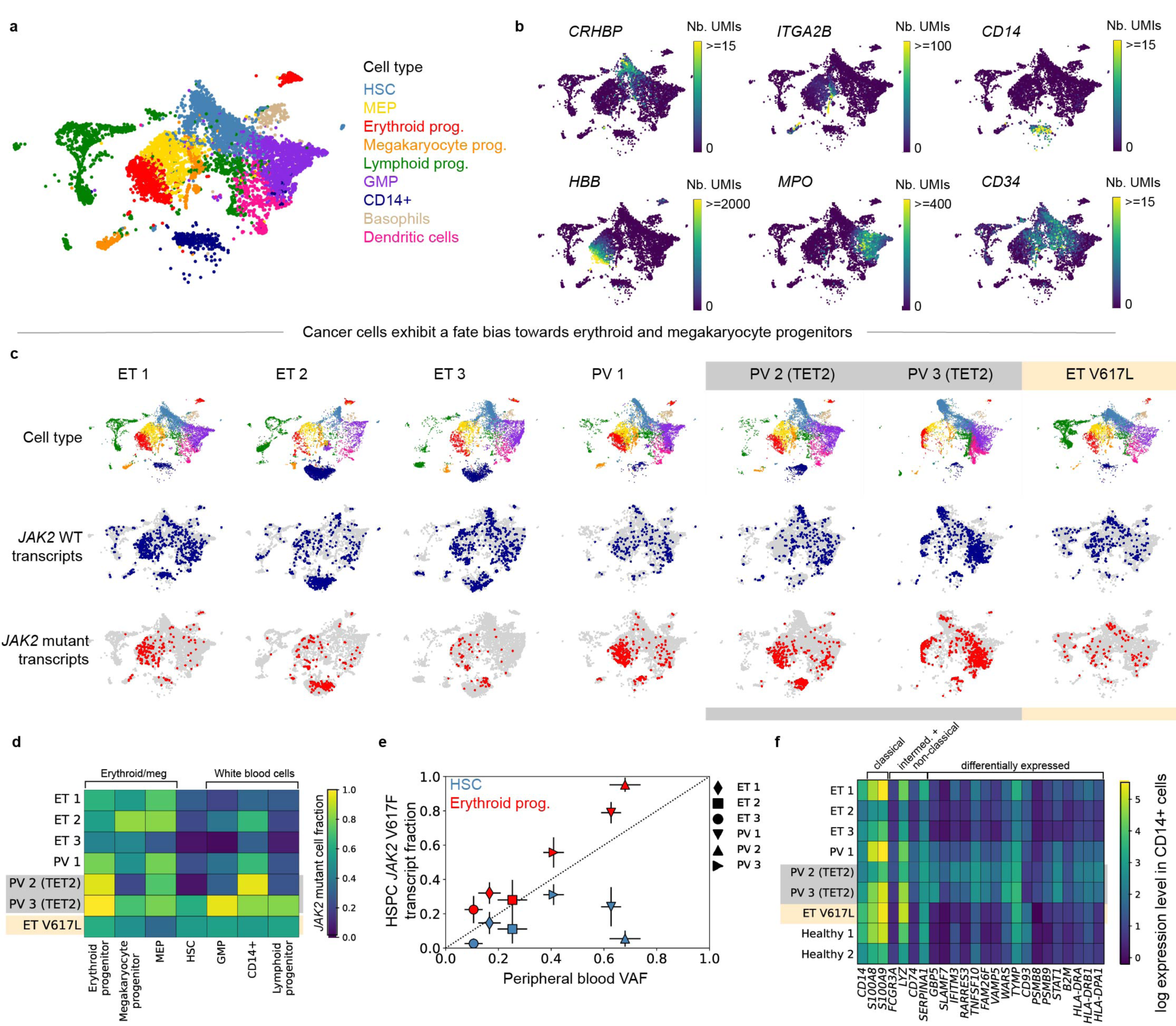
Erythroid and megakaryocyte progenitors from MPN patients are more likely to have the *JAK2*-V617F mutation than other CD34+ bone marrow HSPCs. **a**. UMAP of CD34-enriched bone marrow scRNA-seq data from ET 1, colored by cell type. **b**. Marker gene expression in ET 1 CD34-enriched bone marrow. **c**. Cell type classifications and *JAK2* WT/mutant transcript calls in scRNA-seq data from individual MPN patients (columns). **d**. Fraction of *JAK2* mutated cells (colors) in different bone marrow cell types from individual MPN patients. **e**. Relationship between peripheral blood VAF and *JAK2* V617F mutant transcript fraction in bone marrow HSCs (blue) and erythroid progenitors (red). Error bars are 95% confidence intervals. **f**. Mean expression of selected marker genes in CD14+ cells that are upregulated in monocyte subsets or were differentially expressed in CD14+ cells between ET and PV patients or between MPN patients and healthy controls.

Using this approach, we detected the *JAK2* mutation site (locus encoding codon 617) in at least one transcript in 5-15% of cells (mean 9.5%) in the 7 patient libraries, improving over the existing methods for detecting somatic mutations in single-cell libraries^22,23^. We designated cells in which at least one mutated *JAK2* transcript was detected as *JAK2*-mutant cells. Importantly, cells with a heterozygous *JAK2* mutation express both WT *JAK2* and mutant *JAK2*. We accounted for this when computing the fraction of mutated cells in specific subpopulations. However, for differential expression analysis we conservatively designated all cells in which only WT transcripts were detected as WT (Methods).

Using the expression levels of marker genes, we identified all major hematopoietic lineage progenitors in all samples (Fig. 2a-b) and found that *JAK2-*mutant MPN patients showed a similar hematopoietic differentiation hierarchy as healthy controls (SI Fig. 3). However, we observed that many genes involved in immunity, hematopoiesis, and ribosome biogenesis are differentially expressed between HSCs from healthy controls and both *JAK2* WT and *JAK2*-mutant HSCs from MPN patients, suggesting that the presence of a *JAK2*-mutant clone affects gene expression in both *JAK2* WT and mutant cells (SI Fig. 4). This finding is consistent with the existence of cell-extrinsic factors in the bone marrow of MPN patients that affect WT HSCs as well as mutated HSCs^24,25^.

**Figure 3.**
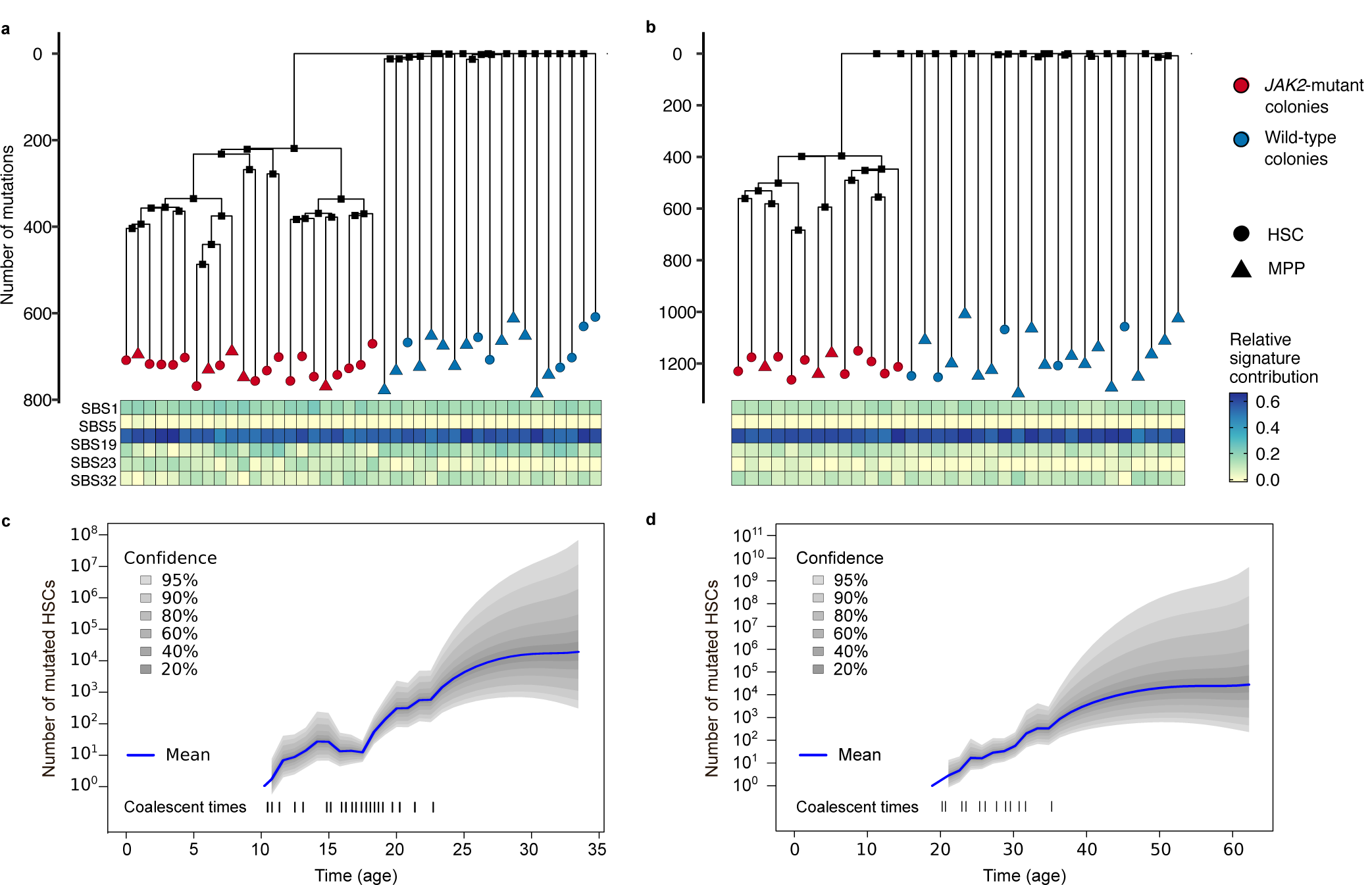
Somatic mutations can be used to reconstruct the lineage trees of wild-type and mutated HSCs. Lineage trees constructed using somatic SNVs for ET 1 (**a**) and ET 2 (**b**). The heatmap below the lineage trees shows the relative contribution of single-base substitution mutational signatures SBS1, SBS2, SBS5, SBS19, SBS23, and SBS32 (see Methods) to the mutational spectrum defined by the private mutations detected in each HSC-derived colony. **c**. Number of mutated stem cells as a function of time inferred from the ET 1 lineage tree assuming one generation per year. The dashed lines on the bottom show the times of the coalescent events in the tree. **d**. Same as c but for the ET 2 patient.

**Figure 4.**
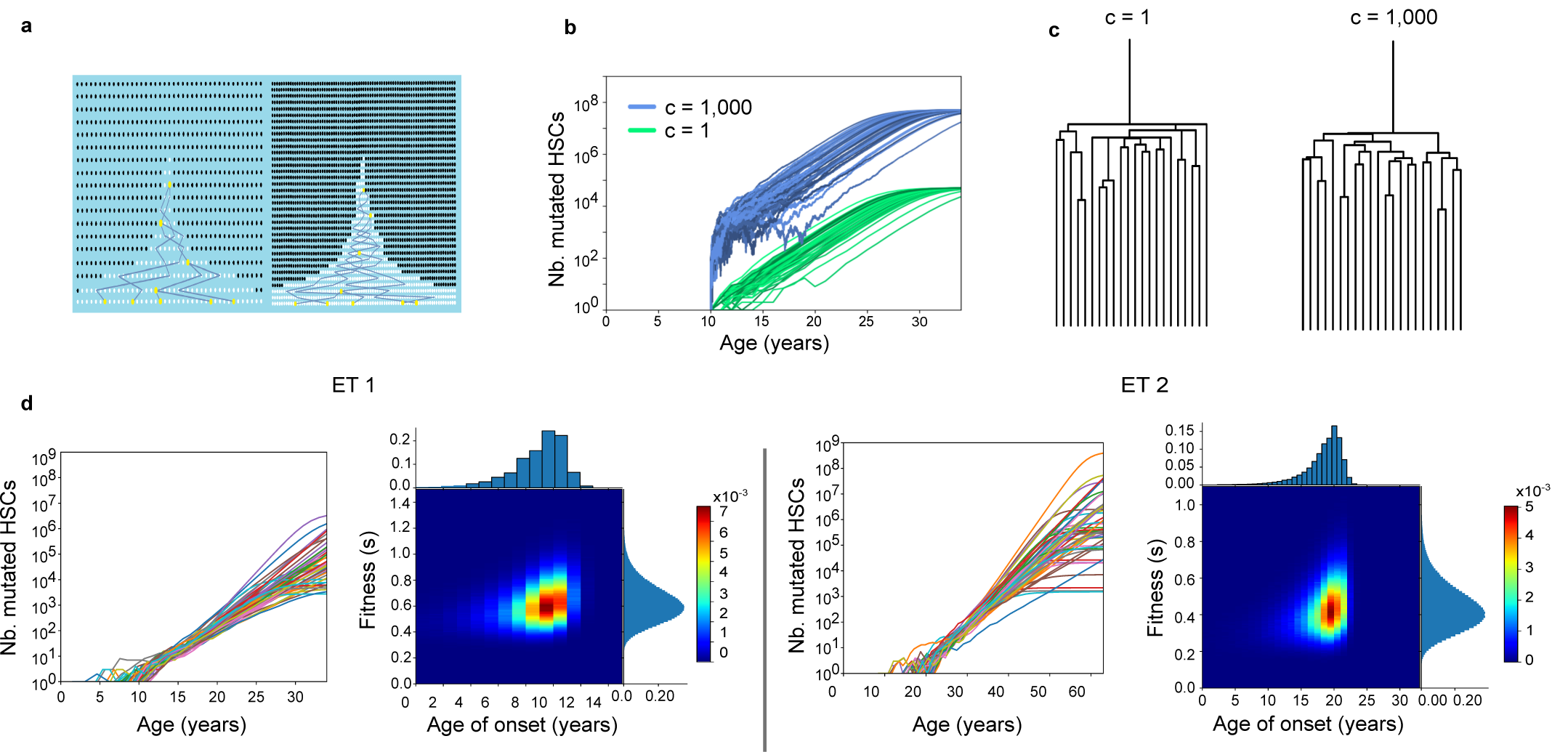
The history of *JAK2-*mutated HSC expansion is reconstructed from the lineage trees. **a**. Schematics show the effect of scaling the number of generations by a factor of two while keeping the onset of the disease and fitness the same. As a result, the number of mutated cells doubles, because early on, the number of mutated cells (shown in white) fluctuates to increase by a factor of two to escape stochastic extinction. Increasing the number of generations increases the coalescent rate. Increasing the number of mutated cells decreases the coalescent rate. Together, these effects cancel and the trees are indistinguishable. **b**. Green curves are 50 simulated mutant HSC trajectories that survived extinction with fitness s = 0.8, and maximum population size of 50,000. Blue curves are similar, except the number of generations and maximum population size are scaled by c = 1,000. This scaling results in 1,000 times as many mutant HSCs through time (blue) because a larger initial population is needed to escape stochastic extinction. **c**. Trees corresponding to the blue and green trajectories in b. are statistically indistinguishable (Methods). **d**. Inference on data from ET 1 and ET 2. For both patients, we show 50 inferred trajectories of the number of mutated stem cells as a function of time. Heatmaps show the inferred joint distribution of the fitness of the cancer cells and the age at which the disease initiating mutation occurred. The marginal distributions are shown as histograms.

*JAK2*-V617F cells had similar gene expression profiles compared with WT cells from the same patient and were found generally intermixed with the WT cells in UMAP visualizations (Fig. 2c). A significant fraction of the HSC subpopulation was mutated in all the patients (ranging from 5% to 62%), consistent with prior observations that the *JAK2*-V617F mutation is detectable in HSPCs^5^. The *JAK2*-V617F mutation frequency was higher in committed megakaryocyte/erythroid lineage progenitors and lower in lymphoid or granulocyte-macrophage (GMP) lineage progenitors (combined *P* < 10^−10^ for both erythroid vs. lymphoid and erythroid vs. GMP for all patients with V617F mutation, Fisher’s exact test with Fisher’s method) (Fig. 2c-d). The fraction of *JAK2*-V617F cells increased along the erythroid differentiation trajectory in both PV and ET patients. In contrast, the *JAK2*-V617L mutation showed no significant lineage bias (Fig. 2c-d, SI Fig. 5), suggesting that it does not have the same effect on HSPC differentiation as the V617F mutation. *TET2* mutations were similarly amplified and identified in the scRNA-seq libraries from patients PV 2 and PV 3, and were present in *JAK2-*mutant cells in both patients (SI Fig. 6). Both patients had a higher *JAK2* allele fraction than *TET2* allele fraction (Fig. 1b; *P* < 10^−10^ for PV 2, *P* = 0.003 for PV 3, Fisher’s exact test), suggesting that the *TET2* mutation occurred in a *JAK2-*mutated cell, as has been previously described^26^.

**Figure 5.**
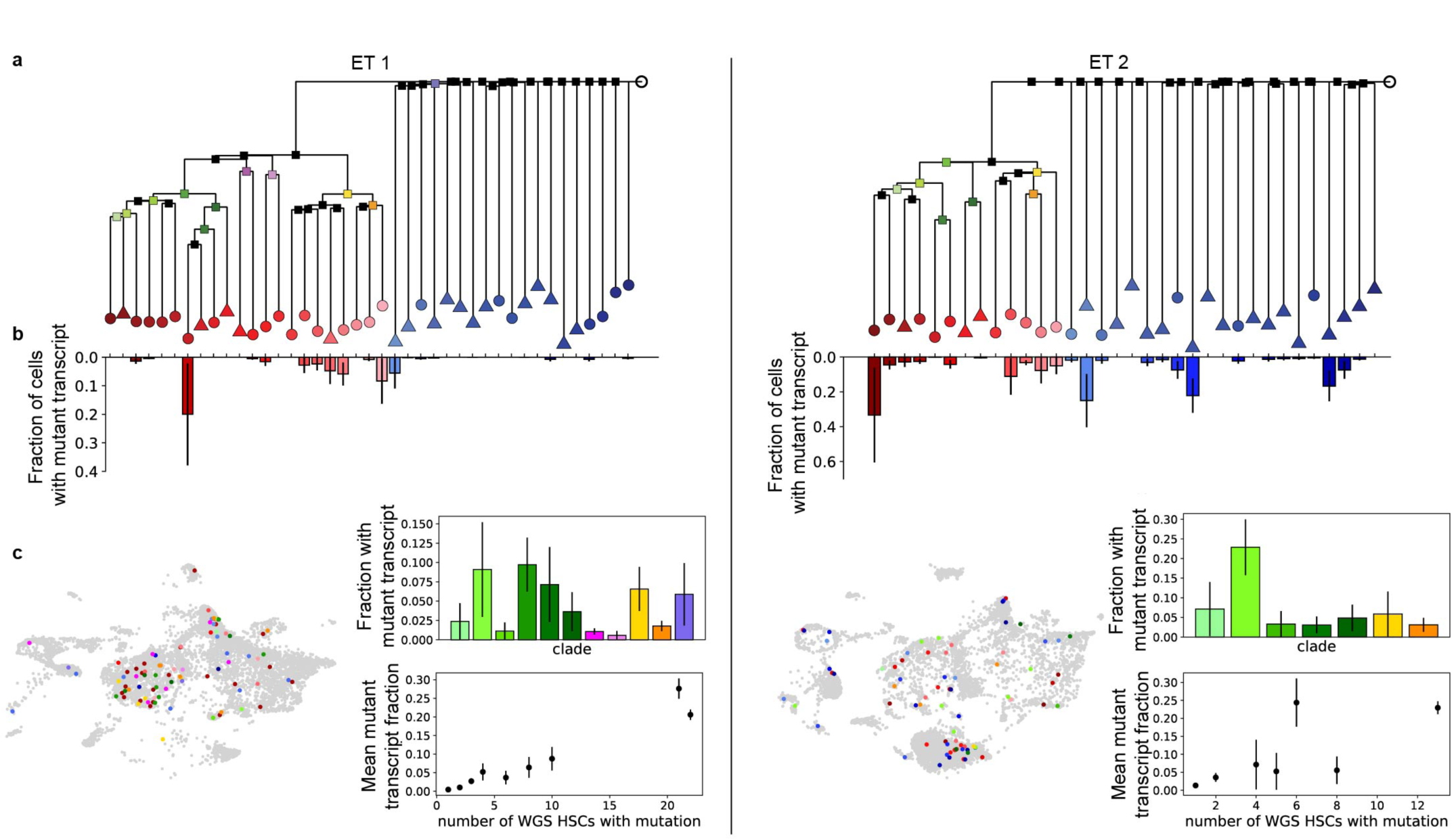
Somatic mutations identify the progeny of individual HSCs in scRNA-seq data. **a**. Lineage trees reconstructed using somatic point mutations detected in the WGS data from single-cell-derived colonies for ET 1 (left) and ET 2 (right). Colored leaves represent the individual HSCs (circles are HSCs and triangles are MPPs). Colored internal nodes represent the HSC clades (a subset of related HSCs) that share a somatic mutation detected in the scRNA-seq data. **b**. Fraction of cells in the single-cell data with a mutant transcript corresponding to a specific HSC out of all cells in which either a WT or mutant version of that HSC-specific transcript was detected. These cells are descendants of the HSC with which they share the somatic mutation. **c**. Left inset: UMAP showing cells with a mutant transcript corresponding to descendants of a single HSC or an HSC clade. Colors correspond to colors of leaves or nodes in a. Right inset: fraction of cells with a mutant transcript corresponding to an HSC clade out of all cells in which either a WT or mutant version of that clade-specific transcript was detected (top). The fraction of mutant transcripts averaged over all mutations shared by HSC clades with the same number of HSCs (bottom). The mutant fraction increases with clade size. All error bars in this figure are +/- one SD.

**Figure 6.**
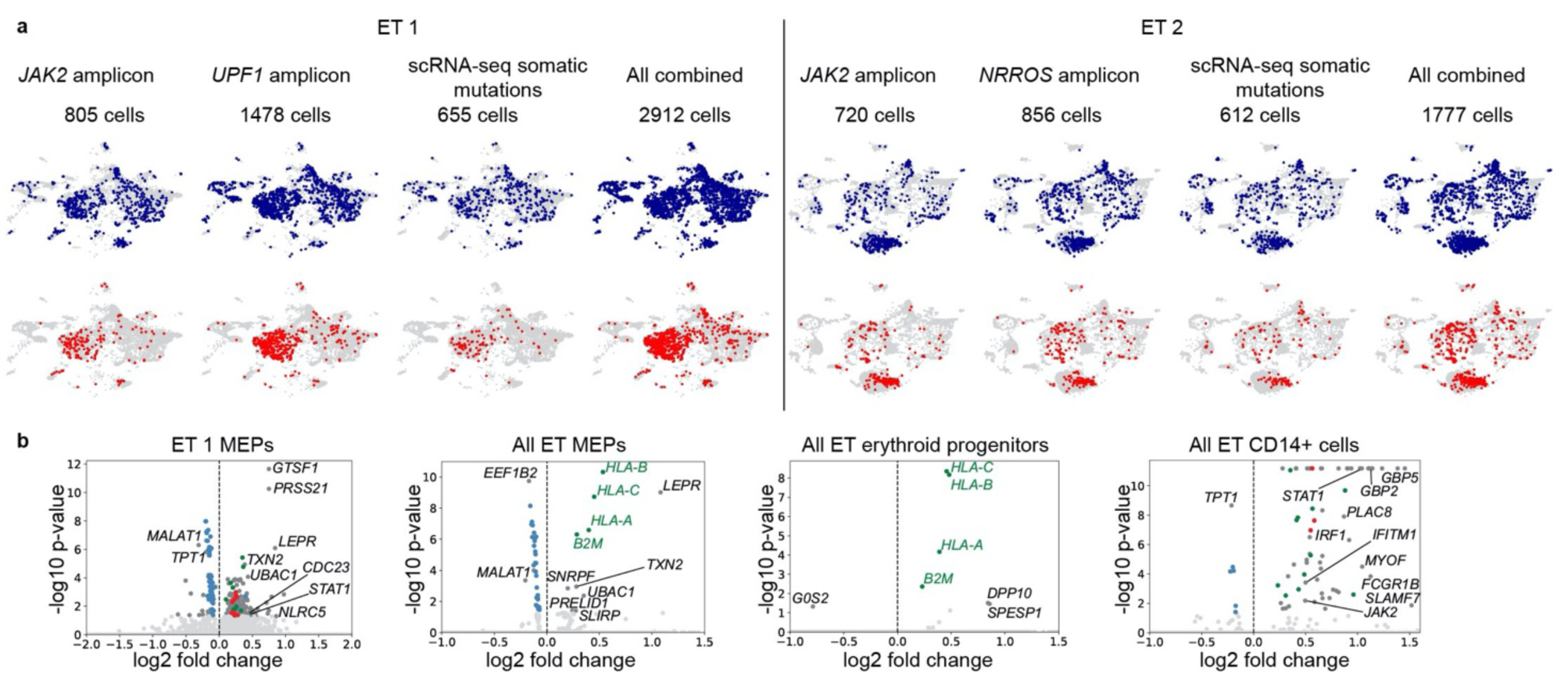
*JAK2*-V617F HSPCs have different gene expression profiles from WT cells. **a**. UMAPs for ET 1 and ET 2 with cells with WT (blue) or mutant (red) transcripts for individual amplicon-sequenced mutations or all somatic mutations called in the scRNA-seq data. Using other somatic mutations in addition to *JAK2* increases the number of cancer cells that are detected. **b**. Differential expression of *JAK2*-V617F mutant vs. WT cells. Blue dots represent ribosomal genes, green dots represent antigen presentation genes, and red dots represent proteasomal genes. Additional selected significant genes are labeled.

In clinical practice, the MPN clone size can be approximated by the peripheral blood *JAK2-*V617F variant allele fraction (VAF), which reflects the fraction of nucleated blood cells that harbor the mutation but does not measure the contribution of the MPN clone to anucleated mature red blood cells and platelets^27^. Our single-cell analysis demonstrates that peripheral blood VAF consistently underestimates the degree to which *JAK2*-mutated cells contribute to steady-state erythropoiesis (Fig. 2e). Indeed, the large fraction of *JAK2-*V617F-mutated erythroid progenitor cells in the PV patients (79% to >95%) suggests that nearly all erythropoiesis arises from the *JAK2-*V617F clone. Furthermore, the peripheral blood VAF does not accurately reflect the extent of disease in HSCs (Fig. 2e).

We identified a CD14+ population in our scRNA-seq data that showed enhanced expression of Type I-II interferon regulated genes in MPN patients relative to healthy controls (Fig. 2f, SI Fig. 7-8). These cells did not express CD34 mRNA and therefore likely represent a contaminating CD34-monocyte-like bone marrow population. Interestingly, the *JAK2*-V617F mutation was highly enriched in CD14+ cells from newly diagnosed patients with ET or PV, suggesting that the *JAK2*-V617F mutation drives the expansion of monocyte-like cells in early-phase MPN patients.

Taken together, our observations show that *JAK2*-V617F HSPCs have gene expression profiles similar to those of WT HSPCs but show a clear bias towards the megakaryocyte-erythroid fate. In addition, a significant fraction of HSCs was mutated in each patient. To understand how this population of mutated stem cells emerged, we set out to determine when the *JAK2*-V617F mutation first occurred in each patient and how the population of mutated stem cells subsequently expanded.

### Lineage trees of individual mutated and WT stem cells

To infer the disease history prior to clinical presentation with MPN, we reconstructed the lineage trees of the *JAK2*-mutated HSCs of two ET patients, ET 1 and ET 2, using the pattern of somatic mutations accrued by individual cells. Somatic mutations occur at random and are passed to a cell’s descendants and thus can be used to establish lineage relationships. To read out the somatic mutations in each cell, we isolated individual HSCs and multi-potent progenitor (MPP) cells using established cell surface markers (Methods) from the CD34+-enriched bone marrow cells of ET 1 and ET 2. We then expanded each HSC or MPP cell *ex vivo* by culturing them for ∼8 weeks and performed whole-genome sequencing (WGS) on the single-cell colonies (SI Fig. 9, Methods). We selected colonies to balance the number of *JAK2*-mutated and WT cells sequenced: 22 *JAK2*-mutated colonies and 20 WT colonies for ET 1; 13 *JAK2*-mutated colonies and 21 WT colonies for ET 2. We observed that *JAK2*-mutated HSCs likely had a proliferative advantage in our culture conditions since the fraction of *JAK2*-mutated HSCs and MPPs after culturing was higher than the fraction of *JAK2*-mutated HSCs found in the scRNA-seq data (9.8% of cultured MPPs and 73% of cultured HSCs vs. 29% of HSCs identified by scRNA-seq for ET 1, see Methods for additional information).

We found that the younger patient (ET 1, 34 years old) had on average 713 ± 45 somatic point mutations in individual HSCs/MPPs, whereas the older patient (ET 2, 63 years old) had 1185 ± 75 mutations in each cell. Using the number of point mutations found in each cell and the age of each patient, we estimated the somatic point mutation rate as 19±1 per year, consistent with previous observations in healthy donors^19,28^ (SI Fig. 10). In both patients, we observed a similar total number of somatic mutations between the *JAK2*-mutated cells and *JAK2*-WT cells (*P* > 0.05, SI Fig. 10 and Methods), indicating that the somatic mutation rate was not altered by the *JAK2*-V617F mutation. The number of somatic mutations was also similar between HSCs and MPPs in both patients (*P* = 0.07 for ET 1, *P* = 0.21 for ET 2; Methods). No somatic structural variants or copy number aberrations were detected, except for the loss of one copy of chromosome X in one colony from patient ET 2 (SI Fig. 11-12).

Analysis of the single-base substitution (SBS) mutation signatures revealed that spontaneous aging-associated clock mutations (COSMIC signatures 1 and 5) predominated in both WT and *JAK2*-V617F HSCs/MPPs^29^ (Fig. 3a-b, SI Figs. 13-16), consistent with previous analyses of somatic mutations in healthy HSPCs^19,28,30^. Other than *JAK2*-V617F, no deleterious somatic mutations were detected in all *JAK2*-mutated cells. The other mutations we identified that occurred in genes that could potentially impact stem cell function, such as *ASH1L* and *FAT1*, were not predicted to affect protein function nor have been previously reported as pathogenic (Methods and SI Fig. 17). Therefore, the *JAK2*-V617F mutation is likely the disease-initiating MPN driver mutation in these two patients.

Next, we used Wagner parsimony to reconstruct the phylogenies of the stem cells from the pattern of somatic mutations (Fig. 3a-b, SI Figs. 18-19, SI Table 2, Methods). Two distinct clades were found in each patient that were defined by the presence or absence of the heterozygous *JAK2*-V617F mutation. These phylogenies suggest that, in both patients, a single *JAK2* mutation event initiated the disease, followed by expansion of the mutated stem cells. No mutations were shared across all *JAK2*-WT stem cells in ET 1 or in ET 2, suggesting that the common ancestor of the *JAK2*-WT stem cells dates back to embryonic development, before most somatic mutations occurred. However, there were many mutations shared across the *JAK2*-mutated stem cells (220 in ET 1 and 398 and ET 2), indicating that all mutated cells descended from a single common ancestor in which the *JAK2* mutation occurred. Using the inferred somatic mutation rate, we estimated that the disease-initiating mutation occurred ∼25 years prior to sampling in ET 1 (34-year-old patient) and ∼40 years prior to sampling in ET 2 (63-year-old patient).

### Reconstructing the history of disease progression in individual patients

To reconstruct the history of disease development, we inferred the number of mutated stem cells in each patient from the time of the initial *JAK2* mutation event to the time of sampling by applying a phylogenetic dynamics inference algorithm^31,19^ to the reconstructed lineage trees. This algorithm assumes that all mutated HSCs in the population are equivalent and that the population size over time can be modelled as a Gaussian process. Additionally, we assumed that HSCs divide symmetrically once per year^16,17,19,28^. We found in both patients that fewer than 100 mutated stem cells were present in the first decade after the *JAK2* mutation occurred. The number of mutated stem cells in both patients grew exponentially for decades (Fig. 3c-d).

To quantitatively estimate the difference in growth rates between *JAK2* WT and *JAK2*-mutant HSCs *in vivo*, we constructed a mathematical model of stem cell self-renewal based on the Wright-Fisher model (Fig. 4a, Methods). Our model contains three parameters: the maximum number of mutated stem cells, the age at which the disease-initiating *JAK2* mutation occurred, and the fitness *s* of the mutated stem cells-corresponding to the proliferative advantage of the mutated stem cells over the wild-type stem cells. If the mutated stem cell population survives stochastic extinction early on when its size is small, initially it will grow exponentially as (1+*s*)^t^ where t is time in years, until the mutated cells take over the majority of the population (Supplemental Text).

Critically, unlike population size^19,32^, fitness *s* can be inferred without any knowledge of the HSC division rate (Fig. 4a-c, SI Fig. 20-21). This is because changing the division rate scales the inferred population size and the minimum population size required to evade stochastic extinction in the same way. Taken together, these two effects cancel each other out and *s* can be inferred directly from the observed lineage trees without knowledge of the division rate.

The model parameters were inferred from the reconstructed single-cell WGS lineage trees using approximate Bayesian computation (ABC). Briefly, we chose the parameter values from a prior distribution, simulated the model, randomly sampled a subset of cells, and obtained their lineage tree (SI Fig. 22). We then compared the simulated tree to the observed tree (as measured by the Lineage-Through-Time metric). If the simulated and observed trees were sufficiently similar, we retained the parameter values. Otherwise, they were discarded, thereby obtaining the posterior distribution for the parameter values (Methods).

We validated this inference procedure using simulated data (SI Figs. 23-25) and then applied it to the observed lineage trees of the *JAK2*-mutated stem cells from ET 1 and ET 2 (Fig. 4d, SI Fig. 26). We inferred that in ET 1, the *JAK2*-V617F mutation first occurred at age 9±2 and had a fitness effect of 63±15%. Similarly, in ET 2, the *JAK2*-V617F mutation first occurred at age 19±3 and had a fitness effect of 44±13%. Taken together, our analyses show that the *JAK2*-mutated HSCs have a selective advantage over the WT HSCs and increase in numbers over the decades before MPN diagnosis.

### Identifying WGS somatic mutations in the scRNA-seq data

To study the hematopoietic output of individual HSCs or HSC clades (subsets of closely related HSCs), we searched for the somatic mutations identified using whole-genome sequencing of individual stem cells in the scRNA-seq data and identified single cells as the progeny of specific HSC lineages or clades (Fig. 5; Methods). To ensure that the somatic mutations identified in the single-cell libraries were not due to sequencing and amplification errors, we constructed a control library comprised of 36 single-cell data sets of bone marrow and peripheral blood mononuclear cells (MNCs) collected from donors other than ET 1 and ET 2 patients and computed the rate of false-positive calls for each somatic mutation site (SI Fig. 27; Methods). Somatic mutations in the single-cell library that occurred at significantly higher frequency than the false positive rate were retained (Methods). Contribution to total hematopoietic output was significantly different between individual HSCs (*P* = 0.015 and 2.99e-10 for ET 1 and ET 2, respectively; *G*-test) and was significantly higher in *JAK2*-V617F HSCs than WT HSCs (*P* = 0.024, 0.001, *G*-test) (Fig. 5b). As expected, the fraction of HSPCs in the scRNA-seq data derived from individual HSC clades increases with clade size (Fig. 5c).

### Transcriptional differences between *JAK2*-V617F and *JAK2*-WT HSPCs

To identify transcriptional changes that could explain the observed effects of the *JAK2*-V617F mutation on the growth and differentiation behavior of mutated HSPCs, we compared the gene expression profiles of *JAK2*-V617F and WT HSPCs. The ability to detect transcriptional changes in the scRNA-seq data depended on the number of sequenced cells for which the *JAK2* mutational status was identified (SI Fig. 28). To increase the power of our differential expression analysis, we used two additional sources of mutational information from the WGS analysis for ET 1 and ET 2 to determine whether individual HSPCs in these patients’ scRNA-seq datasets belonged to the *JAK2*-mutant or *JAK2*-WT clade. First, we selected a subset of somatic mutations in each WGS phylogeny that were common to all sampled *JAK2*-V617F HSCs that define the *JAK2*-mutant clade, and selectively amplified those mutations in the scRNA-seq libraries (Fig. 5d, SI Fig. 28, Methods). Second, we used the transcriptome reads from the scRNA-seq data to detect somatic mutations present in all *JAK2*-mutant HSCs in the WGS data (Fig. 5d, SI Fig. 28, Methods). We found that mutations called from all three sources (*JAK2* amplicon sequencing, amplicon sequencing of other somatic mutations common to all *JAK2*-mutant HSPCs, and somatic mutation calling in the scRNA-seq data) showed a HSPC lineage bias similar to that seen with *JAK2* amplicon sequencing alone. This combined approach increased the fraction of total cells that could be classified into either the *JAK2* mutant or WT clade from 9.8% and 9.8% to 37% and 24% for ET 1 and ET 2, respectively.

After including the expanded set of mutation calls for ET 1 and ET 2, the differential expression analysis comparing *JAK2*-mutant and WT HSPCs in ET patients revealed upregulation of antigen presentation genes, inflammation-related genes, and the leptin receptor in megakaryocyte and erythroid progenitor cells (Fig. 6a-b, SI Fig. 29, SI Tables 3-4). However, we found few significant differences in expression of cell cycle genes between *JAK2*-mutant and WT HSPCs (SI Fig. 30). This suggests that the observed increase in the fraction of more differentiated *JAK2*-mutant megakaryocyte and erythroid progenitors (Fig. 2) cannot be explained by differences in proliferation between *JAK2*-mutant and WT MEPs. Rather, the observed increase is likely due to the mutated HSPCs differentiating more quickly towards the megakaryocyte-erythroid fate (SI Fig. 31, Methods).

Additionally, the normalized expression of ribosomal genes was significantly lower in *JAK2*-mutant (megakaryocyte-erythroid progenitor) MEP cells than WT cells (Fig. 6a-b). This reduction in normalized ribosomal gene expression is consistent with a more differentiated erythroid phenotype, as the relative expression of ribosomal genes decreases along the erythroid differentiation trajectory^33,34^ (SI Fig. 30). While we observed a decrease in the fraction of the transcriptome composed of ribosomal genes during erythroid differentiation, the absolute number of ribosomal gene transcripts per cell increased since the overall number of captured transcripts per cell increased (SI Fig. 30). Interestingly, the total number of transcripts per cell was also higher in *JAK2*-V617F HSCs and MEPs than in WT cells (52% higher in *JAK2*-V617F HSCs, 15% higher in *JAK2*-V617F MEPs; SI Fig. 30). Furthermore, *JAK2*-V617F CD14+ cells had increased expression of inflammation-related genes and proteasomal genes (Fig. 6b). In particular, we found that *SLAMF7*, a cell surface marker previously shown to play a role in fibrocyte differentiation in myelofibrosis, was upregulated in *JAK2*-V617F CD14+ cells^35^. Thus, this population of cells could potentially be an early biomarker for risk of progression to myelofibrosis.

## DISCUSSION

To determine the precise impact of the *JAK2*-V617F mutation on the behavior of human HSCs in their native bone marrow microenvironment, we performed whole-genome sequencing and single-cell profiling on HSPCs from MPN patients. Although it has previously been shown that the *JAK2*-V617F mutation is detectable in HSPC in MPN^5^, we demonstrate that the mutation arises in a single HSC decades before the MPN diagnosis. Subsequently, the population of *JAK2*-mutated stem cells grows exponentially but may exhibit large fluctuations and even stochastic extinction when its size is small in the first few years after the occurrence of the mutation.

Although *JAK2*-V617F clonal hematopoiesis has been described^36–38^ and has been noted to be more common than *JAK2*-V617F+ MPN^39^, to our knowledge, our study is the first to actually measure in an individual patient with MPN, the time interval between *JAK2*-V617F acquisition and MPN development. Our findings that the *JAK2*-V617F mutation occurred in the first decade of life (9±2 years) in a man who developed ET at age 34 and in the second decade of life (19±3 years) in a woman who developed ET at age 63 are striking both in terms of the young age at the time of *JAK2*-V617F acquisition and the decades-long interval to MPN development in both cases.

We found that at the time of MPN diagnosis, a significant fraction of HSCs (5% or more), are descendants of the *JAK2*-mutated HSC. Using the structure of the HSC lineage tree, we inferred the fitness advantage of *JAK2*-mutated HSCs in 2 MPN patients during the pre-diagnosis period to be approximately 40-65%. Our inferred fitness advantage is larger than that found in a population-level study of CHIP, which analyzed peripheral blood variant allele fractions in large cohorts of individuals^40^. This discrepancy suggests that the development of full-blown MPN may require a faster-growing *JAK2*-mutant clone than that observed in clonal hematopoiesis.

In addition to modifying HSC proliferation dynamics, the *JAK2*-V617F mutation also impacts the differentiation dynamics of their progenies. We found that the hematopoietic differentiation hierarchy is largely preserved in ET and PV patients, in contrast to recent observations of aberrant megakaryopoiesis in patients with MF^23^. However, *JAK2*-mutated HSPCs showed a lineage bias towards the megakaryocyte-erythroid fate and strikingly, we found that the fraction of *JAK2*-mutated cells varied significantly across different progenitor cell populations, suggesting that peripheral blood monitoring may not accurately assess *JAK2* mutation burden. Surprisingly, almost all of the erythroid progenitors, even in ET patients, descended from *JAK2*-mutant HSCs.

Several of the novel biological insights we have uncovered have important clinical implications. Although earlier studies using next generation sequencing in MPN patients^41,42^ and mouse models^43,44^ have suggested that *JAK2*-V617F alone is sufficient to engender MPN, our work tracing the pathogenesis of MPN in individual patients to the acquisition of the *JAK2*-V617F mutation in a single HSC, provides a compelling rationale to develop safe and effective treatments to selectively target *JAK2*-V617F e.g. using *JAK2*-V617F mutant-specific inhibitors. This rationale is further supported by the fact that *JAK2*-V617F clonal hematopoiesis is associated with an increased risk of cardiovascular disease^36^ and venous thrombosis^45,46^ and by the decades long time interval between *JAK2*-V617F acquisition and MPN development uncovered by our work. Given that the *JAK2-*V617F mutation has been shown to have cell-intrinsic effects not only in leukocytes^46,47^, but also in erythroid cells^48,49^ and in platelets^50,51^, the finding of a high *JAK2*-V617F allele fraction in megakaryocyte-erythroid lineage cells in patients with low peripheral blood *JAK2-V617F* mutational burden may help explain the development of thrombosis in these patients^52^.

Many cancers start when a genetic alteration arises in a single cell and confers a fitness advantage over the other cells. By the time the disease manifests clinically, this cell has expanded to millions of cells or more. Naturally occurring somatic mutations provide a glimpse into the history of cancer in each patient, revealing when the driver mutations first occurred, how the population of cancer cells expanded, and how their proliferation and differentiation dynamics differ from healthy cells. The framework that we have developed to harness somatic mutations as a clock to reconstruct the lineage tree of cancer cells, follow the differentiation trajectories of their progenies, and assess the transcriptional consequences of such mutations, is broadly applicable to solid tumors and other hematopoietic malignancies.

## ACKNOWLEDGEMENTS

We thank the patients and their families for their participation in our study. S.H. acknowledges funding from NIH NIGMS R00GM118910, DFCI BCB Fund Award, Jayne Koskinas Ted Giovanis Foundation, The William F. Milton Fund at Harvard University, AACR-MPM Oncology Charitable Foundation Transformative Cancer Research Grant, and Gabrielle’s Angel Foundation for Cancer Research. S.H. and A.M. acknowledge funding from Claudia Adams Barr Program in Cancer Research. A.M. acknowledges funding from NIH NHLBI (R01HL131835). A.M. is a Scholar of The Leukemia & Lymphoma Society. C.R.R acknowledges funding from NIH NHLBI T32HL116324. D.V.E. acknowledges funding from the NSF-Simons Center for Mathematical and Statistical Analysis of Biology at Harvard, award number #1764269, and the Harvard Quantitative Biology Initiative. D.V.E and F.M. acknowledge support by the Ludwig Center at Harvard and the Dana-Farber Cancer Institute’s Physical Science-Oncology Center (NIH U54CA193461, to F.M). I.C.-C. and M.K. acknowledge funding from EMBL. J.E. acknowledges funding from NIH NIGMS R25GM109436. Portions of this research were conducted on the O2 High Performance Compute Cluster, supported by the Research Computing Group, at Harvard Medical School. See http://rc.hms.harvard.edu for more information. We thank Drs. David Weinstock and Julie-Aurore Losman for valuable feedback on the manuscript.

## AUTHOR CONTRIBUTIONS

R.M.S., A.M., and S.H. conceived the project. M.N., S.L., S.H. designed the experiments. M.N. devised and optimized single-cell amplicon sequencing protocol with help from S.L. and supervision from S.H. M.N. and S.L. processed the patient samples and generated all the sequencing libraries with help from B.K. C.R.R., G.S.H., and A.M. devised the patient selection criteria. D.J.D., I.G., M.W., E.S.W., M.R.L., R.S., J.S.G., G.H., and A.M. helped obtain patient samples coordinated by C.R.R. D.V.E analyzed all the single cell data with help from C.R.R. and B.K. supervised by F.M. and S.H. M.K. and I.C.-C. analyzed the whole-genome sequencing data and reconstructed the lineage trees. S.P. isolated and cultured individual HSCs supervised by F.D.C. D.V.E. and I.C.-C. devised and implemented algorithm for identifying somatic mutations in single-cell libraries. J.E. devised and implemented the algorithms for inference of growth dynamics from lineage trees supervised by S.H. D.V.E., J.E., M.N., S.L., C.R.R., F.M., A.M., I.C.-C., and S.H. wrote the manuscript with input from all authors. F.M., A.M., I.C.-C. and S.H. supervised the project.

