## Supplemental Figures for "Reconstructing the lineage histories and differentiation trajectories of individual cancer cells in *JAK2*-mutant myeloproliferative neoplasms"

### Supplemental Figure 1

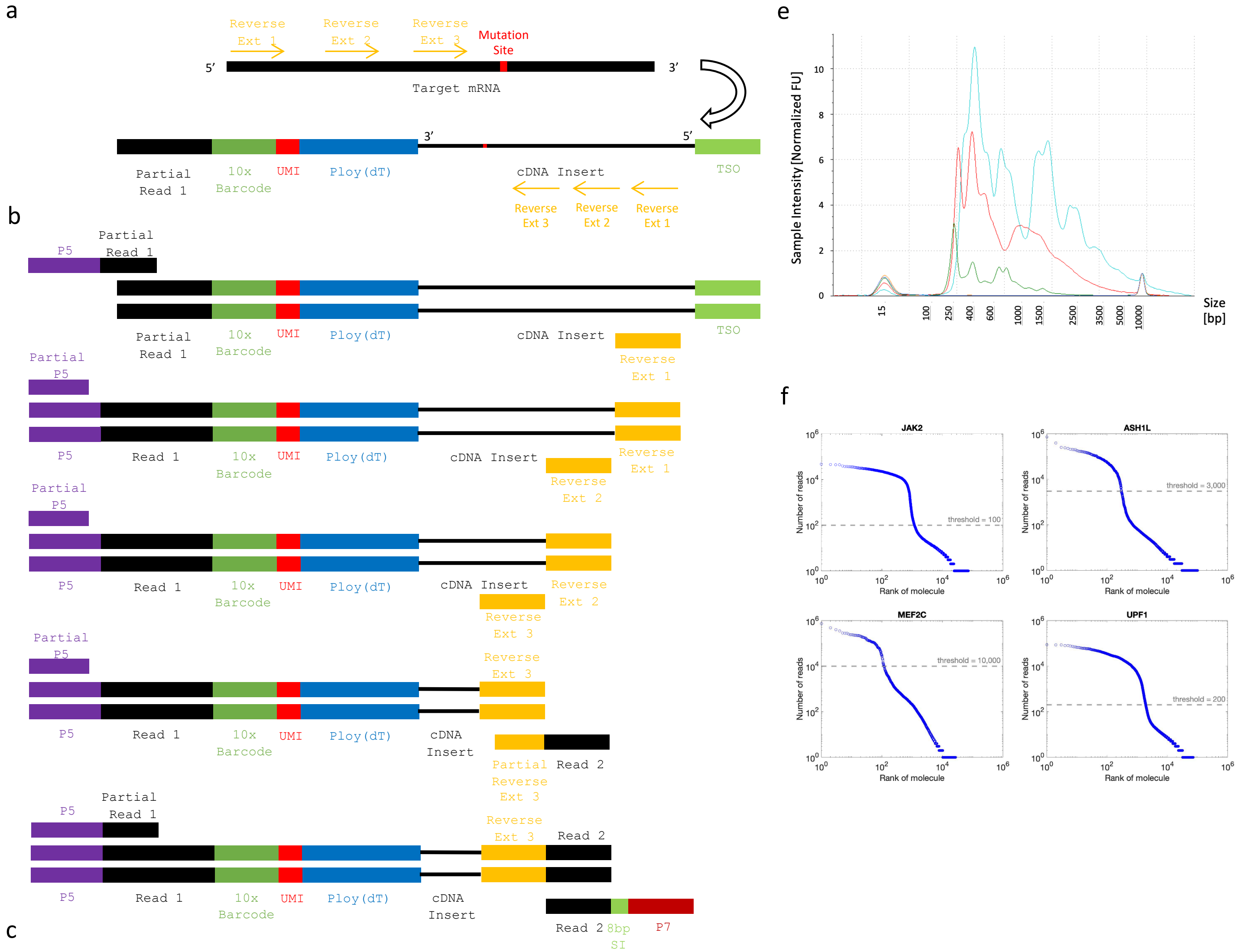


c

5'- AATGATACGGCGACCACCGAGATCTACAC-TCTTTCCCTACACGACGCTC ------>
5'- CTACACGACGCTCTTCCGATCT-NNNNNNNNNNNNNNNN-NNNNNNNNNNNN-(dT)-cDNA_Insert-CCCATGTACTCTGCGTTGATACCACTGCTT -3'
3'- GATGTGCTGCGAGAAGGCTAGA-NNNNNNNNNNNNNNNN-NNNNNNNNNNNN-(pA)-cDNA_Insert-GGGTACATGAGACGCAACTATGGTGACGAA -5'
<------ Reverse_Ext_1 -5'

5'- AATGATACGGCGACCACCGAGATCT ------>
5'- AATGATACGGCGACCACCGAGATCTACAC-TCTTTCCCTACACGACGCTCTTCCGATCT-NNNNNNNNNNNNNNNN-NNNNNNNNNNNN-(dT)-cDNA_Insert_Reverse_Ext_1 -3'
3'- TTACTATGCCGCTGGTGGCTCTAGATGTG-AGAAAGGGATGTGCTGCGAGAAGGCTAGA-NNNNNNNNNNNNNNNN-NNNNNNNNNNNN-(pA)-cDNA_Insert_Reverse_Ext_1 -5'
<------ Reverse_Ext_2 -5'

5'- AATGATACGGCGACCACCGAGATCT ------>
5'- AATGATACGGCGACCACCGAGATCTACAC-TCTTTCCCTACACGACGCTCTTCCGATCT-NNNNNNNNNNNNNNNN-NNNNNNNNNNNN-(dT)-cDNA_Insert_Reverse_Ext_2 -3'
3'- TTACTATGCCGCTGGTGGCTCTAGATGTG-AGAAAGGGATGTGCTGCGAGAAGGCTAGA-NNNNNNNNNNNNNNNN-NNNNNNNNNNNN-(pA)-cDNA_Insert_Reverse_Ext_2 -5'
<------ Reverse_Ext_3 -5'

5'- AATGATACGGCGACCACCGAGATCT ------>
5'- AATGATACGGCGACCACCGAGATCTACAC-TCTTTCCCTACACGACGCTCTTCCGATCT-NN...NN-NNNNNNNNNNNN-(dT)-cDNA_Insert_Reverse_Ext_3 -3'
3'- TTACTATGCCGCTGGTGGCTCTAGATGTG-AGAAAGGGATGTGCTGCGAGAAGGCTAGA-NN...NN-NNNNNNNNNNNN-(pA)-cDNA_Insert_Reverse_Ext_3 -5'
<------Partial_Reverse_Ext_3-TCTAGCCTTCTCGTGTGCAGACTTGAGGTCAGTG -5'

5'- AATGATACGGCGACCACCGAGATCT ------>
5'- AATGATACGGCGACCACCGAGATCTACAC-TCTTTCCCTACACGACGCTCTTCCGATCT-NN...NN-NN...NN-(dT)-cDNA_Insert_Reverse_Ext_3-AGATCGGAAGAGCACACGTCTGAACTCCAGTCAC -3'
3'- TTACTATGCCGCTGGTGGCTCTAGATGTG-AGAAAGGGATGTGCTGCGAGAAGGCTAGA-NN...NN-NN...NN-(pA)-cDNA_Insert_Reverse_Ext_3-TCTAGCCTTCTCGTGTGCAGACTTGAGGTCAGTG -5'
<------ TCTAGCCTTCTCGTGTGCAGACTTGAGGTCAGTG-NNNNNNNN-
TAGAGCATACGGCAGAAGACGAAC -5'

d

JAK2_V617F (G→T)

JAK2_EXT_1
5'-...GTGTTTCTGATGTACCAACCTCACCAACATTACAGAGGCCTACTCATATGAACCAAATGGTGTTTCACAAAATCAGAAATGAAGATTTGATATTTAATGAAAGCCTTGGCCAAGGCACTTTTACAAAGATTTTTAAAGGCGTACGAAGAGAAGT
JAK2_EXT_2 JAK2_EXT_3
AGGAGACTACGGTCAACTGCATGAAACAGAAGTTCTTTTAAAAGTTCTGGATAAAGCACACAGAAACTATTCAGAGTCTTTCTTTGAAGCAGCAAGTATGATGAGCAAGCTTTCTCACAAGCATTTGGTTTTAAATTATGGAGTATGTGTCTGTGG...-3'

UPF1(G→T)

UPF1_EXT_1 UPF1_EXT_2
5'-... ATTTATTCCCATTGCTCTAGGGCTTTCGGTTTCCCCTTCTTCCGGTAGGCCGCGTAGAGGCATGCACCGGGTAGGTTTCCGCGGTGACCCCGCGGCGGCCTGAGGGACGCTCCCTGCCCCATCCCGGCTGTTGGGCTGGGCCGCTTTGCCTCTGCTTC
UPF1_EXT_1
GCCCTGTGCTGTGTTCTCCAGCTTTGTAGCAGCAGCCTTGACAAACCCAGGCGCA...-3'

### Supplemental Figure 2

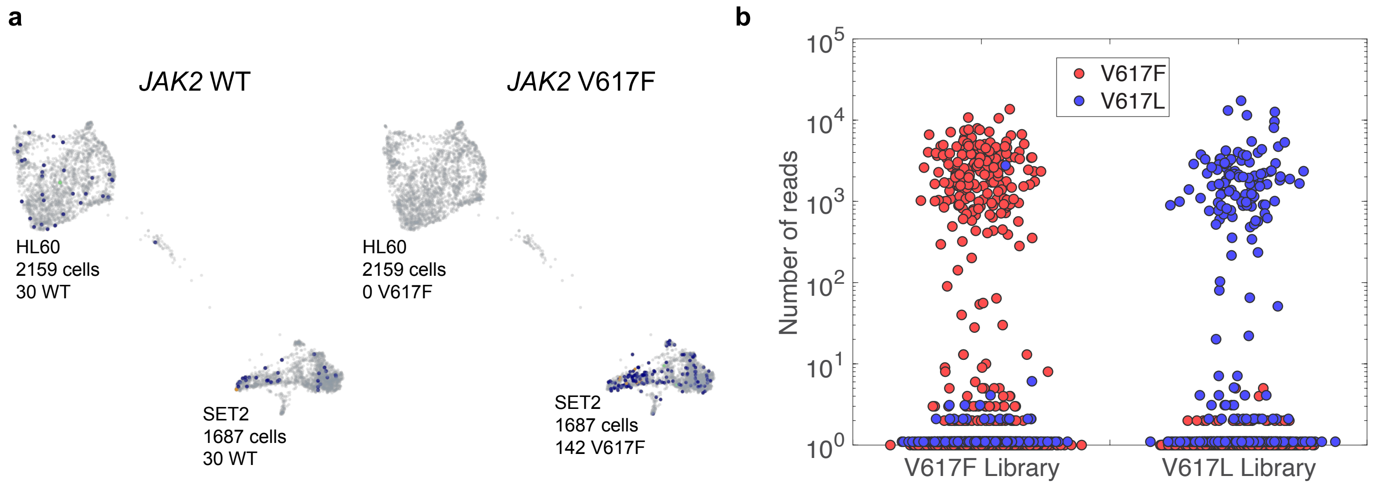

Supplemental Figure 3

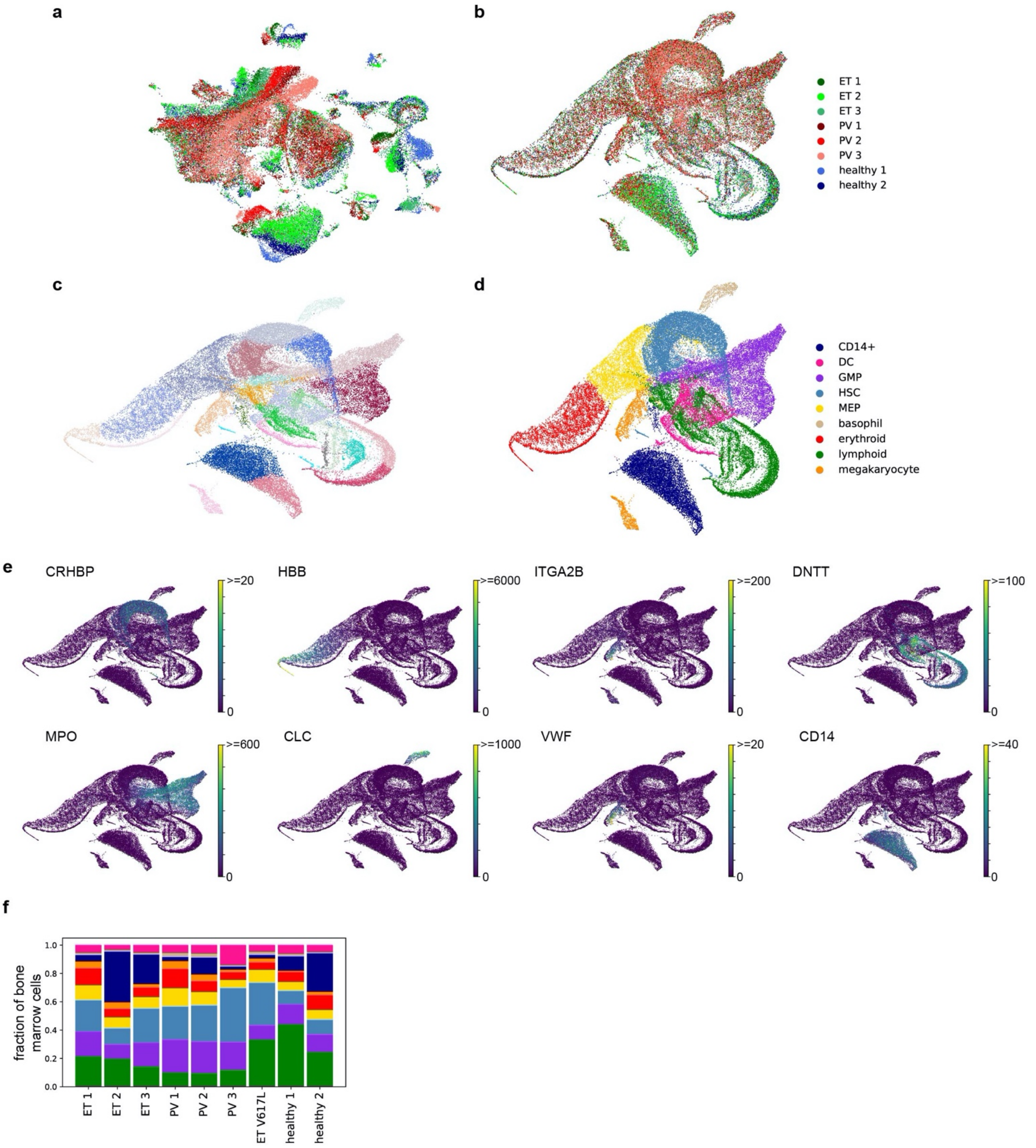

Supplemental Figure 4

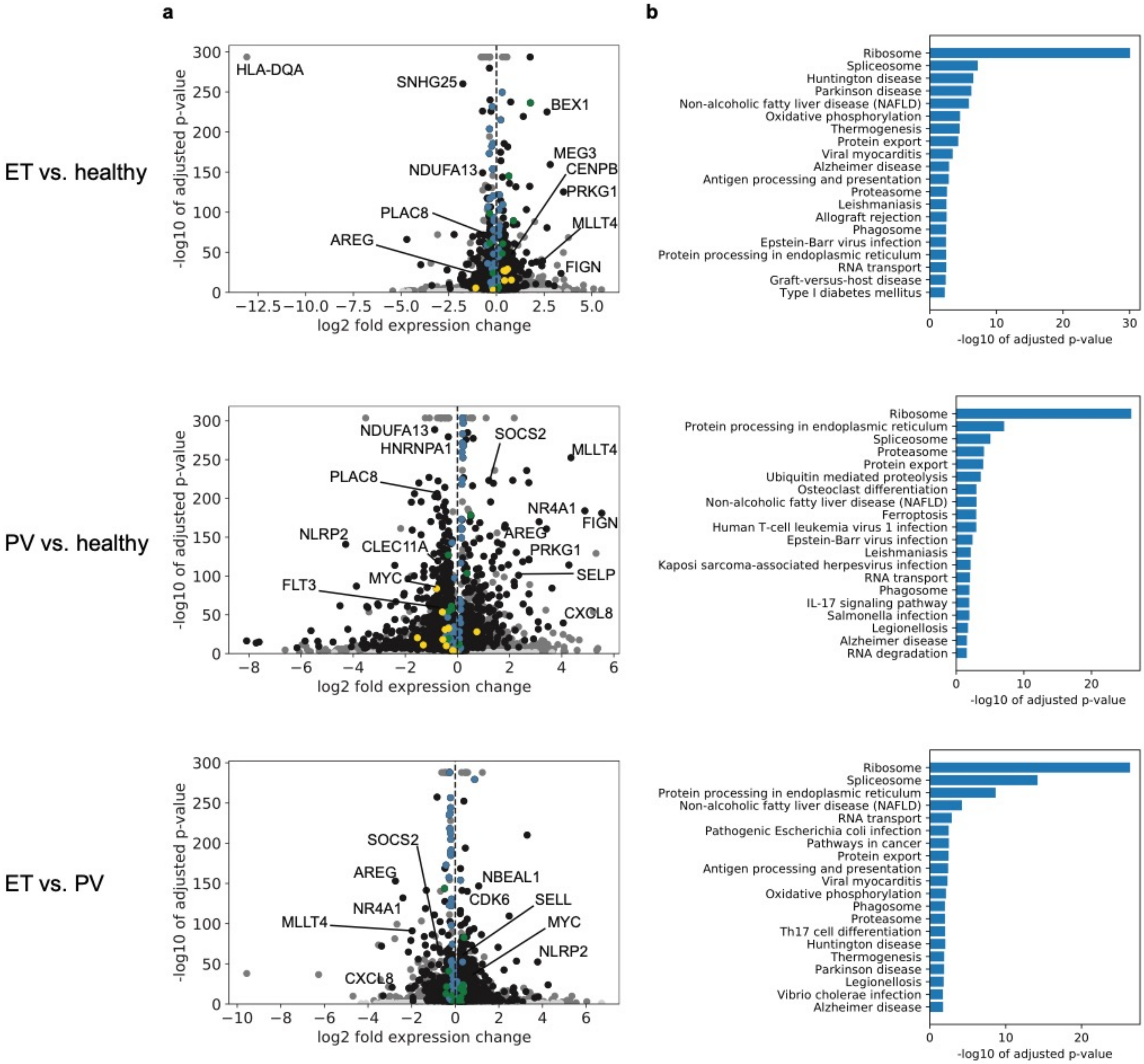

Supplemental Figure 5

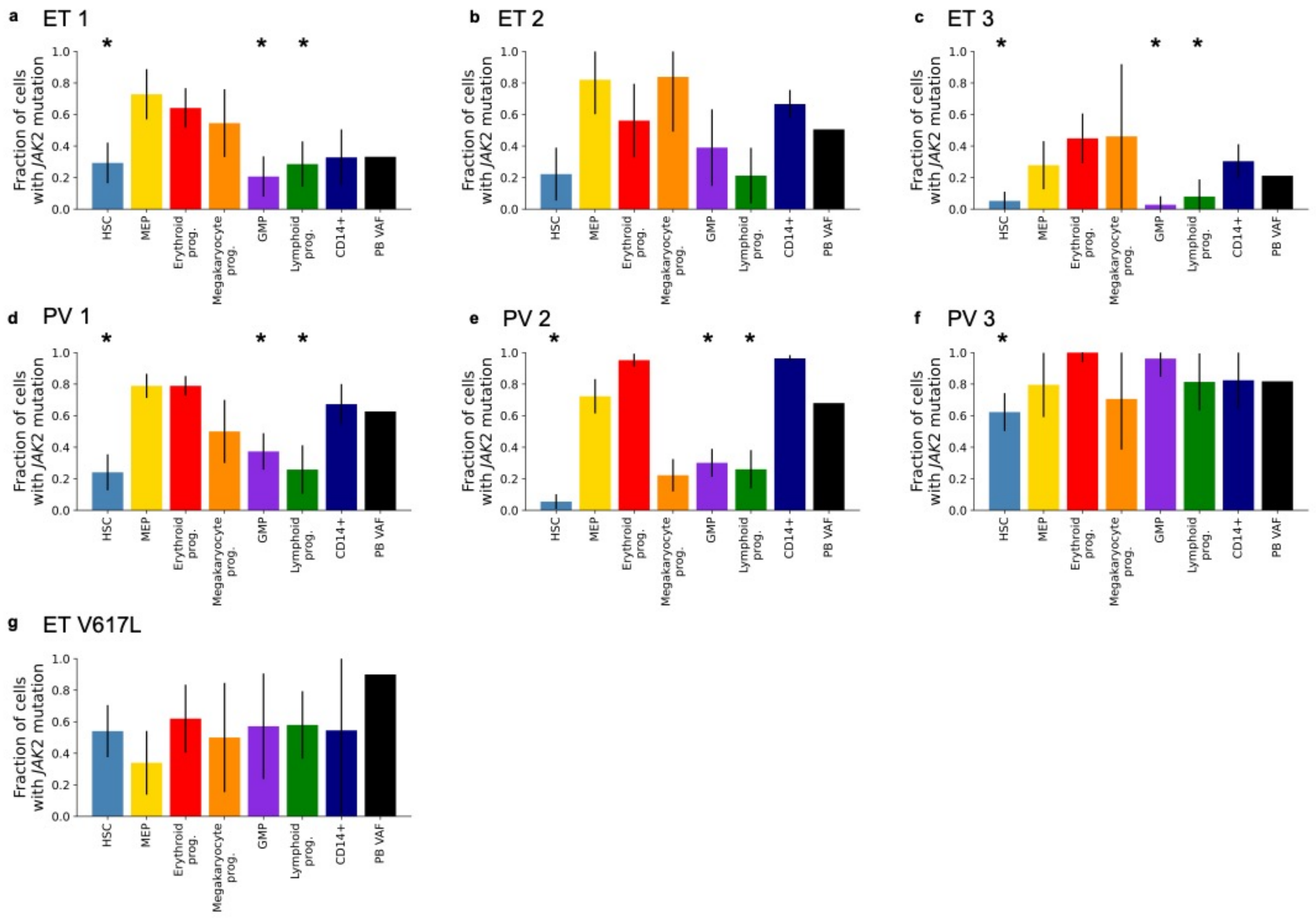

Supplemental Figure 6

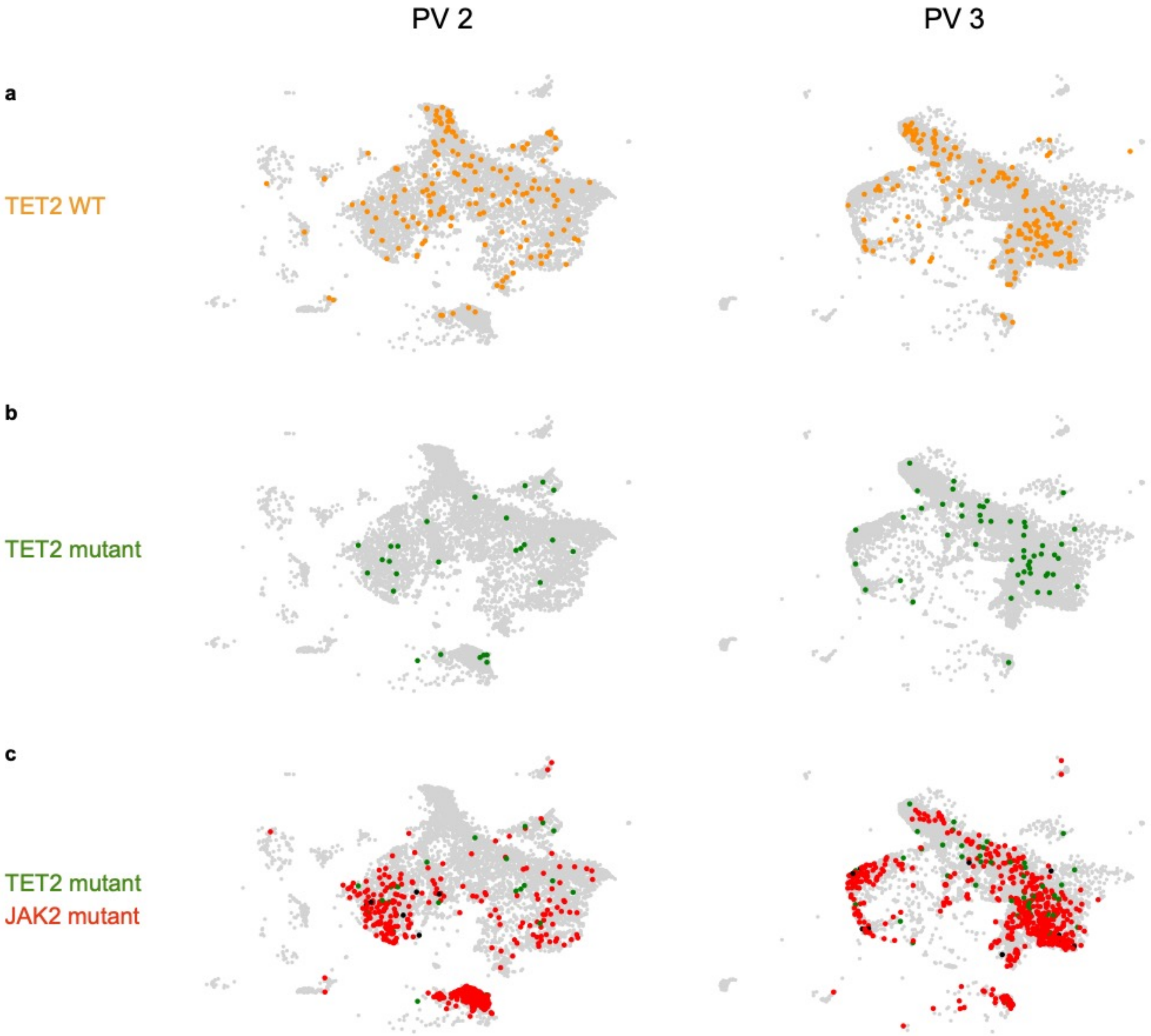


d

| PV 2 | TET2 WT | TET2 mut |
|---|---|---|
| JAK2 WT | 9 | 0 |
| JAK2 mut | 10 | 6 |

| PV 3 | TET2 WT | TET2 mut |
|---|---|---|
| JAK2 WT | 34 | 8 |
| JAK2 mut | 13 | 10 |

Supplemental Figure 7

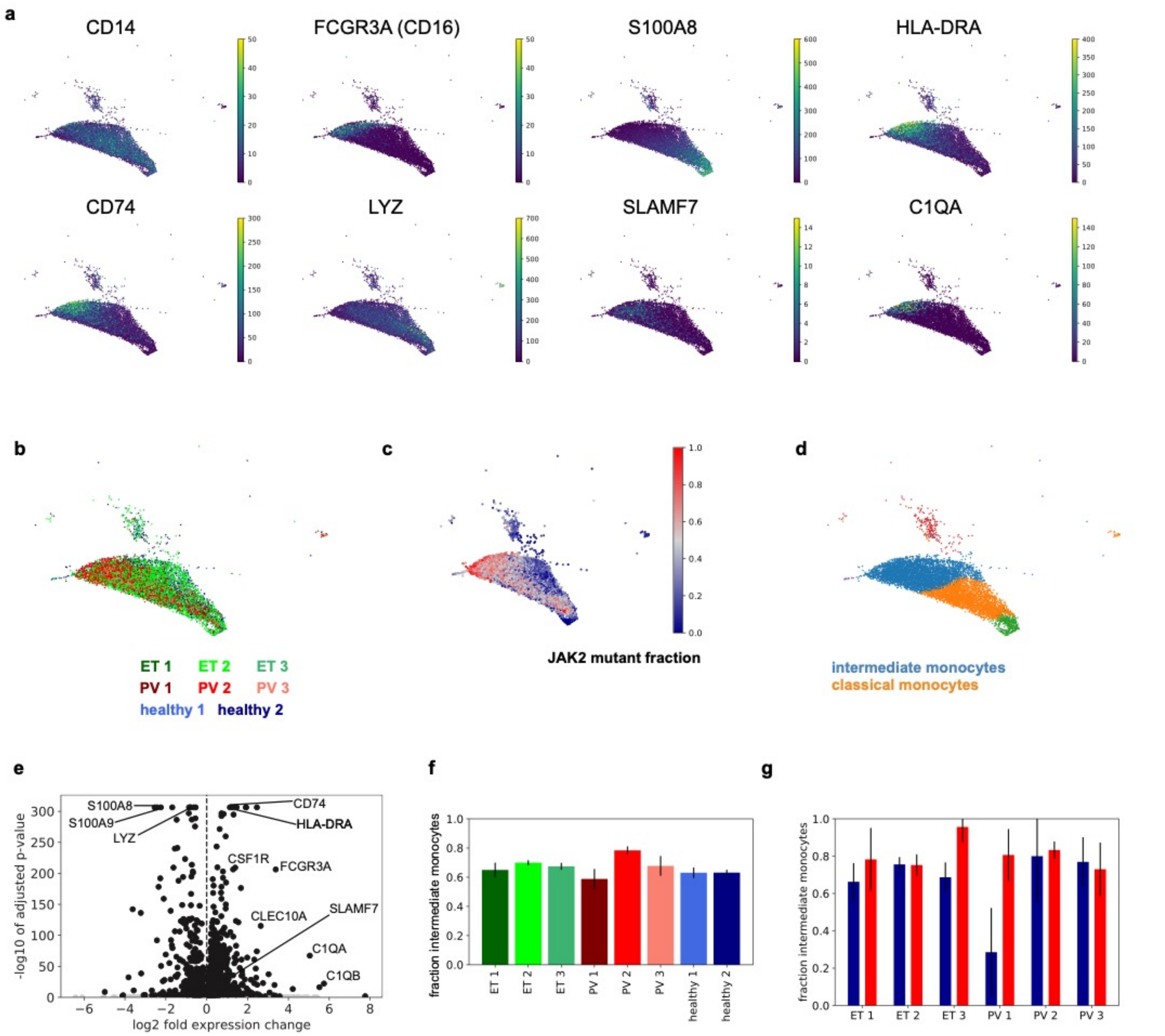

Supplemental Figure 8

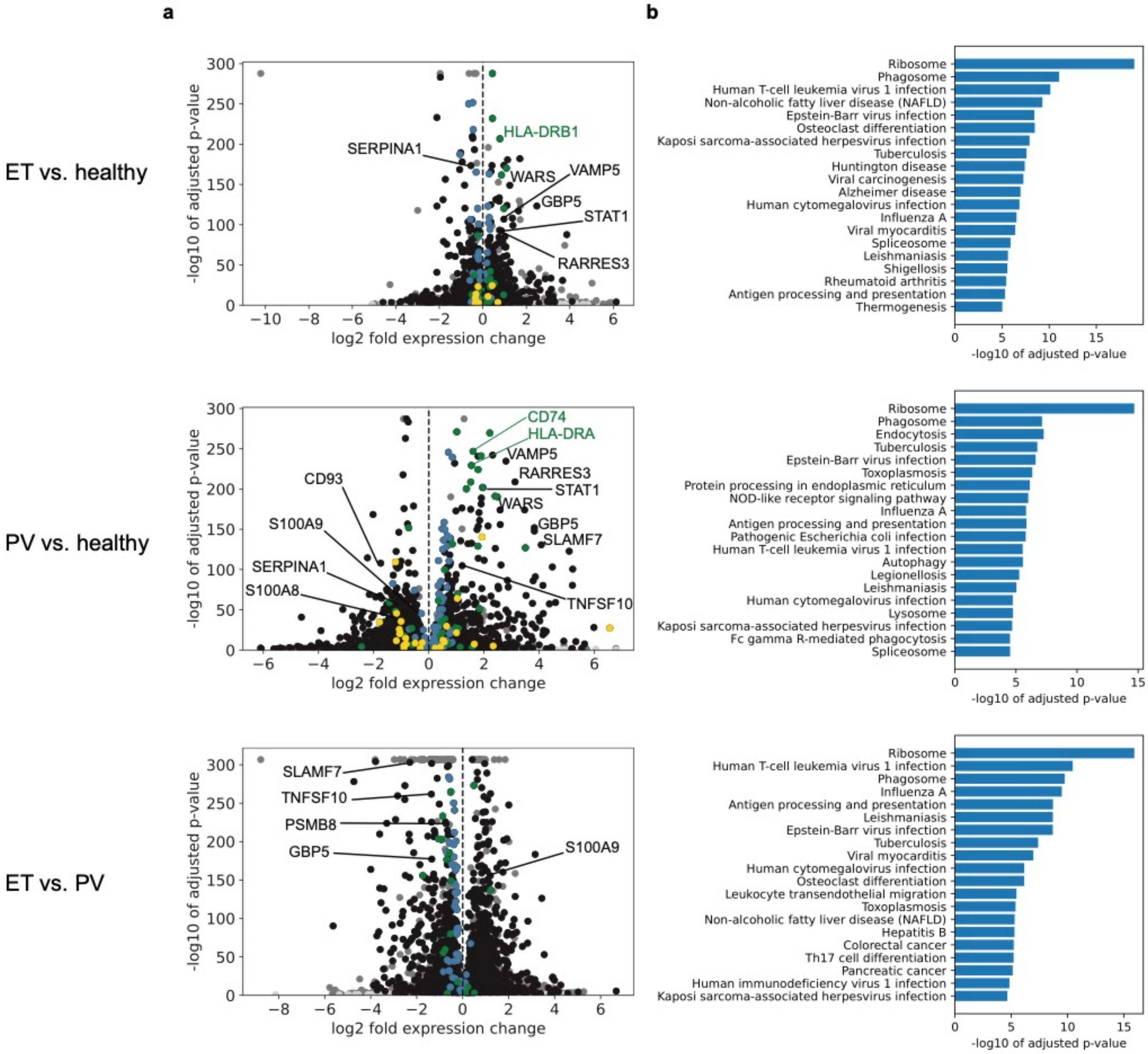

Supplemental Figure 9

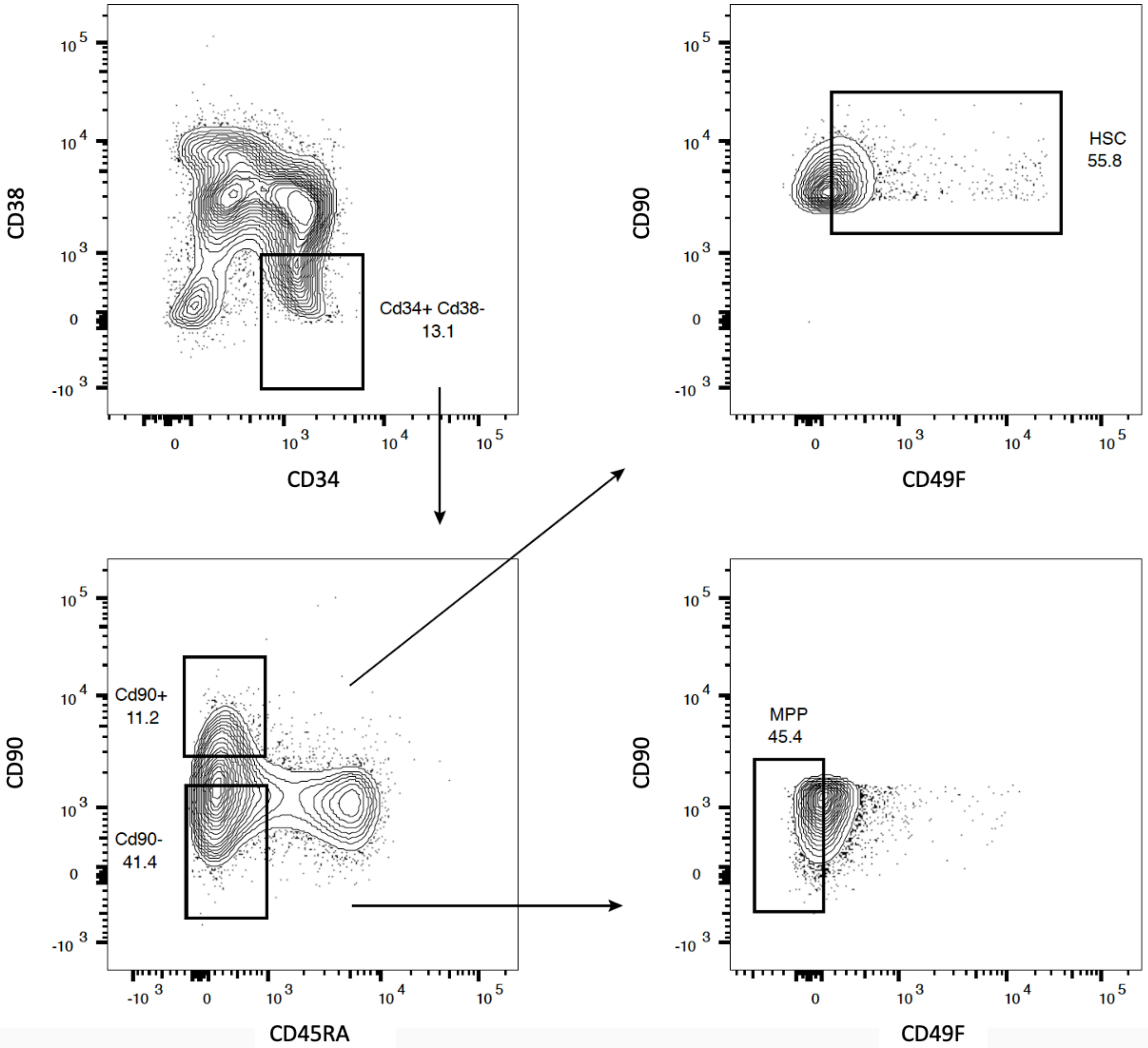

Supplemental Figure 10

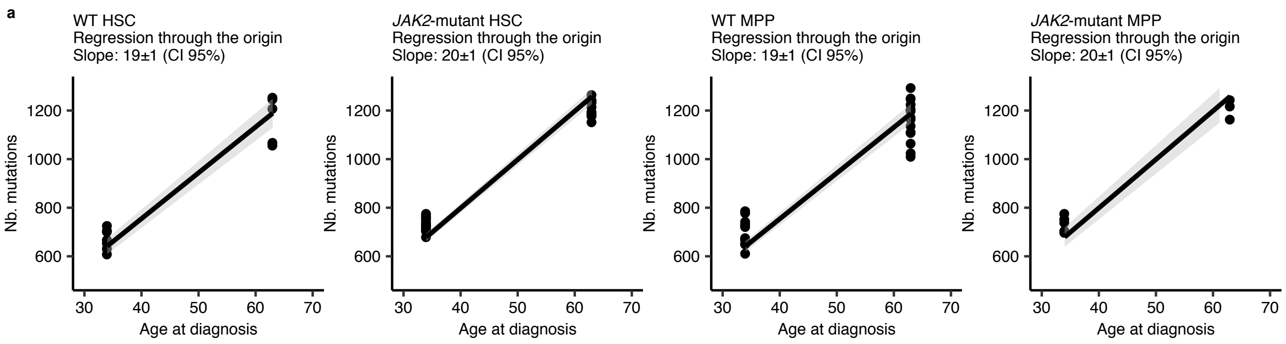

Supplemental Figure 11

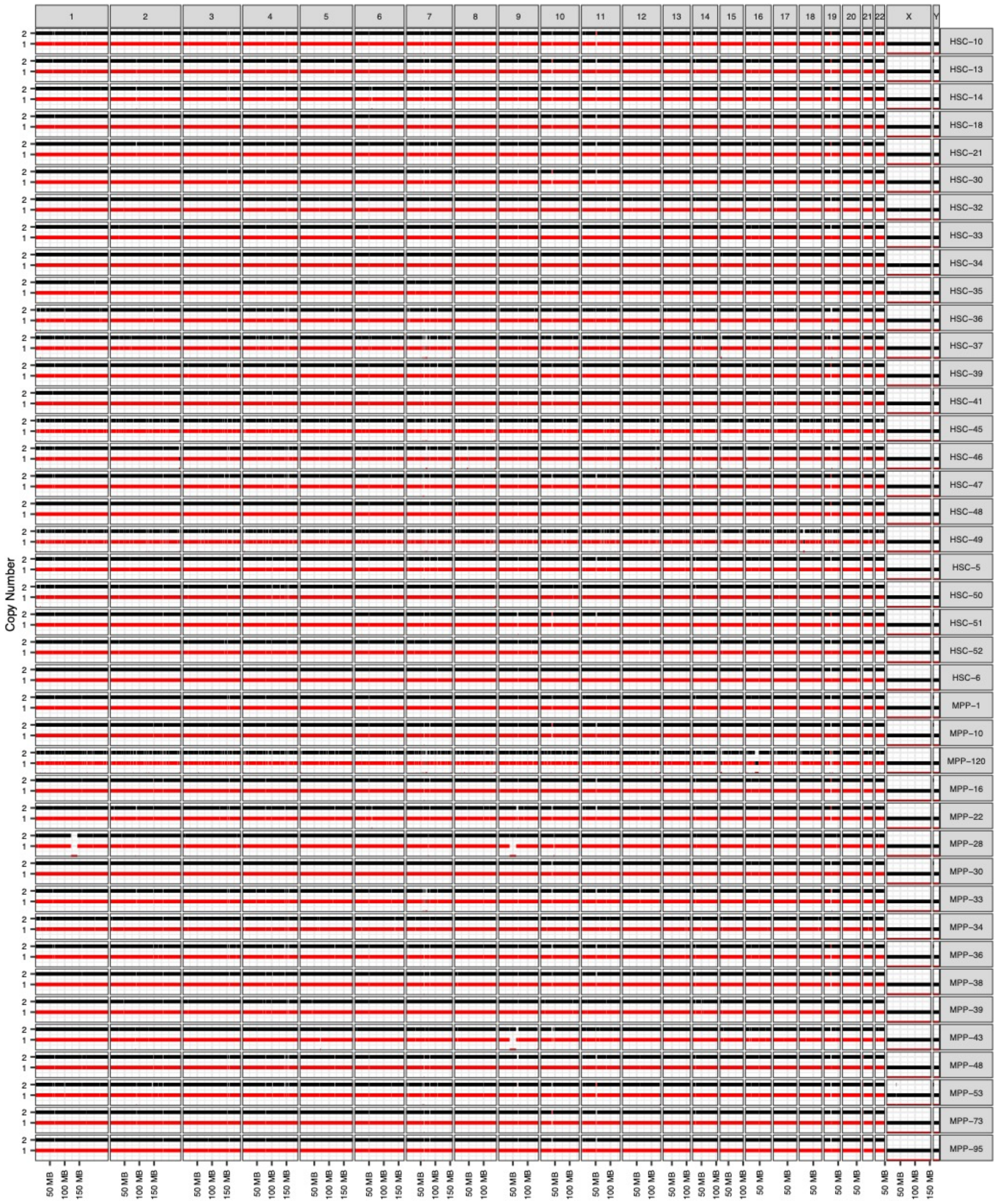

Supplemental Figure 12

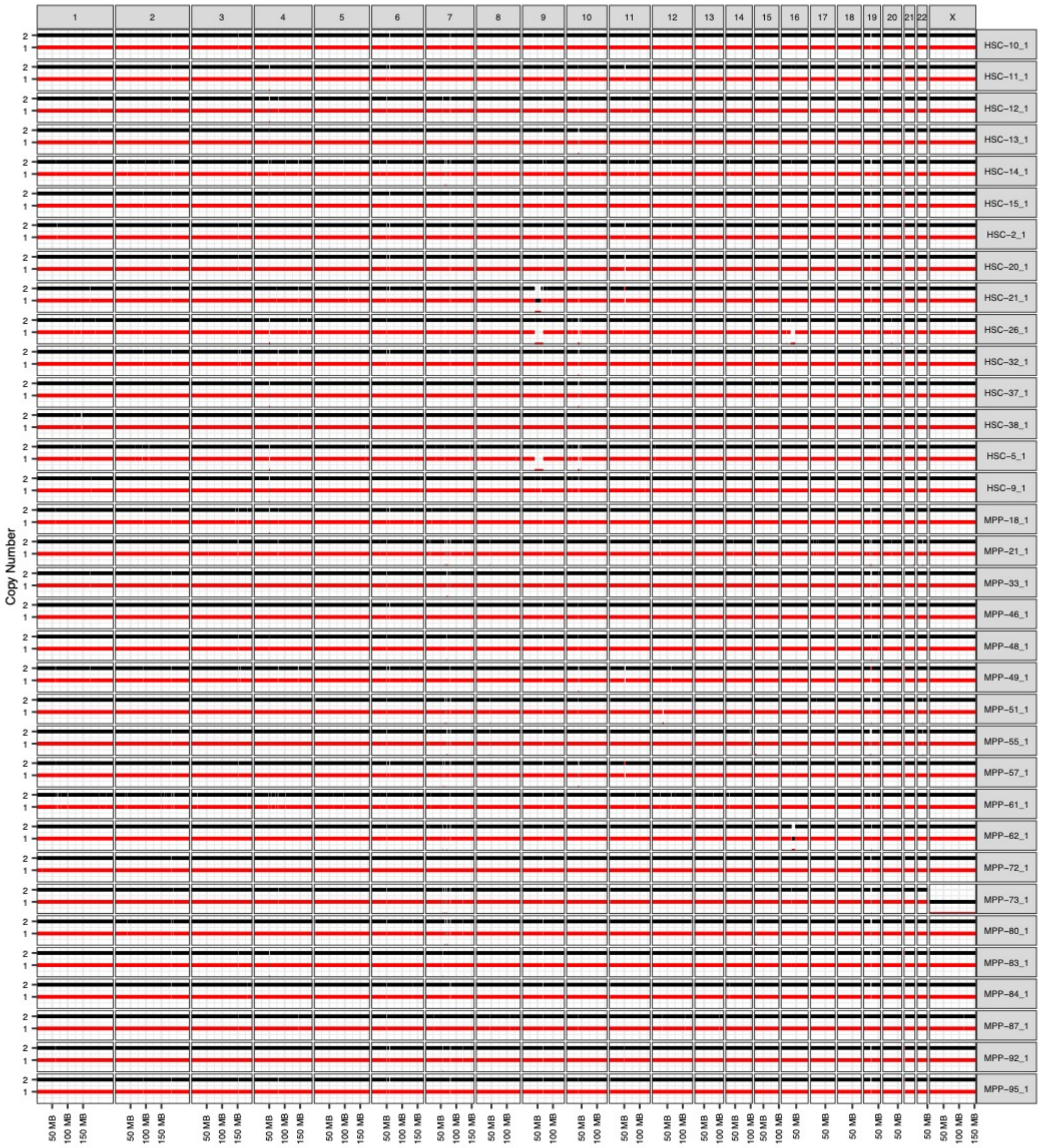

Supplemental Figure 13

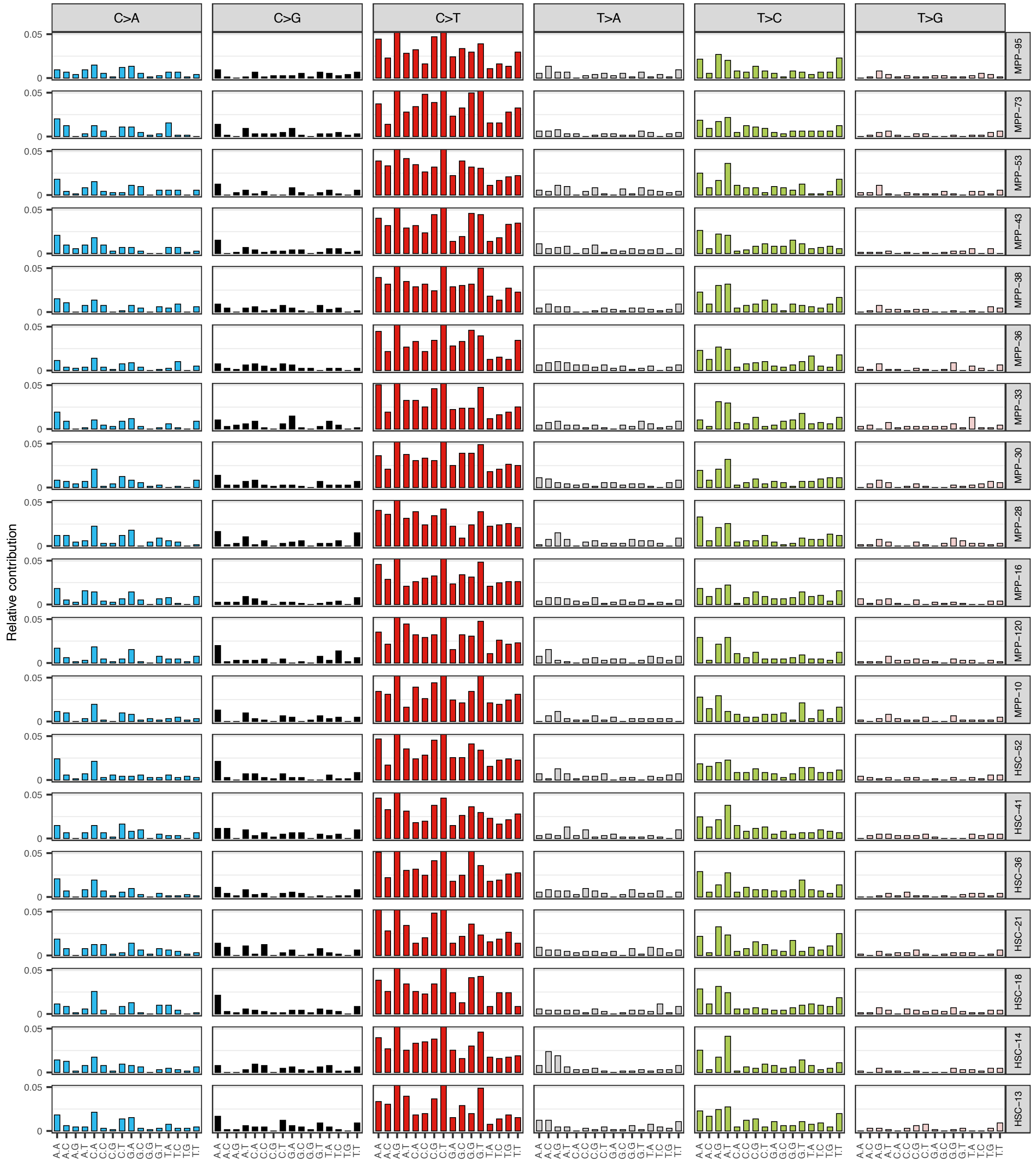

Supplemental Figure 14

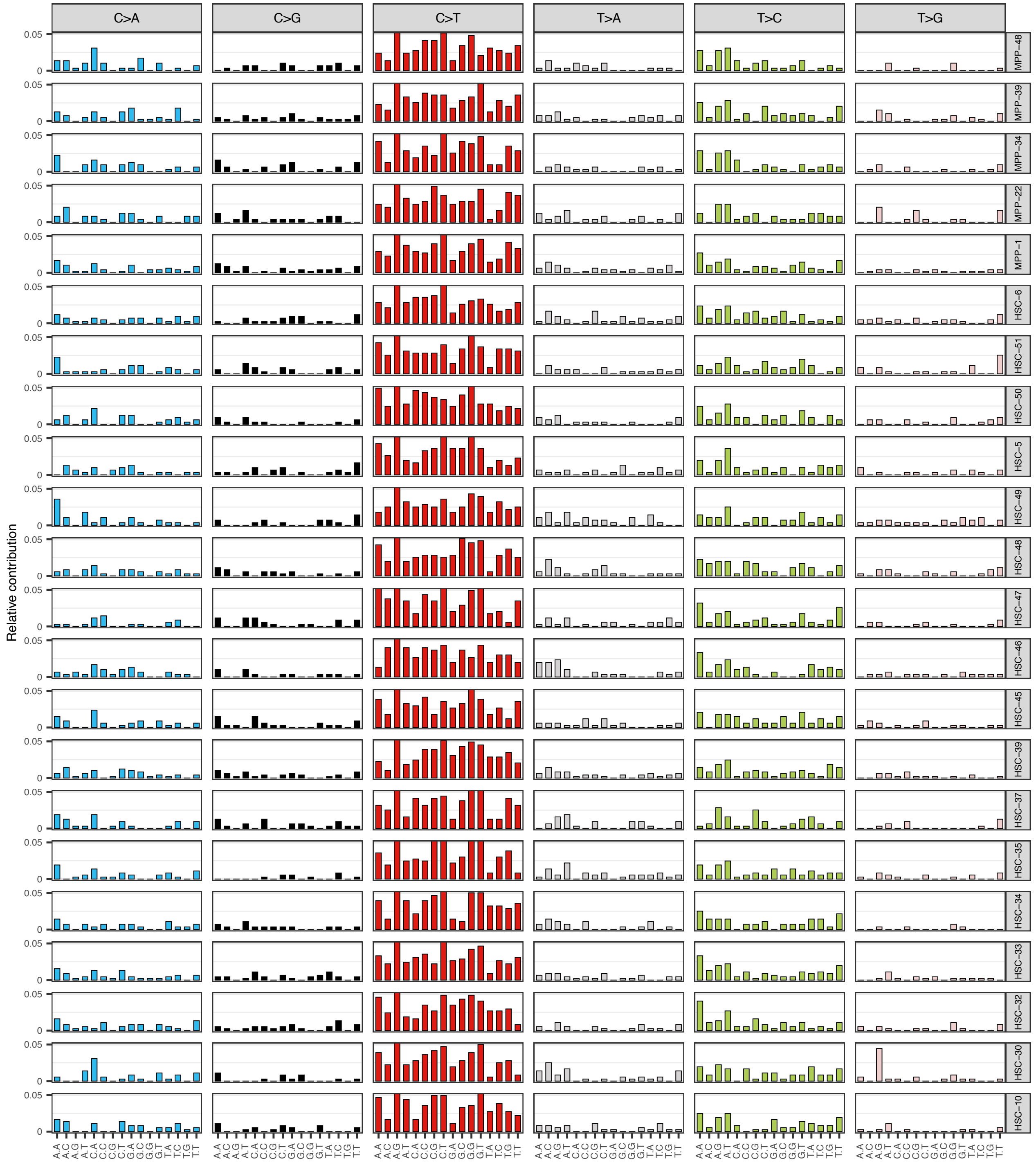

Supplemental Figure 15

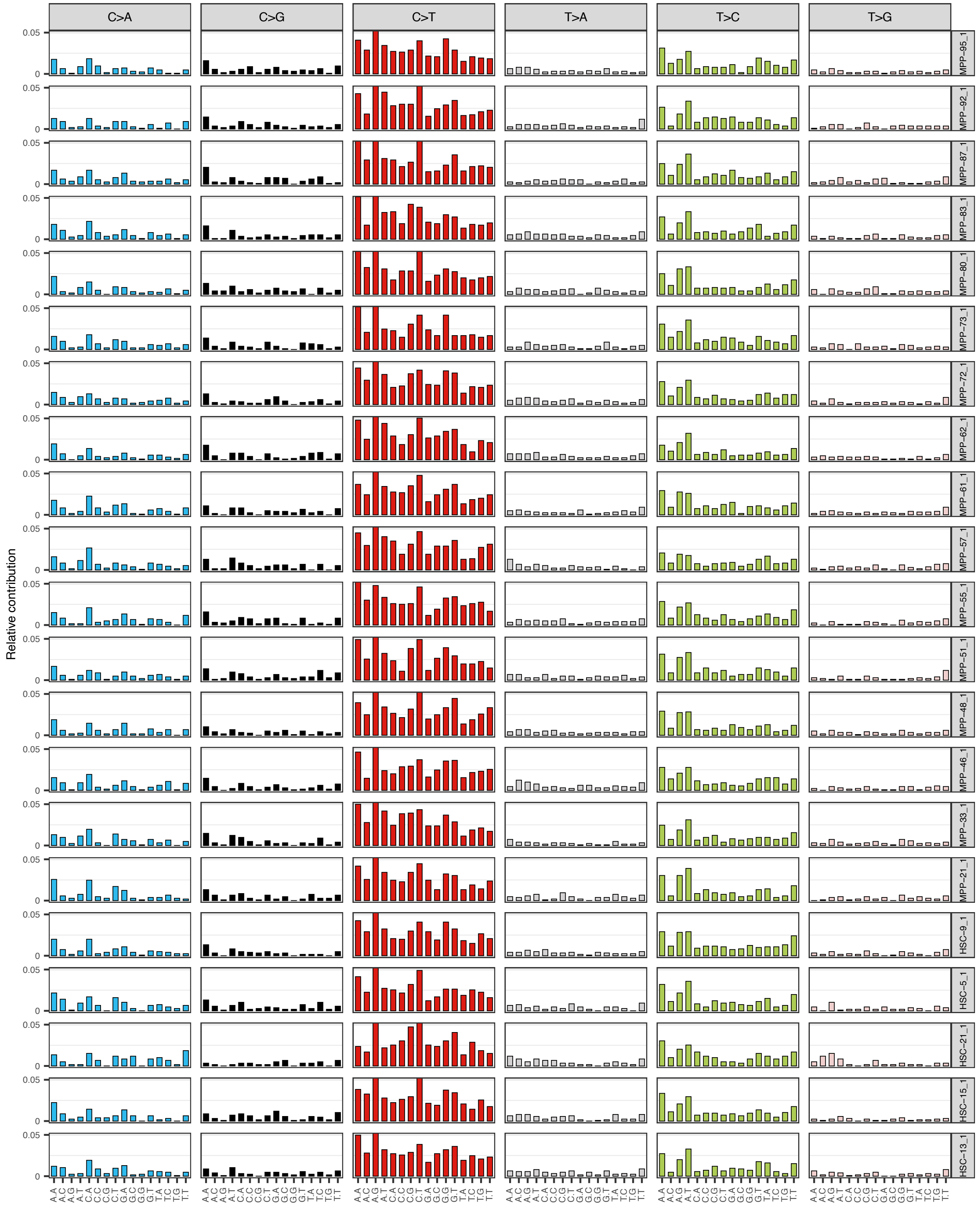

Supplemental Figure 16

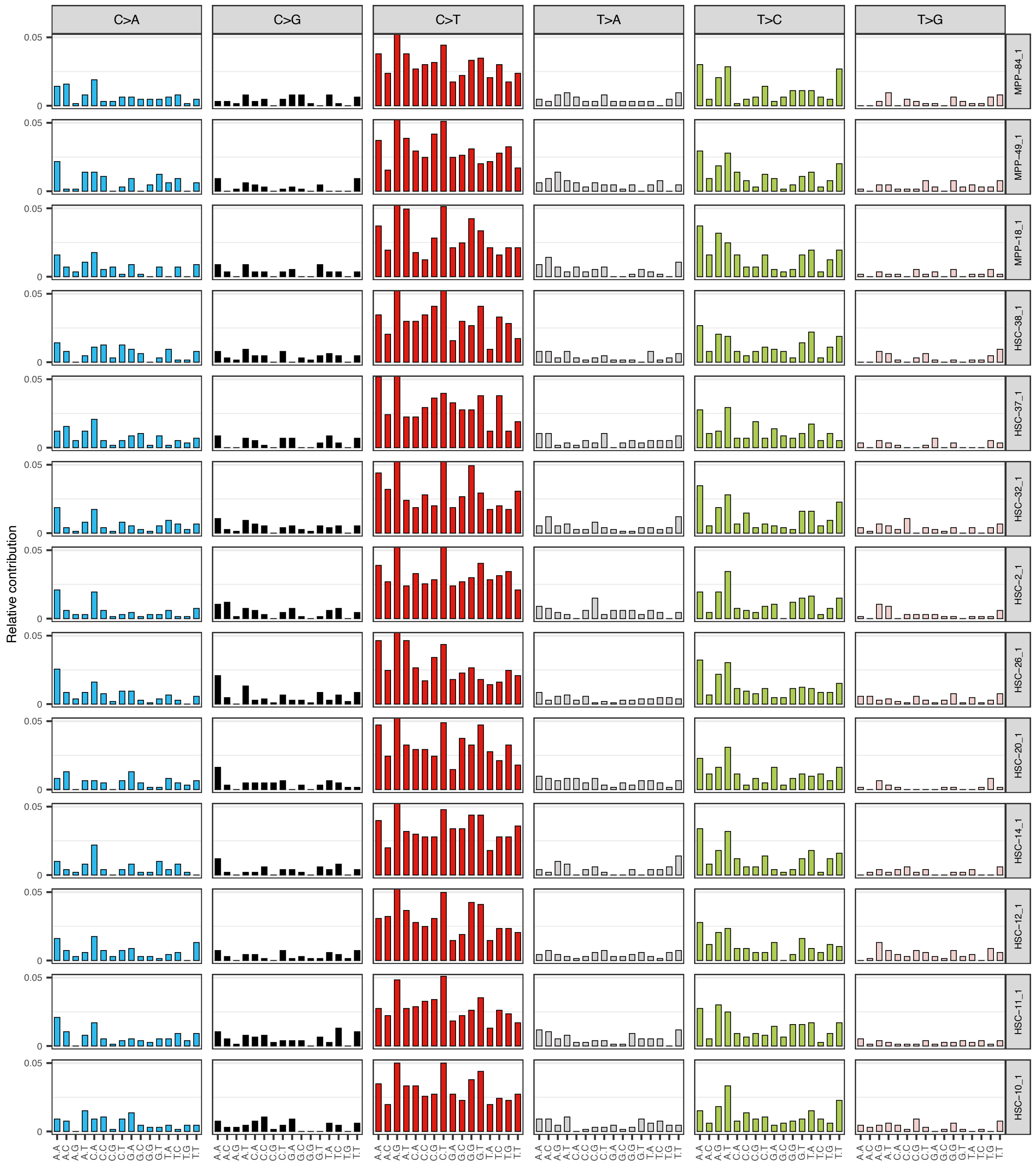

### Supplemental Figure 17

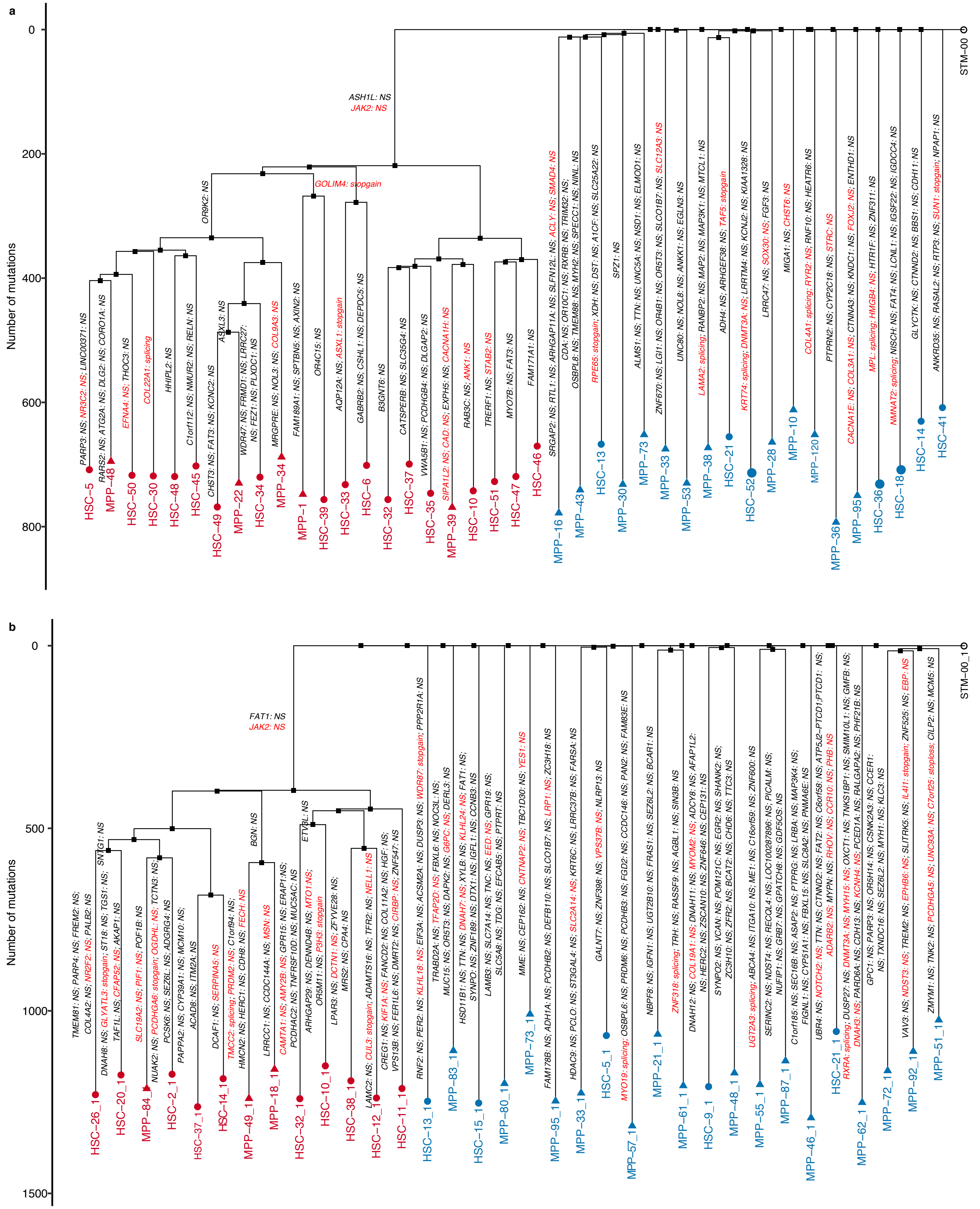

### Supplemental Figure 18

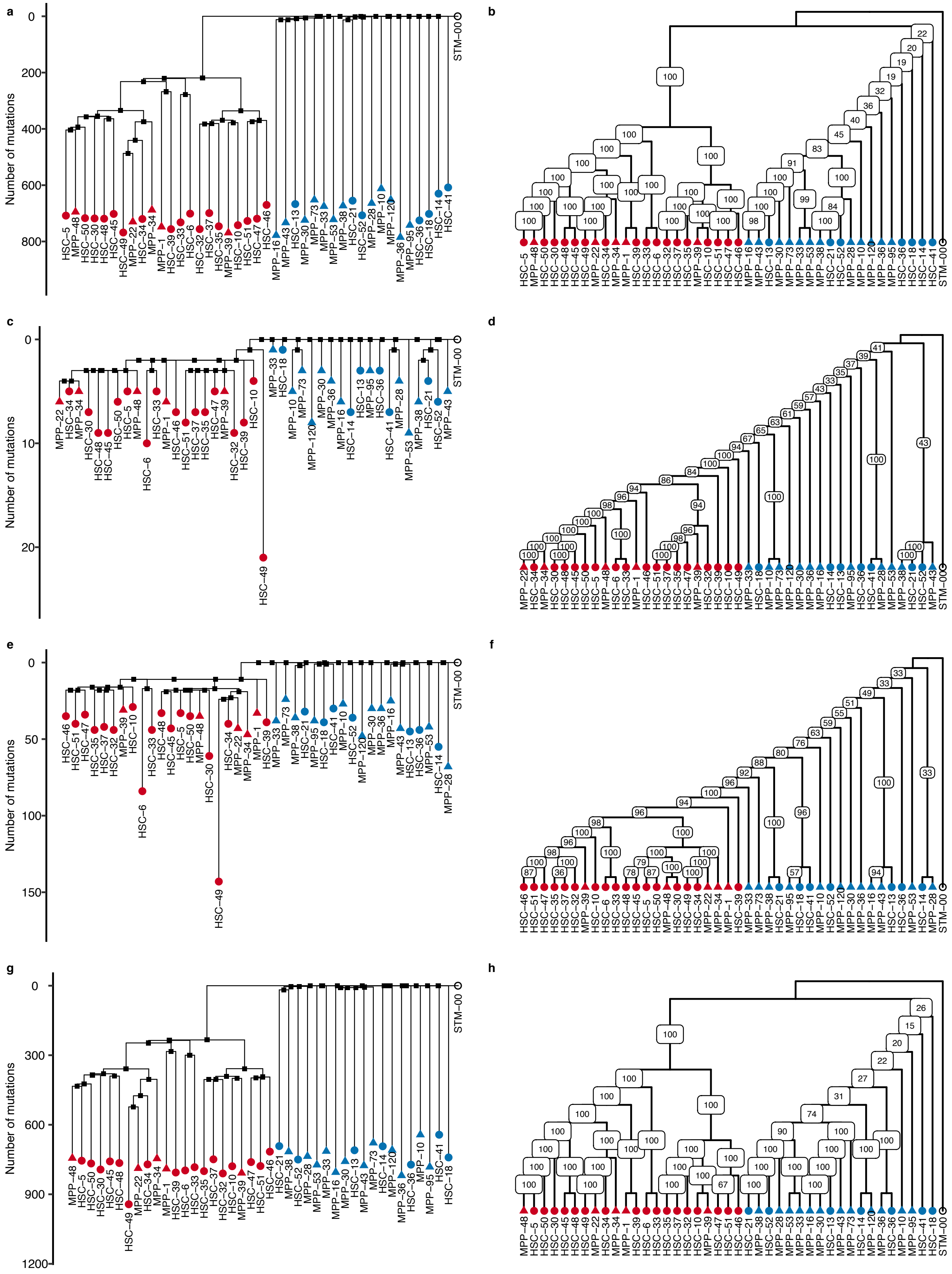

### Supplemental Figure 19

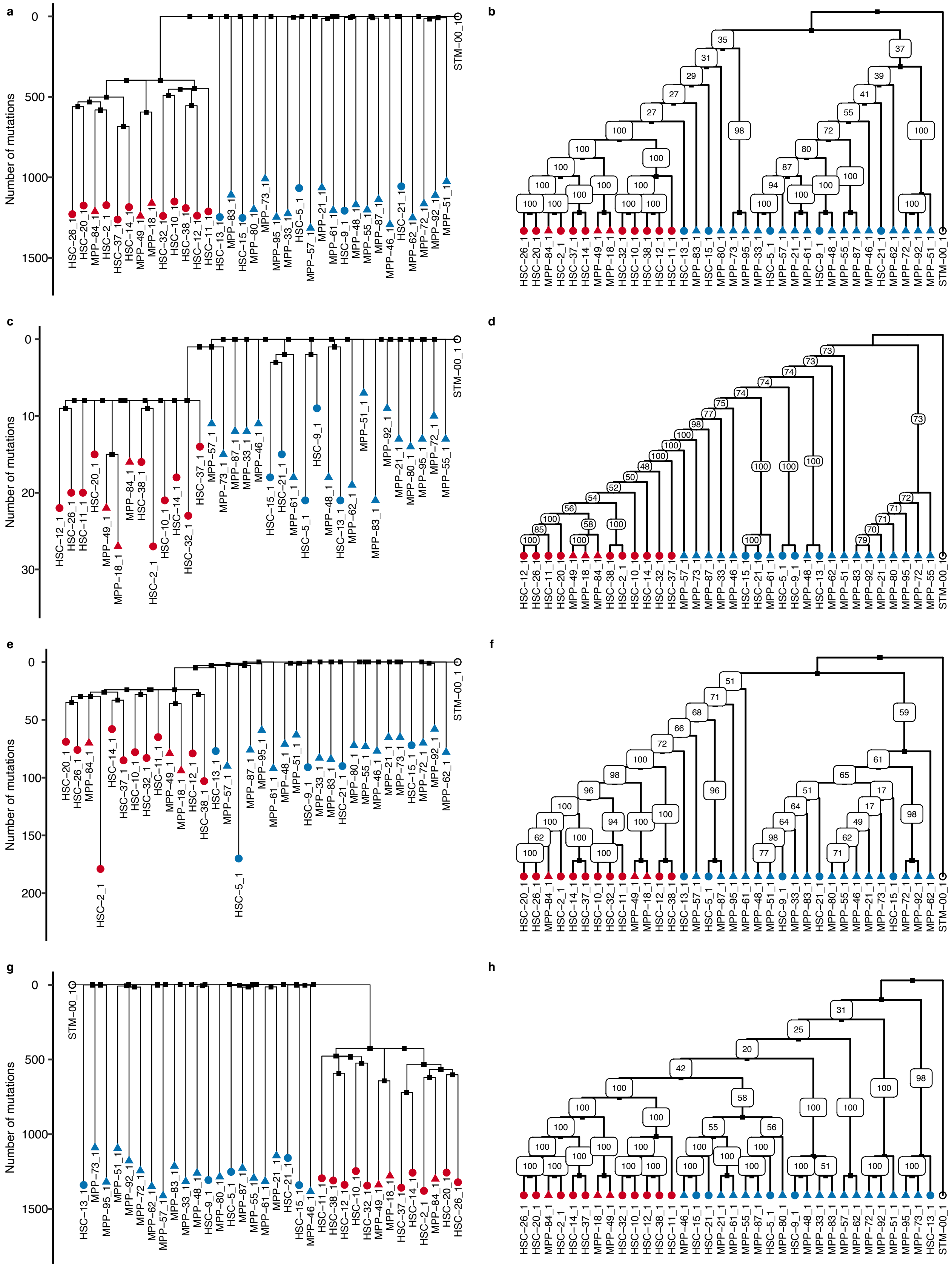

Supplemental Figure 20

a

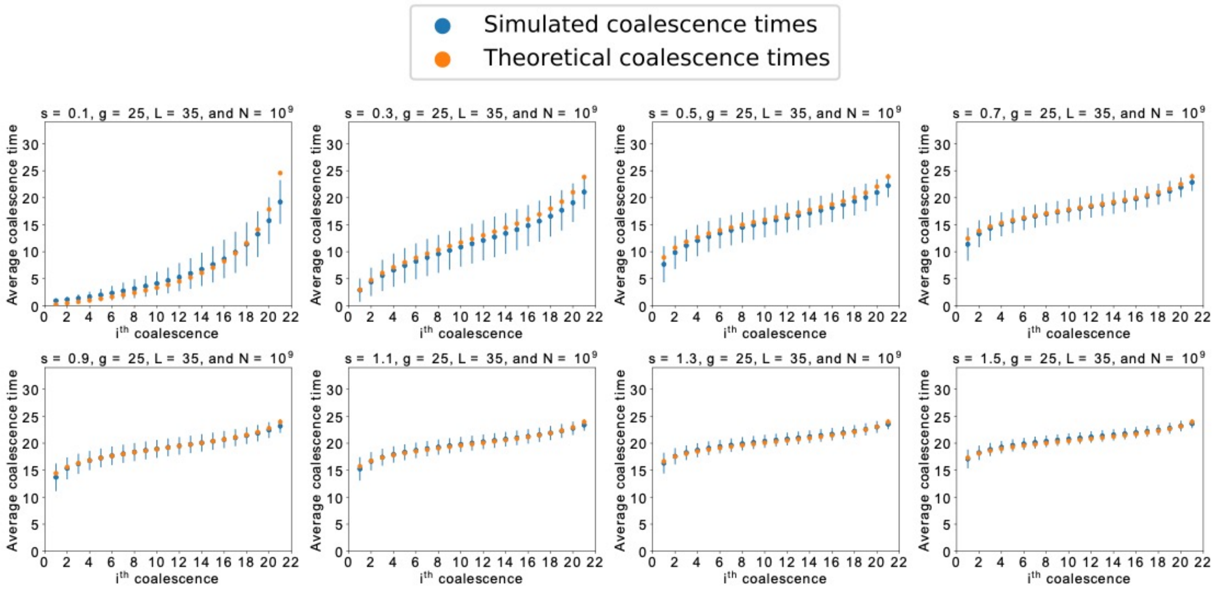


b

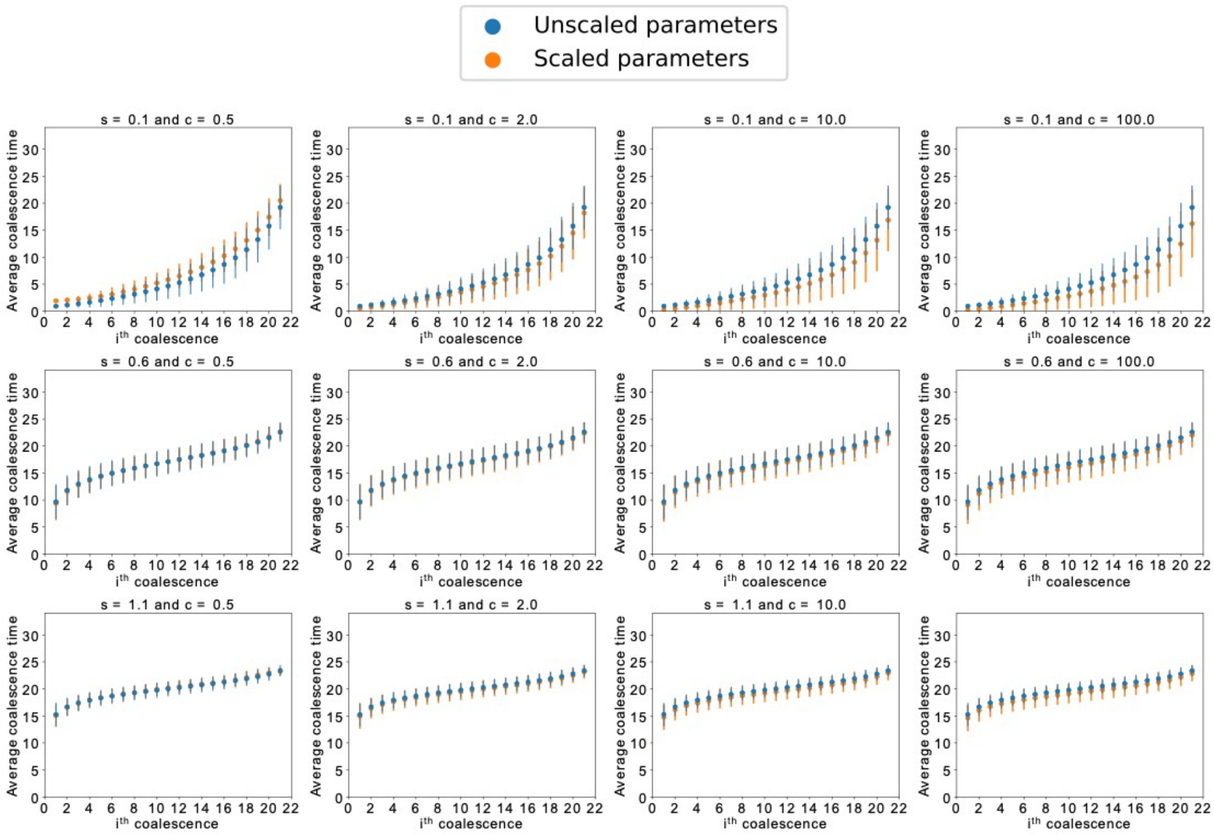

Supplemental Figure 21

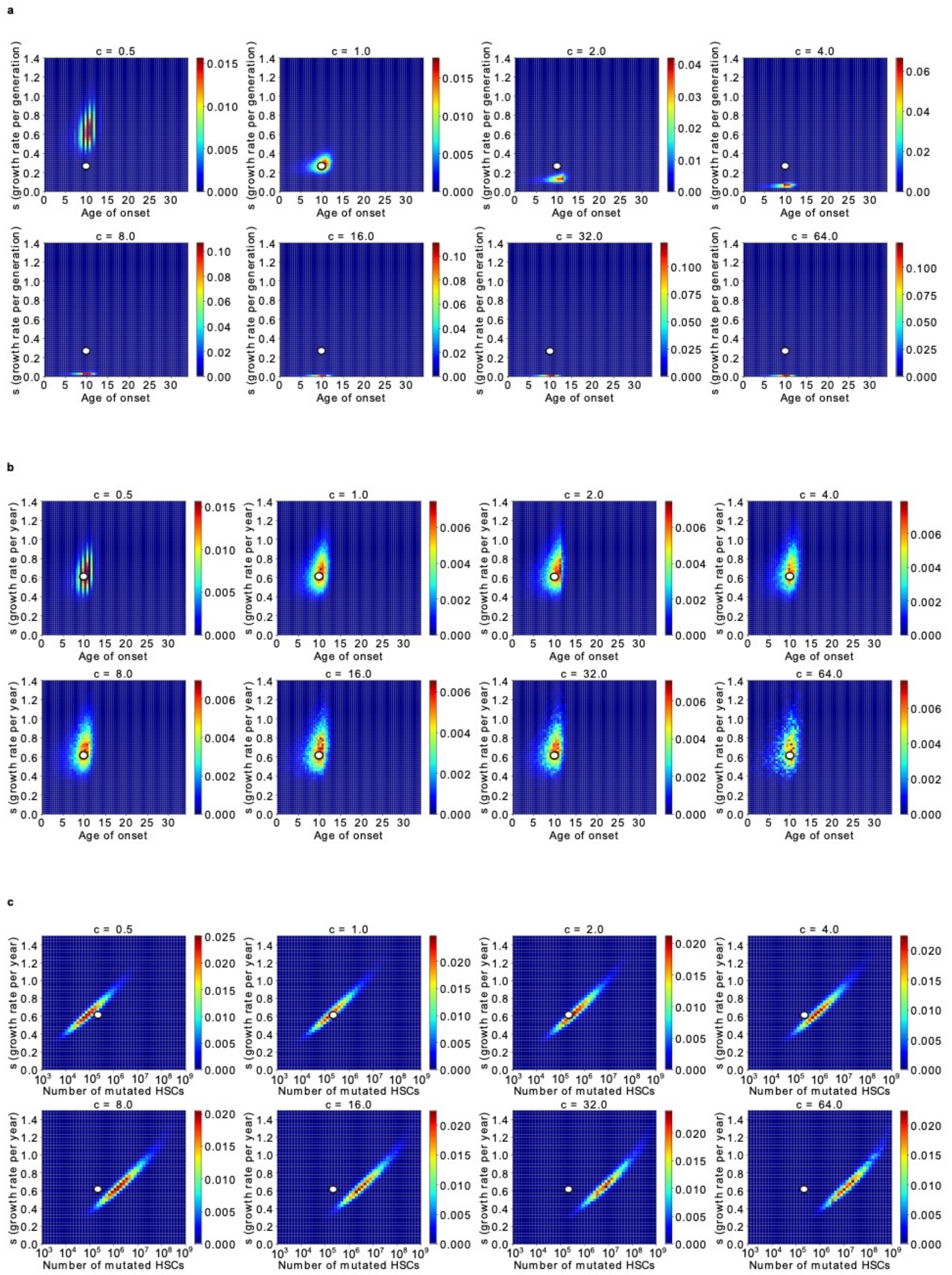

Supplemental Figure 22

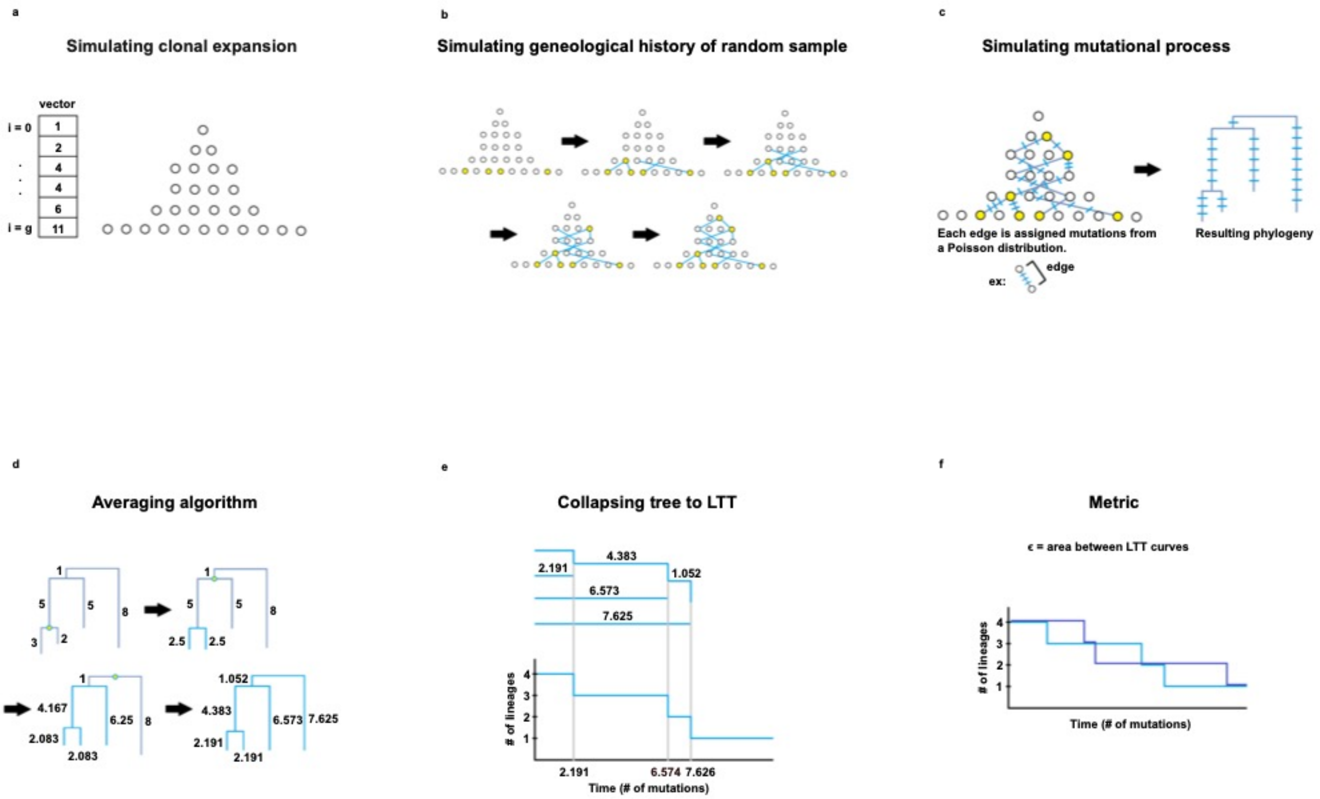

Supplemental Figure 23

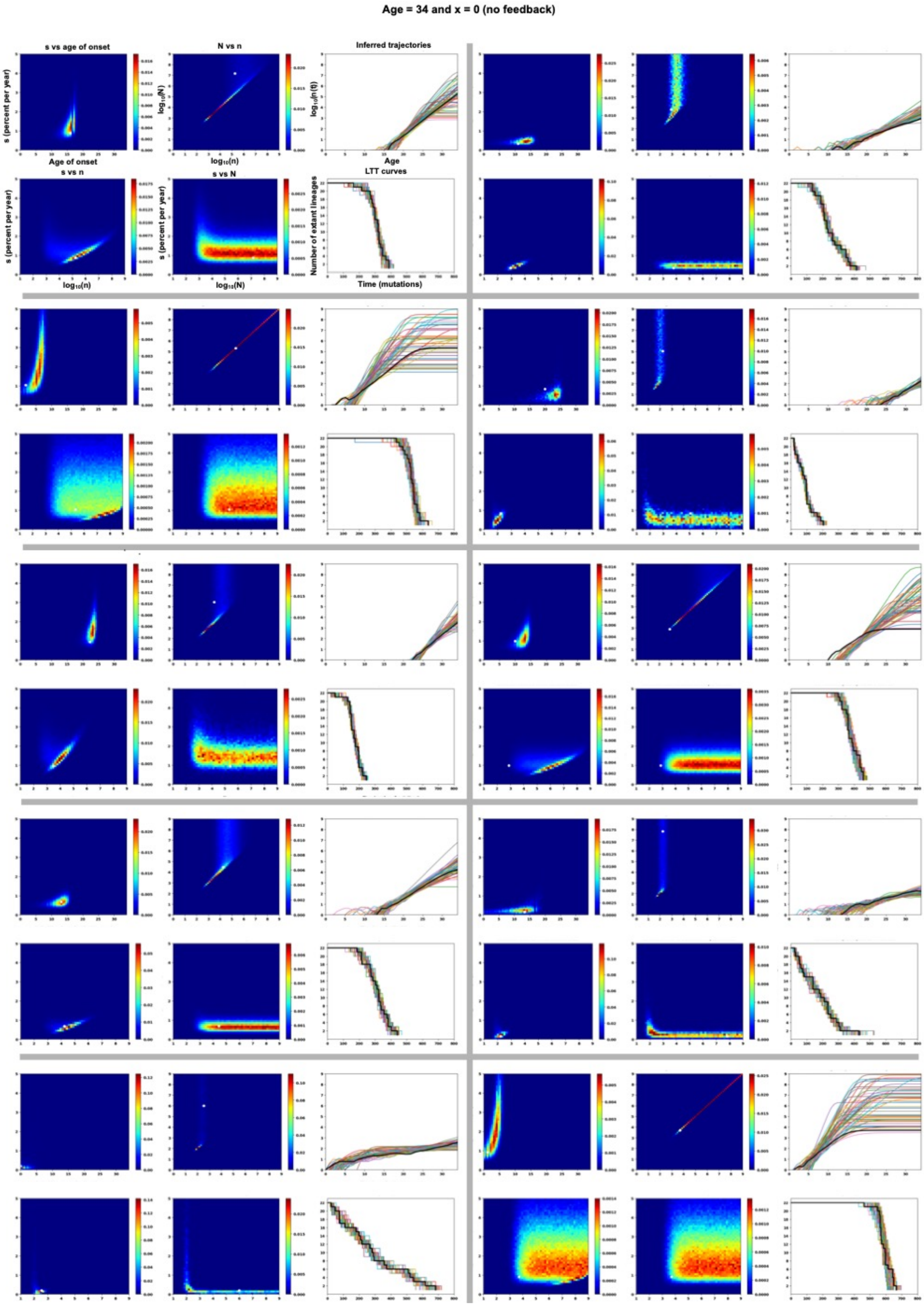

Supplemental Figure 24

**Age = 34 and x = 30 (with feedback)**

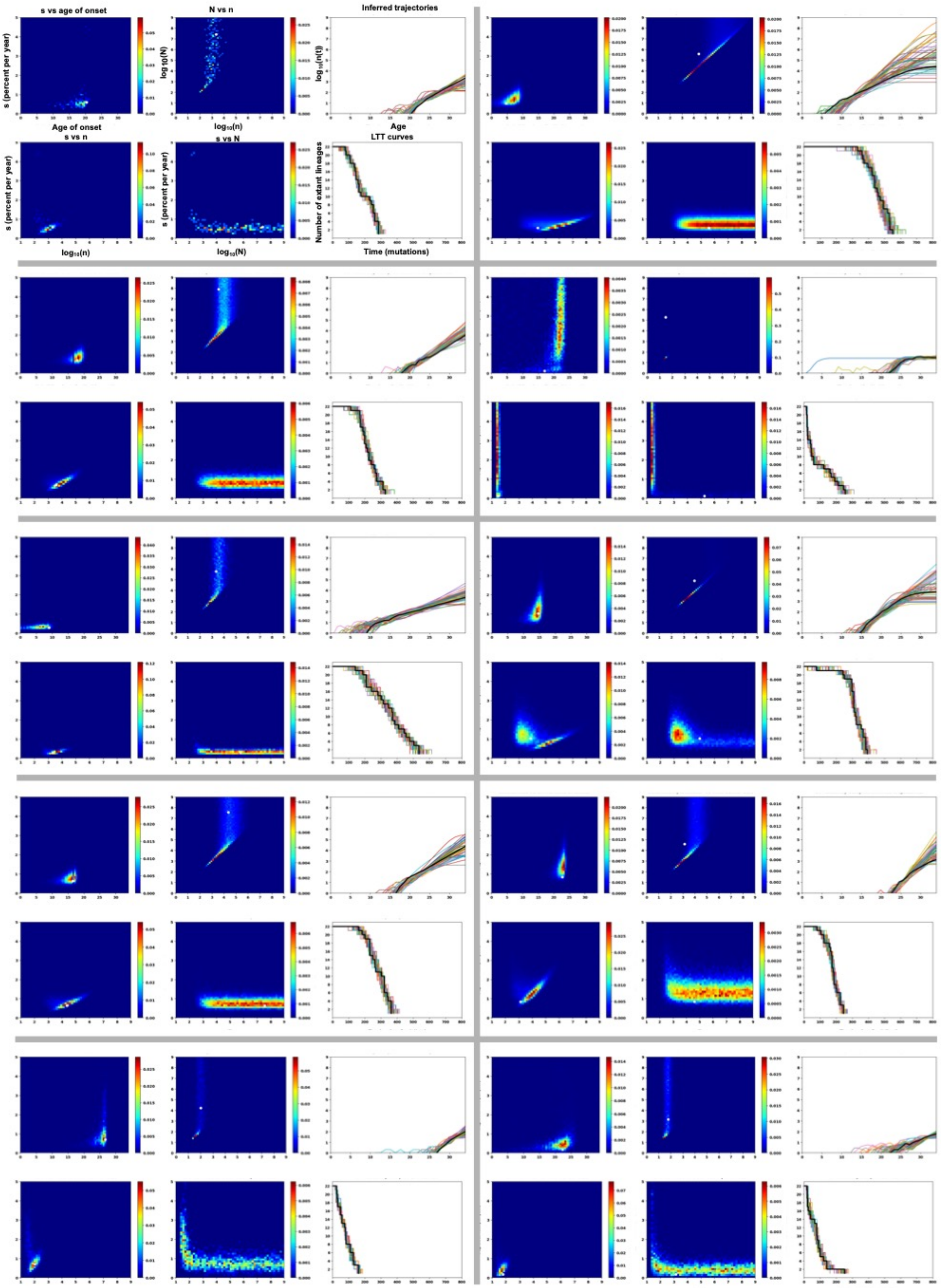

Supplemental Figure 25

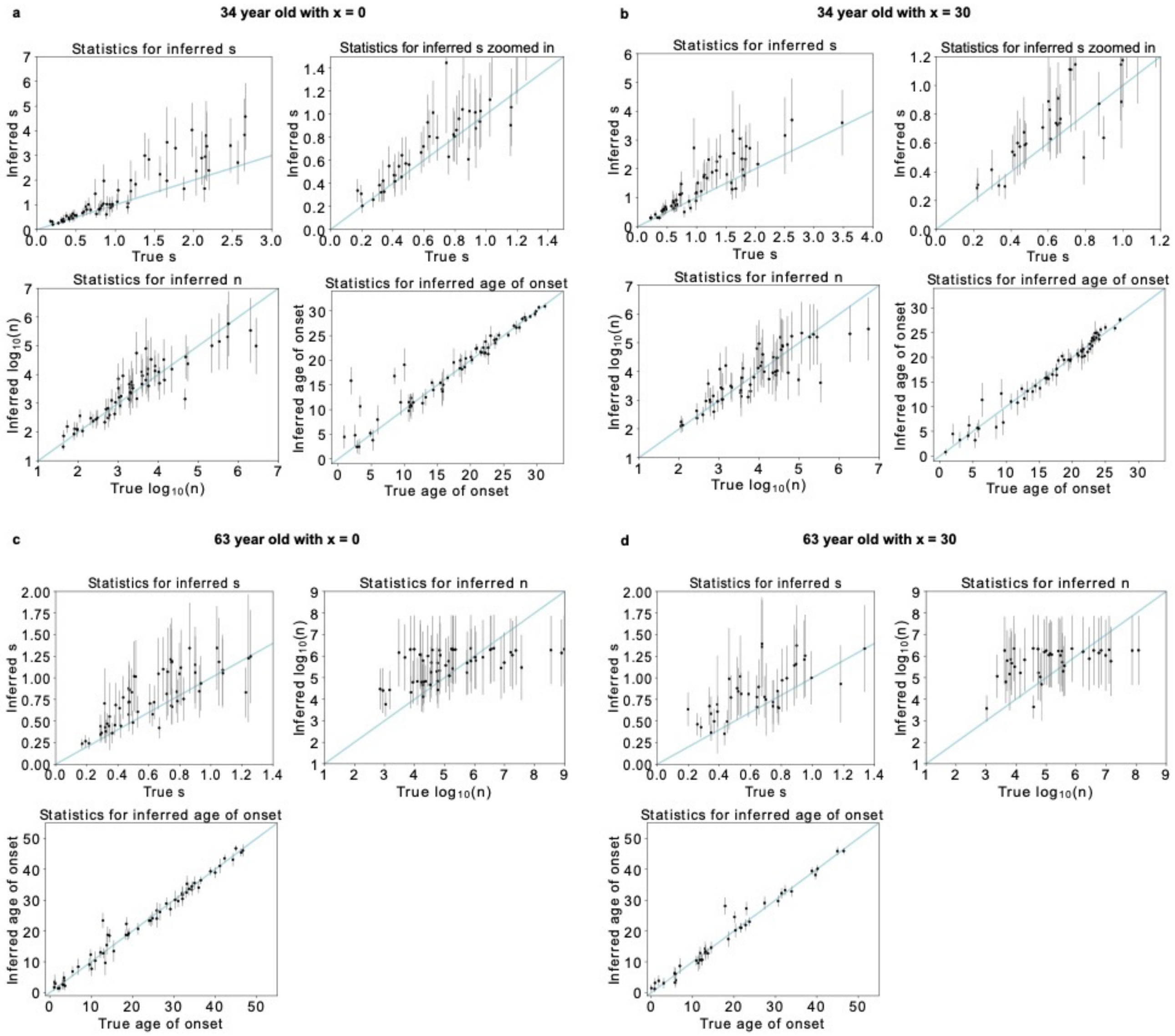

Supplemental Figure 26

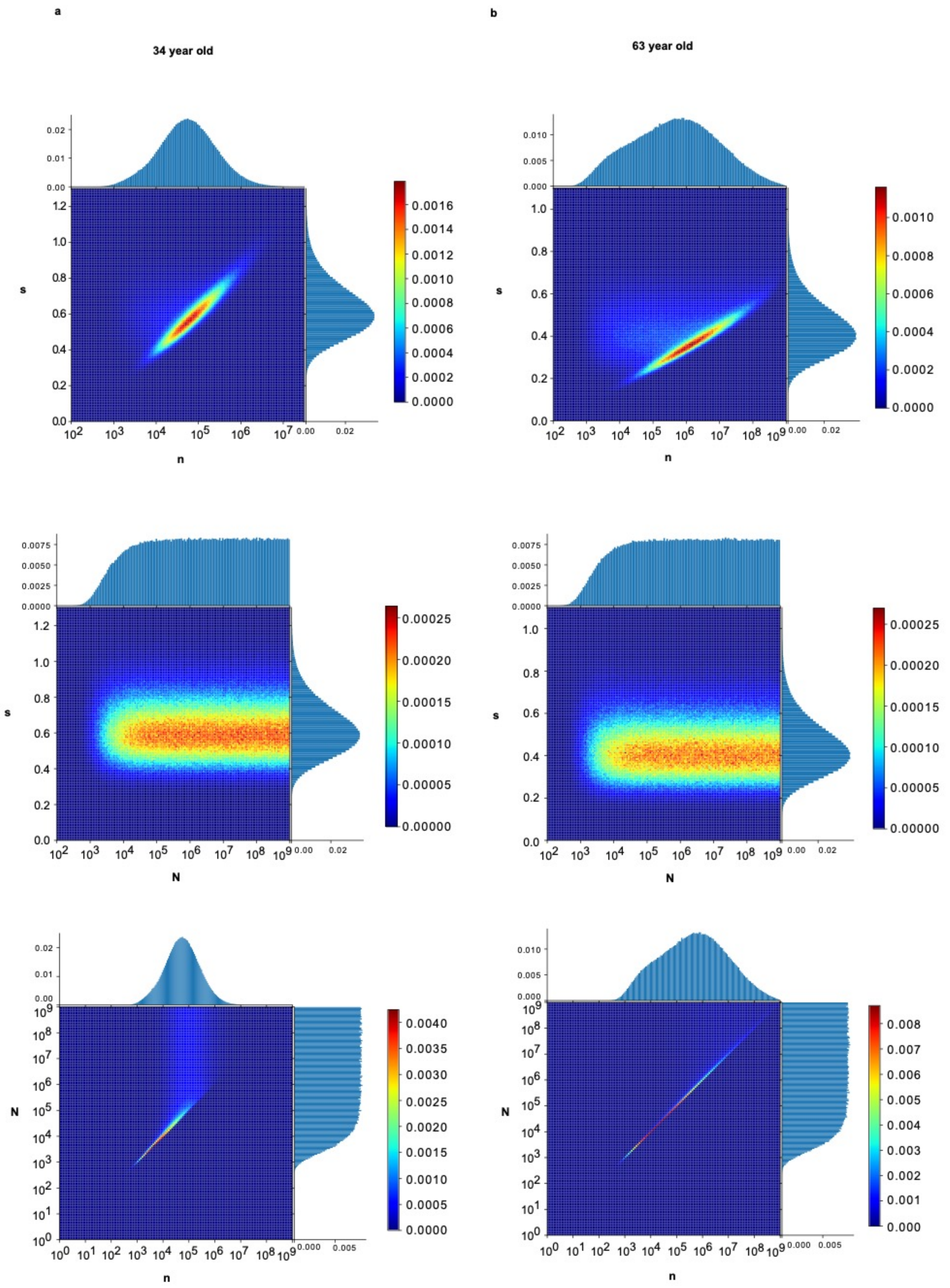

Supplemental Figure 27

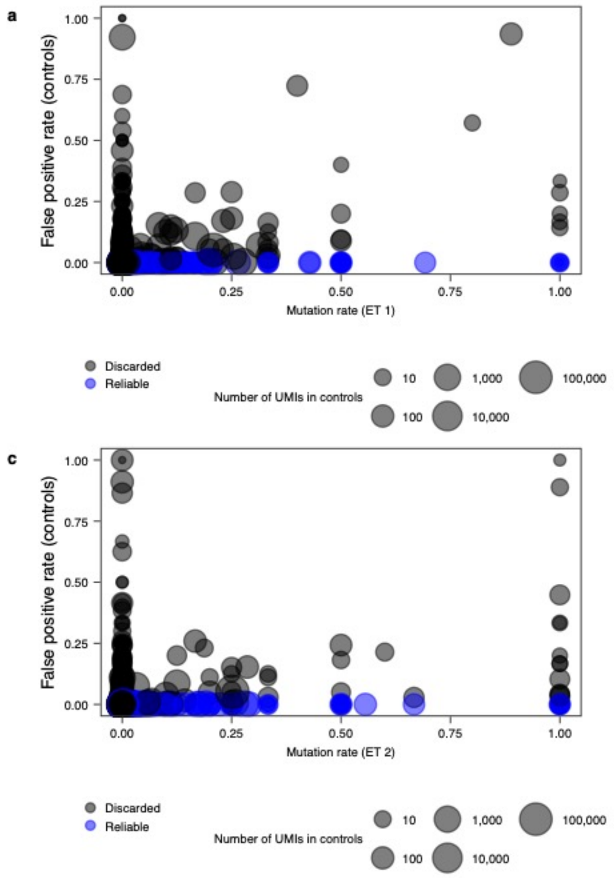

Supplemental Figure 28

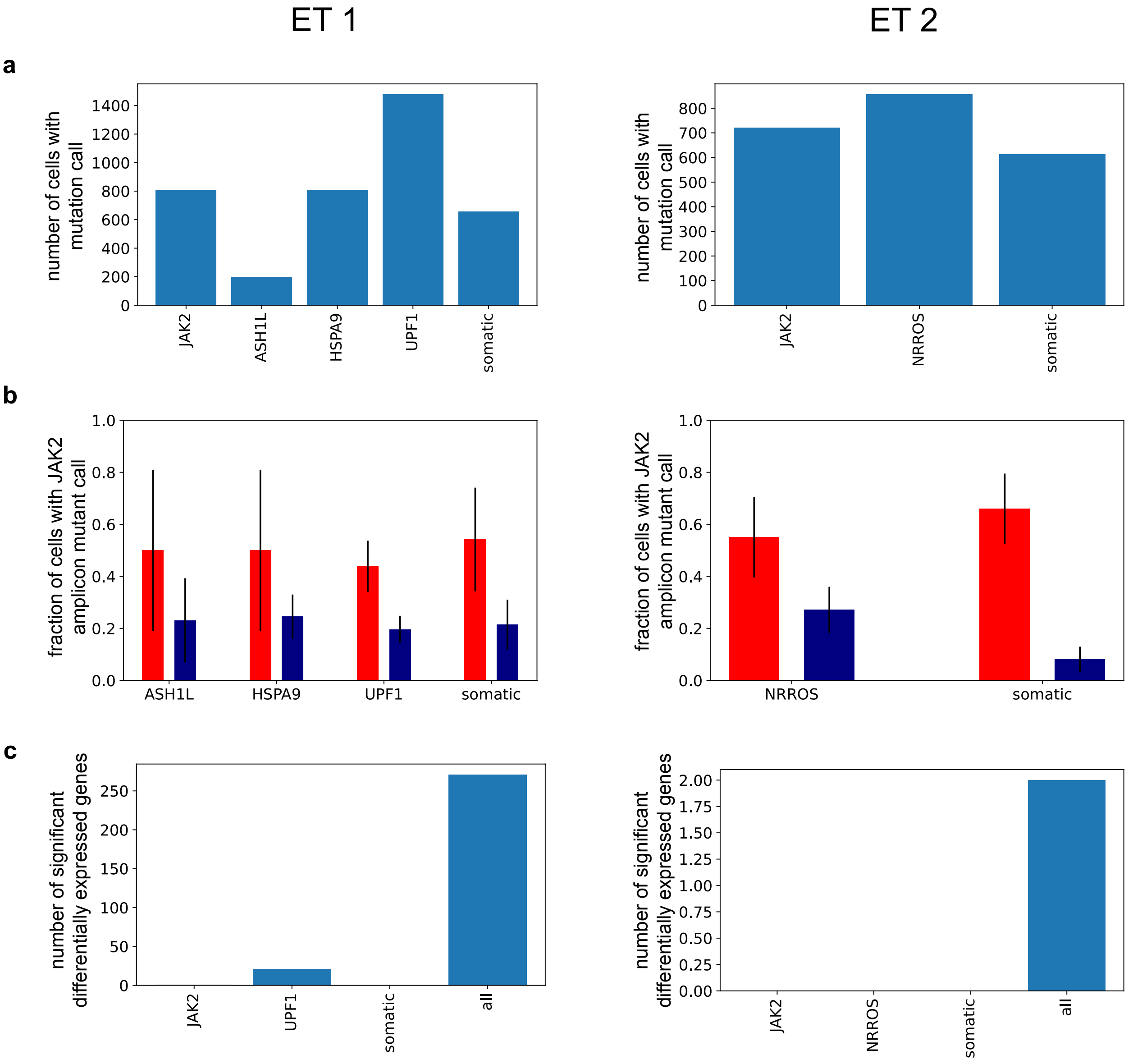

Supplemental Figure 29

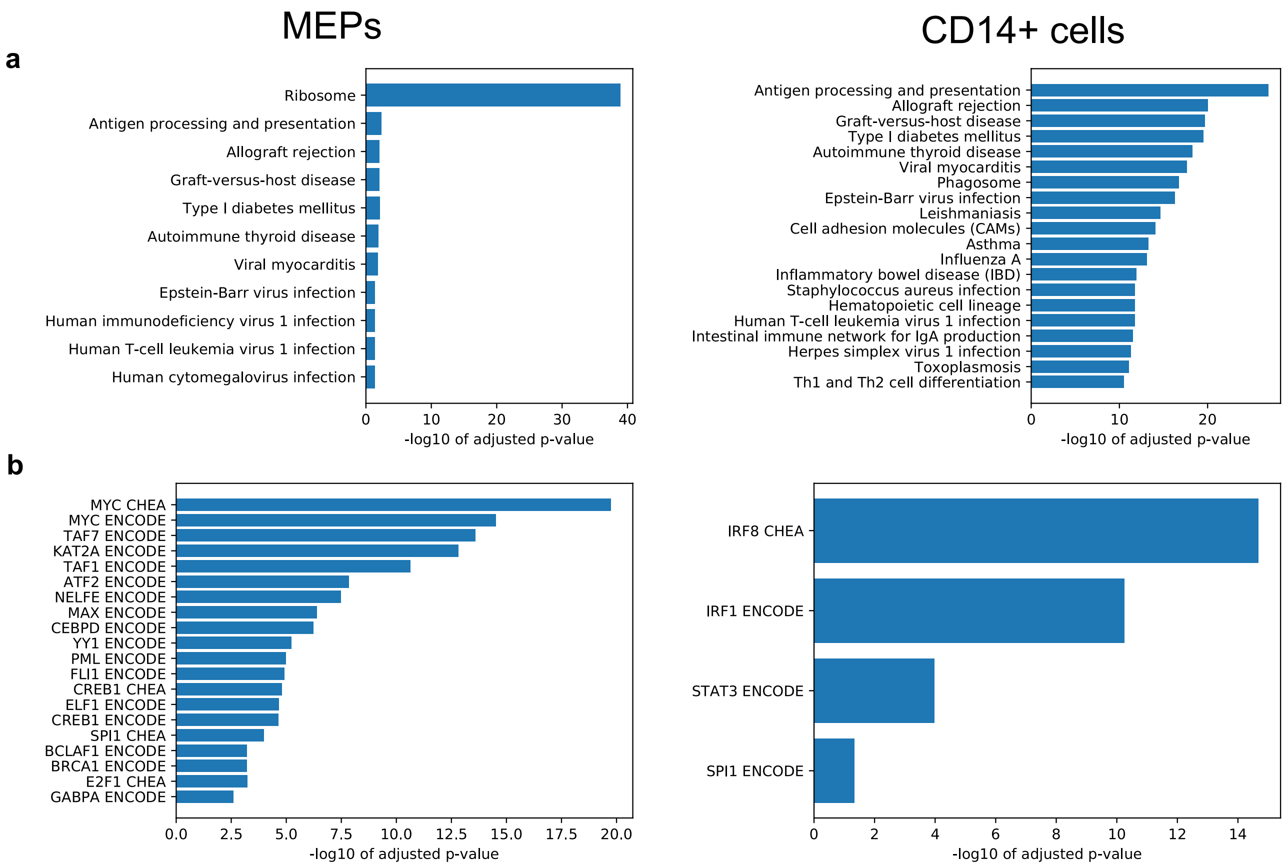

### Supplemental Figure 30

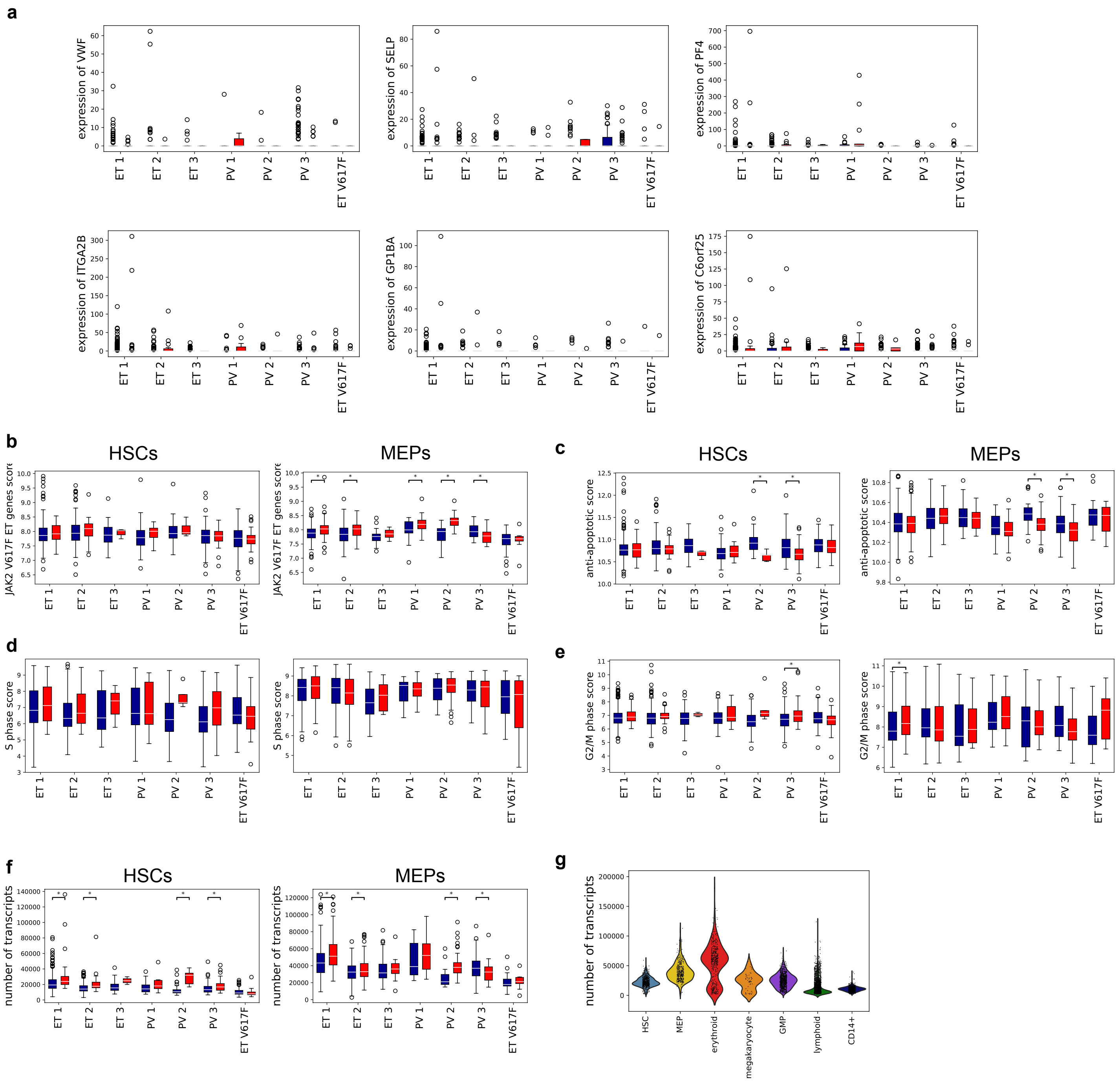

Supplemental Figure 31

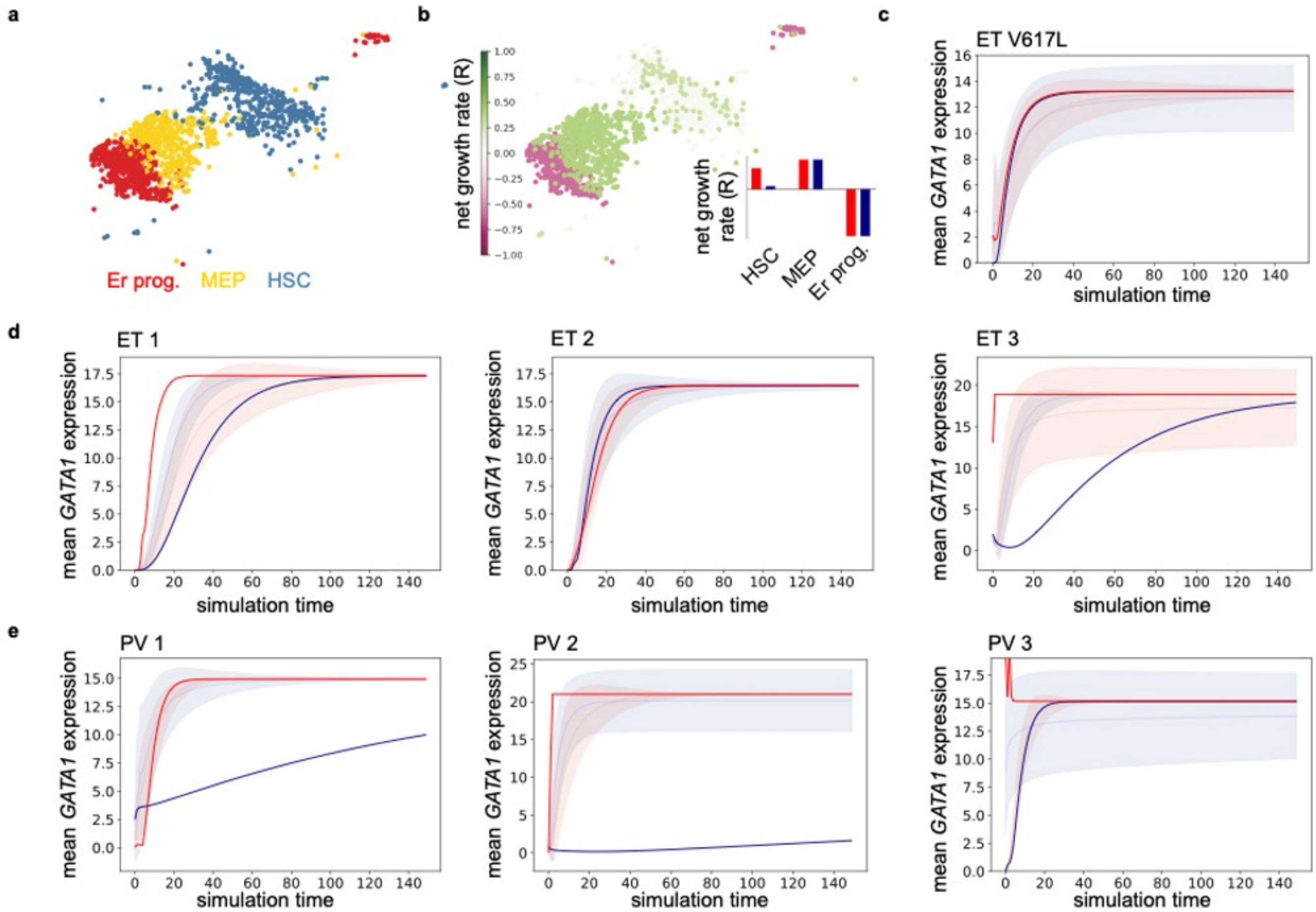

### Supplemental Figure 32

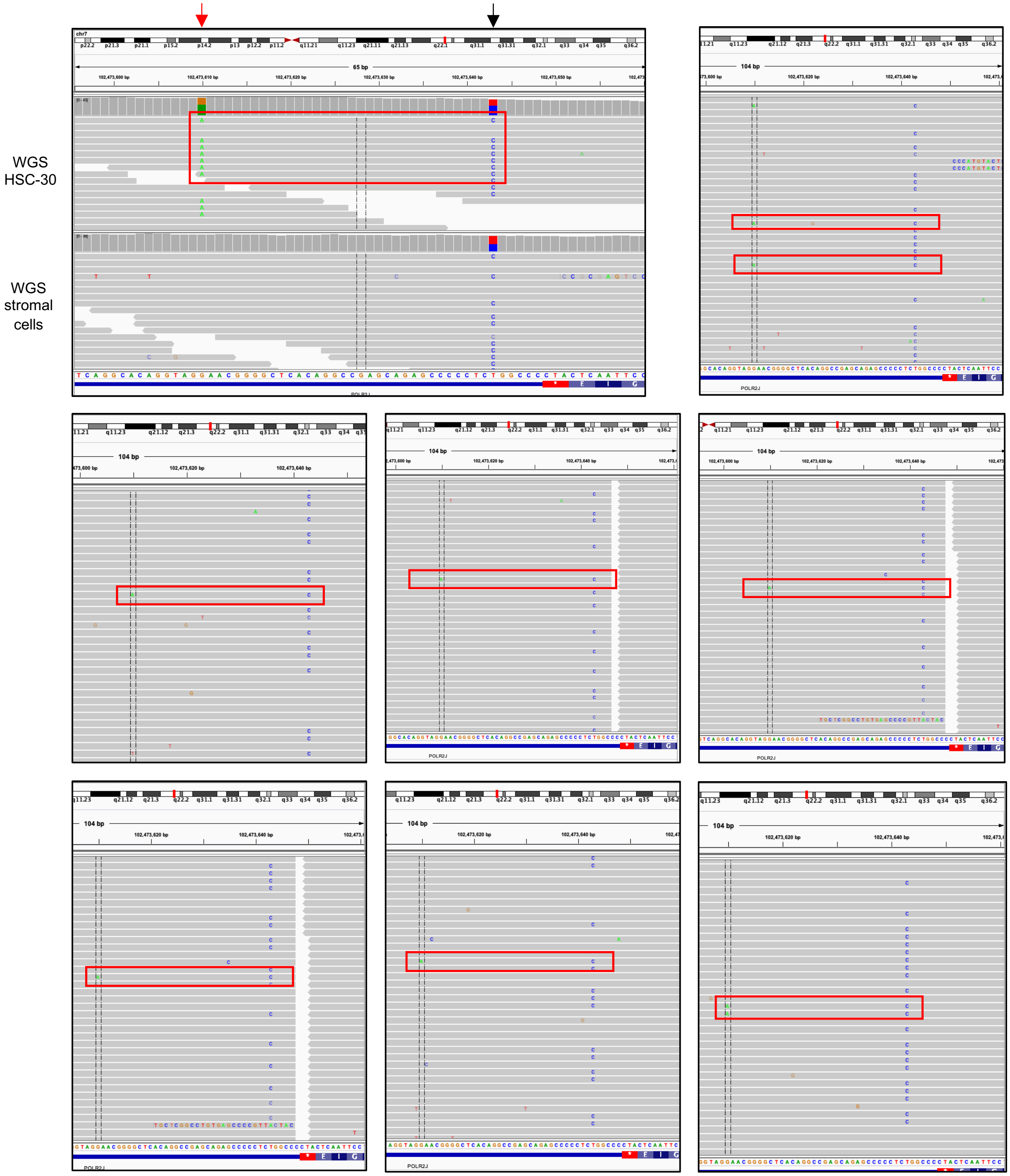

**FIGURE LEGENDS - SUPPLEMENT**

**Supplemental Figure 1. Primer designs, directions, locations, sequences of target mutation amplification and troubleshooting**. **a**. Schematic illustration of primer direction against target mRNA and the change of primer directionality during amplification of sc-cDNA. **b.** Schematic diagrams of the nested PCR from step 1 to step 5, respectively. **c.** Oligonucleotide sequences and localization of common primers and adaptors. **d.** Example primer positions and sequences of targeted mutation. **e.** Example TapeStation trace for QC and optimization from step 1 to 5. **f.** Number of reads vs rank of molecule and threshold of cell calling.

**Supplemental Figure 2. The amplicon libraries can be used to accurately identify the mutated cells. a.** In a control experiment HL60 (WT cells) were mixed with SET2 cells (heterozygous *JAK2* V617F mutation) and ran through the experimental and analysis pipeline. The two cell populations could be distinguished based on their transcriptional profiles: two distinct clusters were seen when transcriptomes of the cells were visualized using force directed graphs. Marker genes were used to identify the clusters as either HL60 or SET2 cells. Cells in which a WT *JAK2* transcript (left panel) or a mutated *JAK2* transcript (right panel) were detected in the amplicon libraries are shown as colored points. All other cells are shown in gray. As expected, *JAK2* WT transcripts were detected in both HL60 and SET2 cells, but *JAK2* V617F transcripts were only detected in SET2 cells. **b.** In another control experiment we combined the single-cell libraries of a *JAK2* V617F patient (ET 1) and that of the JAK2 V617L patient before the libraries were fragmented and indexed. We then ran the combined library through the experimental and analysis pipeline. The *JAK2* amplicon sequences could be mapped back to the library from which they originated based on their single-cell barcode. The plot shows the number of reads of each *JAK2* transcript detected for the V617F library (left) and V617L library (right). The color of the points denote whether the transcript sequence contained a V617F or V617L mutation. Blue dots on the left side and red dots on the right side correspond to incorrect mapping of a mutation to a single-cell barcode, most likely due to PCR crossover events during amplification. Above the threshold of 100 reads, the false positive rate is negligible.

**Supplemental Figure 3. CD34+ bone marrow cell types were identified in scRNA-seq data after integration of data from all samples. a.** UMAP of CD34-enriched bone marrow scRNA-seq data from all patients before batch correction, colored by donor as in **b**. **b-d.** UMAP of CD34-enriched bone marrow scRNA-seq data from all patients after Seurat batch correction, colored by donor (**b**), Louvain cluster (**c**), and final cell type identification (**d**). **e.** Expression of selected marker genes used to identify cell types. Colors denote raw counts of each marker gene. **f.** Fraction of each cell type in CD34-enriched bone marrow samples from each donor.

**Supplemental Figure 4. Differences in gene expression between ET, PV, and healthy HSCs. a.** Volcano plots of differential expression between all HSCs from ET, PV, and healthy HSCs. Genes found to be differentially expressed in all pairwise comparisons between different patient subsets are highlighted and colored by KEGG 2019 biological process group (gold: hematopoiesis-related, green: antigen presentation, blue: ribosomal, black: other). **b.** KEGG biological process gene set enrichment analysis results for genes that are differentially expressed in all pairwise comparisons.

**Supplemental Figure 5. *JAK2*-mutant cell fractions. a-g.** Fraction of cells in each compartment with a JAK2 mutation in patients ET 1-3 (**a-c**), PV 1-3 (**d-f**), and ET V617L (**g**). Error bars are 95% confidence intervals. Asterisks denote compartments with significantly different ($P < 0.05$, Fisher's exact test) *JAK2* mutant fractions from erythroid progenitors, after Benjamini-Hochberg multiple hypothesis correction.

**Supplemental Figure 6. *TET2* mutations occurred after *JAK2* mutations in PV 2 and PV 3. a-b.** Cells with *TET2* WT (a) or mutant (b) transcripts identified on UMAP projections of scRNA-seq data from PV 2 and PV 3. **c.** UMAPs with cells with *JAK2* and/or *TET2* mutant transcripts highlighted. **d.** Number of cells in the PV 2 and PV 3 scRNA-seq datasets with both *JAK2* and *TET2* transcripts observed that have a WT transcript for both *JAK2* and *TET2*, either a *JAK2* or *TET2* mutant transcript, or have both *JAK2* and *TET2* mutant transcripts observed.

**Supplemental Figure 7. CD14+ cells sampled from bone marrow are a mixture of intermediate and classical monocytes. a.** UMAPs of CD14+ cells from all patients showing expression of selected marker genes. Colors denote raw counts of each marker gene. **b-d.** CD14+ cell UMAPs colored by patient (**b**), kNN-smoothed

*JAK2* mutant fraction (**c**), and monocyte subset identified by Louvain clustering. **e.** Volcano plot showing differential expression results comparing intermediate monocyte cluster with classical monocyte cluster. **f**. Fraction of CD14+ cells in each patient that were identified as intermediate monocytes. **g**. Fraction of *JAK2* WT (blue) or *JAK2* mutant (red) CD14+ cells in each patient that were identified as intermediate monocytes.

**Supplemental Figure 8. Differences in gene expression between ET, PV, and healthy CD14+ bone marrow cells. a.** Volcano plots of differential expression between all HSCs from ET, PV, and healthy HSCs. Genes found to be differentially expressed in all pairwise comparisons between different patient subsets are highlighted and colored by KEGG 2019 biological process group (gold: hematopoiesis-related, green: antigen presentation, blue: ribosomal, black: other). **b.** KEGG biological process gene set enrichment analysis results for genes that are differentially expressed in all pairwise comparisons.

**Supplemental Figure 9. HSC and MPP isolation strategy by FACS.** Representative FACS plot for purifying HSCs and MPPs, starting with a CD34+ enriched bone marrow-derived cell suspension.

**Supplemental Figure 10. Correlation between the number of somatic point mutations and age at diagnosis for *JAK2*-mutant and wild-type (WT) colonies**. **a**. Correlation between age at diagnosis (years; x-axis) and the number of somatic SNVs (y-axis) detected in WT HSCs, JAK2-mutant HSCs, WT MPPs, and JAK2-mutant MPPS. Each dot corresponds to a single-derived colony, and the lines represent the regression through the origin. The estimated values for the slope and the 95% confidence intervals (CI) are shown.

**Supplemental Figure 11. Copy number for single-cell colonies derived from patient ET 1.** The total copy number is shown in black, whereas the minor copy number, which corresponds to the number of copies of the least amplified allele, is shown in red. The bulk whole-genome sequencing data from the stromal cells was used as the normal sample in all cases. Regions with no copy number information correspond to segments for which a reliable call could not be made.

**Supplemental Figure 12. Copy number for single-cell colonies derived from patient ET 2.** The total copy number is shown in black, whereas the minor copy number is shown in red. The bulk whole-genome sequencing data from the stromal cells was used as the normal sample in all cases. Regions with no copy number information correspond to segments for which a reliable call could not be made.

**Supplemental Figure 13. Patterns of somatic mutations for wild-type colonies from patient ET 1.** The relative fraction of each mutation type in the catalogue of point mutations detected in each colony is reported. Base substitutions are further stratified into categories based on the trinucleotide context in which the mutation occurs.

**Supplemental Figure 14. Patterns of somatic mutations for *JAK2*-mutant colonies from patient ET 1.** The relative fraction of each mutation type in the catalogue of point mutations detected in each colony is reported. Base substitutions are further stratified into categories based on the trinucleotide context in which the mutation occurs.

**Supplemental Figure 15. Patterns of somatic mutations for wild-type colonies from patient ET 2.** The relative fraction of each mutation type in the catalogue of point mutations detected in each colony is reported. Base substitutions are further stratified into categories based on the trinucleotide context in which the mutation occurs.

**Supplemental Figure 16. Patterns of somatic mutations for *JAK2*-mutant colonies from patient ET 2.** The relative fraction of each mutation type in the catalogue of point mutations detected in each colony is reported. Base substitutions are further stratified into categories based on the trinucleotide context in which the mutation occurs.

**Supplemental Figure 17. Mapping somatic mutations detected in the WGS data to the consensus tree for patients ET 1 and ET 2.** The consensus lineage trees across 100 bootstrap resamples reconstructed using somatic point mutations as input for patient ET 1 (**a**) and ET 2 (**b**). Mutations predicted to be pathogenic

(Methods) are shown in red. The mutations are listed along each branch in no particular order. NS: nonsynonymous.

**Supplemental Figure 18. Consensus lineage trees for patient ET 1.** The consensus lineage trees across 100 bootstrap resamples reconstructed using the extended majority rule method when using the following mutation types as input are shown: (**a**) somatic single-nucleotide mutations, (**c**) microsatellite mutations, (**e**) indels, and (**g**) single-nucleotide mutations, microsatellite mutations, and indels combined. Consensus lineage trees reporting the percentage of trees generated using bootstrap resamples that contained the clade shown in the consensus tree generated using (**b**) somatic single-nucleotide mutations, (**d**) microsatellite mutations, (**f**) indels, and (**h**) single-nucleotide mutations, microsatellite mutations, and indels combined. The white circles denote the outgroup.

**Supplemental Figure 19. Consensus lineage trees for patient ET 2.** The consensus lineage trees across 100 bootstrap resamples reconstructed using the extended majority rule method when using the following mutation types as input are shown: (**a**) somatic single-nucleotide mutations, (**c**) microsatellite mutations, (**e**) indels, and (**g**) single-nucleotide mutations, microsatellite mutations, and indels combined. Consensus lineage trees reporting the percentage of trees generated using bootstrap resamples that contained the clade shown in the consensus tree generated using (**b**) somatic single-nucleotide mutations, (**d**) microsatellite mutations, (**f**) indels, and (**h**) single-nucleotide mutations, microsatellite mutations, and indels combined. The white circles denote the outgroup.

**Supplemental Figure 20. The analytical calculation of coalescent times matches the simulations results. a.** We verify that our analytical calculation of the average coalescence times of our model matches the empirically observed coalescence times. Blue dots show the average coalescence times computed empirically for different s by simulating a large number of cancer expansions, constructing trees, and then averaging over the times until coalescence. Error bars denote 1 std in the coalescence times. Orange dots show the average coalescence times derived analytically for comparison. **b.** Our mathematical derivation shows that scaling the number of generations by a factor of *c* while maintaining the percent growth per year, *s*, fixed does not change the average coalescence times of trees. We used *s* = 0.1, 0.6, and 1.1 in percent growth per year corresponding to rows 1, 2, and 3 respectively, scaled the number of generations by different factors of *c*, and computed the average coalescent times empirically. Blue dots are the empirically computed coalescent times before scaling the number of generations by *c*, and the orange dots are the empirically computed coalescent times after scaling the number of generations by *c*. The average coalescent times using scaled parameter values always lie within 1 std of the average coalescent times using unscaled parameters. The small deviations are due to the fact that *s* in growth per year is not sufficiently small when the number of generations is not scaled. As *c* increases, increasing the number of generations, the growth per generation decreases to maintain constant percent growth per year. As the growth per generation decreases, the coalescent times converge according to our mathematical derivation.

**Supplemental Figure 21. Fitness can be inferred without knowing the number of generations.** Ground truth growth dynamics was simulated, and multiple ABC inferences were carried out under different *c* values. For each value of *c*, the number of generations used in ABC was multiplied by c for the ABC inference. **a.** inferred *s* (percent per generation) vs inferred age of onset of disease (years) are plotted for different values of *c*. As expected, increasing *c*, and hence the number of generations, decreases the growth per generation to keep the percent growth per year invariant. **b.** Inferred *s* was then converted to growth per year. As suggested by our derivation, the inferred *s* in growth per year remains invariant as the number of generations is scaled. **c.** The inferred *s* (percent per year) vs the inferred number of mutated HSCs at the final time-point is plotted for different values of *c*. As expected, increasing *c*, and hence the number of generations, decreases the growth per generation and thereby the rate of stochastic extinction. The number of mutated HSCs must then fluctuate to a larger population size early on to escape extinction. This results in a larger number of mutated stem cells at the final time-point.

**Supplemental Figure 22. Schematic of the Approximate Bayesian computation**. First, the parameters that determine the growth dynamics (i.e. fitness, population size, total number of stem cells, and age of onset) are randomly drawn from a prior distribution. **a.** The clonal expansion of the cancer cells given the selected parameters is simulated (Methods). **b.** A specified number of cells is randomly chosen from the final population. The lineage tree is reconstructed for these cells. **c.** We assign the number of mutations accrued along each branch by drawing from a Poisson distribution with the mean set at the mutation rate. Those mutations are shown pictorially as blue dashes on the tree. The resulting tree is a phylogeny with branch lengths in mutations as opposed to generations. **d.** Next, the branch lengths are scaled so that the total number of mutations from the root of the tree to each leaf node is the same (Methods). **e.** The rescaled trees are then plotted as LTT curves. **f.** A distance is computed between the LTT curve of the simulated tree and the observed tree. If the distance is below a threshold, the initial set of parameters is retained, otherwise they are discarded. This process is iterated.

**Supplemental Figure 23. *In silico* validation of the inference algorithm.** To validate the inference algorithm, we generated simulated ground truth growth dynamics and then inferred the parameters using ABC (described in Methods). In these simulations, the ground truth dynamics were simulated using the same model as in the ABC. In all the heatmaps, the ground truth parameters are shown as white dots. In the traces, the ground truth is shown in black. 10 illustrative examples are shown. For each example, the heatmaps of inference of *s* (fitness parameter), *n* (number of cancer cells), *N* (total populations size), and *g* (age of onset) are shown alongside the LTT curves that were retained and the inferred trajectories of population growth.

**Supplemental Figure 24. *In silico* validation of the inference algorithm.** To validate the inference algorithm, we generated simulated ground truth growth dynamics and then inferred the parameters using ABC (described in Methods). In these simulations, the ground truth dynamics were simulated with a feedback where the fitness of cancer cells decreased as the population size increased. The growth dynamics for generating trajectories for ABC did not incorporate the feedback. In all the heatmaps, the ground truth parameters are shown as white dots. In the traces, the ground truth is shown in black. 10 illustrative examples are shown. For each example, the heatmaps of inference of *s* (fitness parameter), *n* (number of cancer cells), *N* (total populations size), and *g* (age of onset) are shown alongside the LTT curves that were retained and the inferred trajectories of population growth. Taken together, accurate inference is possible even if additional features, such as feedback, are not incorporated in the ABC dynamics.

**Supplemental Figure 25. ABC accurately infers model parameters**. To verify that the inference framework can accurately infer model parameters, we simulated growth dynamics across a range of scenarios (corresponding to different parameter values) and inferred the parameters using ABC. The inferred parameter values were plotted against their true values for *s* (percent per year), *n*, and age of onset of the disease (years), and error bars were included to denote 1 std in the inference. Filters were applied to exclude inferences with large error margins (Methods) and were done completely agnostic of the ground truths. **a-b.** Many iterations of ground truths were simulated for a 34-year-old patient. In **a**, the model used to simulate ground truths did not incorporate feedback, while in **b** the model did. In both cases, the model used for the ABC inference did not incorporate feedback but was able to infer the parameters correctly within the statistical error. **c.** same as **a** and **d.** same as **b**, except the ground truth simulations and ABC inferences were carried out for a 63-year-old patient. In both **c** and **d**, the model used for the ABC inference also did not incorporate feedback but was able to infer the parameter values of *s* and the age of onset of disease within the statistical error. However, for **c** and **d**, *n* could not be inferred since there were not enough coalescent events in the later history to extract information about its dynamics after the initial expansion. For the inferred *n* vs true *n* plots in **c** and **d**, we decided to show them with no filter so the reader could see that the inferences contain no information.

**Supplemental Figure 26. Inference on patient data.** ABC was run on the patient data, and the model parameters were inferred. Joint distributions of the parameter values were plotted along with their marginals. *s* is in growth per year, and age of onset of the disease is in years. **a.** Distributions for inference on 34-year-old patient data. **b.** Distributions for inference on 63-year-old patient data. As observed, inference of *n* can indicate saturation when the number of cancer cells approaches *N*. In this case, the cancer expansion slows down and begins to exhibit neutral dynamics. This changes the coalescent structure, allowing ABC to detect the saturation and infer a saturation parameter value of $n = N$. When *N* is too large to affect the exponential growth dynamics of the cancer cells, ABC can only put a bound on *N*, namely that *N* must be larger than the number of cancer cells at the final time-point.

**Supplemental Figure 27. Identification of reliable sites to genotype single cells by detecting somatic mutations in the scRNAseq data**. We looked for sequencing reads supporting the mutated allele in the single-cell RNAseq data for all somatic point mutations detected in the WGS data from patient ET 1 and ET 2, including both mutations found in individual HSCs or shared by multiple HSCs, across 36 single-cell RNAseq data sets from bone marrow and peripheral blood samples of other donors. We assumed that the somatic mutations should not be observed in single-cell libraries that were not derived from the two patients. Thus, we estimated the false positive rate for each site as the number of molecules (identified using its unique UMI and cell barcode) in the control libraries supporting the mutated allele over the total number of molecules across all the controls mapping to that site. **a, b**. Fraction of UMIs in the scRNAseq data from ET 1 supporting the mutated allele vs the false positive rate estimated using the 36 control data sets as indicated above. Each dot corresponds to a somatic mutation, and its size is proportional to the number of UMIs mapping to the mutation site across the 36 control data sets (**a**), or in the scRNAseq library from ET 1 (**b**). **c, d**. Similar to (**a**) and (**b**) for patient ET 2. Blue dots indicate those mutations that were deemed reliable for mapping individual cells in the single-cell RNAseq library to the HSCs characterized by WGS for each patient.

**Supplemental Figure 28. *JAK2* mutated cells were identified in the scRNA-seq data sets using somatic mutations revealed by WGS. a.** Number of cells in scRNA-seq data for ET 1 and ET 2 with any transcript call (WT or mutant) from amplicon sequencing (specific gene names as labels) or 10X scRNA-seq reads ("somatic" label) of WGS-identified mutations. **b.** Fraction of cells with a *JAK2* amplicon mutant call out of all cells with a *JAK2* amplicon transcript observed and a WT (blue) or mutant (red) transcript observed from either another amplicon mutation or from a mutation identified from 10X scRNA-seq reads for mutations shared by all *JAK2* mutant HSCs ("somatic"). If all mutations are heterozygous and only a single *JAK2* amplicon transcript is observed per cell, we expect 50% of cells with a mutant non-*JAK2* transcript call to have a *JAK2* mutant transcript. **c**. Number of differentially expressed genes found comparing *JAK2* mutant to WT MEPs, using *JAK2* mutant/WT cells identified using a single amplicon mutation (*JAK2*, *UPF1*, or *NRROS*), mutations identified from 10X scRNA-seq reads for mutations shared by all *JAK2* mutant HSCs ("somatic"), or all mutation sources combined.

**Supplemental Figure 29. Gene ontology analysis of differentially expressed genes between *JAK2* mutant and WT cells. a-b.** Gene set enrichment analysis for KEGG 2019 Human biological processes (**a**) and ChEA/ENCODE transcription factor targets (**b**) of genes differentially expressed between *JAK2* WT and V617F MEPs and CD14+ cells.

**Supplemental Figure 30. Expression of specific genes and gene modules in *JAK2* mutant and WT cells. a.** Expression of specific genes in *JAK2* mutant (red) and WT (blue) HSCs. **b-e.** Gene set module scores for *JAK2* WT (blue) and *JAK2* mutant (red) HSCs and MEPs from each MPN patient. * indicates $p < 0.05$, Wilcoxon rank sum test. **f.** Total transcript counts per cell for *JAK2* WT (blue) and *JAK2* mutant (red) HSCs and MEPs from each MPN patient. * indicates $p < 0.05$, Wilcoxon rank sum test. **g**. Total transcript count per cell for each HSPC cell type from donor healthy 1. For all box plots, white center line is median, box limits are upper and lower quartiles, whiskers are 1.5x interquartile range, points are outliers.

**Supplemental Figure 31. Population balance analysis (PBA) of *JAK2*-mutant and WT HSPC differentiation dynamics. a.** UMAP showing cell types used for PBA in patient ET 1. **b.** Input net growth rate for individual HSCs, MEPs, and erythroid progenitors for each cell in patient ET 1. Inset bar plot shows the growth rate for *JAK2* WT (blue) and *JAK2* mutant (red) cells separately. **c-e.** Simulated *GATA1* expression levels over time for *JAK2* WT (blue) and mutant (red) HSPCs during erythroid differentiation, for ET V617L (**c**), other ET patients (**d**), and PV patients (**e**). The darker curves are the curves estimated using the true *JAK2* mutation status for each cell, while the lighter curves and shaded regions represent the mean and standard deviation from the null model with *JAK2* mutation status randomly assigned (25 permutations per condition).

**Supplemental Figure 32. Detection of somatic mutations in the scRNAseq data.** Example of a somatic point mutation (red arrow) detected only in the colony HSC-30 from patient ET 1. The mutation (G>A) could be phased with a nearby heterozygous SNP (black arrow). The seven read pileups correspond to a randomly chosen subset of sequencing reads from the single-cell RNAseq library where the mutation was detected.

**TABLES**

**Supplemental Table 1.** Primers and sequences for mutation-specific single-cell amplicon libraries (5′→3′).

**Supplemental Table 2**. The somatic single-nucleotide variants, indels, and microsatellites detected in the single-cell-derived WGS data from patients ET 1 and ET 2 that were used to reconstruct lineage trees are listed.

**Supplemental Table 3.** Differentially expressed genes in ET patients between *JAK2*-mutant and WT cells of different cell types. P-values for all ET patients with the *JAK2* V617F mutation are listed, along with the number of PV and ET patients in which the gene was differentially expressed and the *JAK2*-mutant vs WT average log2 fold change across all ET patients. All patients in which each gene is significantly differentially expressed are listed in the sig_pts column.

**Supplemental Table 4.** Differentially expressed genes in PV patients between *JAK2*-mutant and WT cells of different cell types. P-values for all PV patients with the *JAK2* V617F mutation are listed, along with the number of PV and ET patients in which the gene was differentially expressed and the *JAK2*-mutant vs WT average log2 fold change across all PV patients. All patients in which each gene is significantly differentially expressed are listed in the sig_pts column.

**Supplemental Table 5.** List of genes in each gene module.
