## Supplemental Text for "Reconstructing the lineage histories and differentiation trajectories of individual cancer cells in *JAK2*-mutant myeloproliferative neoplasms"

**Mathematical Description of Growth Dynamics of Mutated Cells**

**Outline.** We first define our mathematical model for mutated HSC growth dynamics, followed by the model with feedback where the fitness of the mutant cells decreases over time, and then describe an efficient way to simulate the mutant growth dynamics for inference. We then go on to derive analytical mathematical expressions for the average growth rate of mutant cells in our model, and use these to show that the fitness in terms of growth per year can be inferred from lineage trees without knowledge of the mutation rate.

**WF model with selection**

We chose to model mutated HSC growth (or the growth of a population of mutant cells within a population of wild-type cells) as a variation of the Wright-Fisher stochastic process [ Fisher (1922); Wright (1931) ] with selection included. We begin with an initial population of $N$ stem cells at $t = 0$. The stochastic process arises by iterating the following rule on the initial population: the cells at generation $t$ give birth to $N$ stem cells which correspond to generation $t+1$. Each cell in generation $t+1$ then selects a cell at random from generation $t$ as its parent, and the cells at generation $t$ die off. At generation $t'$, a mutation arises in one of the stem cells that gives it a selective advantage. From then on, the iterative rule changes in that instead of the cells from the parent generation being selected at random, each wild-type cell has probability p of being selected as a parent, and each mutant cell has probability $(1+s)p$ of being selected as a parent. The cells always inherit the phenotypic state of their parents (mutant or wild-type). After L iterations, we produce an evolving population of stem cells for $t = 0,1,\ldots,t',t'+1,\ldots,L$ along with a set of genealogical relationships.

Note that here there are a total of $L+1$ generations of cells, since the first generation corresponds to $t = 0$. In the methods section, we instead define $L$ to be the total number of generations. The way we have defined $L$ here is more convenient for the mathematical derivations, but it should be noted that any expression we have derived here in the supplement that uses $L$ will be replaced with $L-1$ in the methods.

**Computing the number of mutant cells as a function of time**

Here, we derive the distributions for the number of mutant cells as a function of time. Given the current generation of mutant cells, the number of mutant cells in the generation that follows is binomially distributed with parameter values that depend on the number of mutant cells in the current generation. This fact is used in a later section to compute the mean growth of mutant cells, and is also used in the Methods for efficient simulation of clonal expansions.

Suppose that there are $n$ mutant cells at generation $t$ (Note that in the Methods, we define $n$ to be the number of mutant cells at the final time-point. This $n$ is not the same). Since each wild-type cells has probability $p$ of being selected as a parent, and each mutant cell has probability $(1+s)p$ of being selected as a parent, then the probability that a cell at t + 1 choses a wild-type cell is $(N-n)p$, and the probability it chooses a mutant cell is (1 + s)np. Since probabilities must sum to one, p can be derived from the condition that $(N-n)p \ + (1+s)np \ = \ 1$, and so we obtain $p = \frac{1}{N+ns}$. After substituting, we obtain the probability a cell selects a mutant cell as its parent as:

$$\frac{(1+s)n}{N+ns}$$

Since each of the N cells at t + 1 either chooses a mutant cell or it does not, and since the choices are independent, then the number of mutant cells at generation t + 1 is binomially distributed with parameters $N$ and $\frac{(1+s)n}{N+ns}$. It then follows that we can compute the number of mutant cells as a function of time by beginning with an initial condition of $n = 1$ mutant cell , and then carrying out a series of binomial draws where we update $n$ before each draw to equal the current number of mutant cells.

**Wright-Fisher model with feedback**

The Wright-Fisher model is an idealized model that ignores a wide range of biologically plausible scenarios. For example, as the number of mutant cells increases, it is conceivable that there are underlying biological mechanisms that slow the growth of the mutant cells. To simulate such scenarios, we decided to incorporate feedback into the Wright-Fisher model. This is accomplished by letting the value of s change depending on the current number of mutant cells. In particular, if the number of mutant cells at generation $t$ is $n$, then the selection parameter at $t$ is $s(1-\frac{n-1}{N-1})^x$. Notice that when $x = 0$, the selection parameter remains constant and we recover the usual Wright-Fisher model with selection.

To simulate clonal expansions for mutant cells with feedback, we simply draw a series of numbers from binomial distributions as described in the previous section, except that instead of just updating $n$ before each draw, we first update $n$ and then $s(1-\frac{n-1}{N-1})^x$. Clonal expansions with feedback in the Methods section are simulated in this way.

**Dynamics of average population size**

Define n(t) as the number of mutant cells as a function of time. We now compute the expectation of $n(t)$, which we will call the mean trajectory. We consider the expected value of $n(t)$, conditioned on the mutant clone consisting of $n$ cells at time $t-1$. Upon conditioning, $n(t)$ reduces to a binomially distributed random variable as shown before, and so its mean is given by:

$$E_N[n(t) \mid n(t-1)=n] \;=\; N\frac{(1+s)\,n}{N+ns}$$

Subscript N is used to emphasize the dependence on population size. We then assume $N >> ns$, and drop subscript $N$to obtain:

$$E[\,n(t) \mid n(t-1)=n\,] \;=\; (1+s)n$$

Rewriting the expected value as a conditional expectation gives us

$$E[\,n(t) \mid n(t-1)\,] \;=\; (1+s)\,n(\,t-1\,)$$

We then take the expectation of both sides to generate the following recursion

$$E[n(t)] \;=\; (1+s)\,E[n(t-1)]$$

initial condition: $n(0)=1$

where without loss of generality we have let the time at which the mutation arrives be $t = 0$.

The recursion is then easily solved to obtain

$$E[n(t)] = (1+s)^t$$

**Dynamics of average population size conditioned on survival.**

When the number of mutant cells is small, they are susceptible to stochastic fluctuations and extinction. After growing to a sufficiently large size, their growth dynamics become deterministic and fluctuations can be safely ignored.

When using ABC to infer our model's parameters, we only consider trees where mutant cells have not gone extinct. We are thus interested in the growth dynamics conditioned on no stochastic extinction.

In the previous section, we computed the expectation value of the number of mutant cells as a function time across all trajectories. Here, we will constrain the expectation value to trajectories that do not go extinct. As would be expected, the average population size is larger when extinction is not allowed.

We begin by defining the conditioning event for our trajectories as $F = \{fixation\ will\ occur\}$ . Then we use Bayes' theorem to compute

$$E_N[\, n(t) \mid F \,] = \sum_{n=1}^{N} n\, P(\, n(t) = n \mid F\,) = \sum_{n=0}^{N} \frac{n\, P(\, n(t) = n\,)\, P(\, F \mid n(t) = n\,)}{P(\,F\,)}$$

The probability of fixation of a clone of size $n$ within a sufficiently large population of size $N$ and with fitness $1 + s$ is given by Kimura's diffusion approximation [ Kimura (1962)]

$$\frac{1 - e^{-2sn}}{1 - e^{-2sN}}$$

We therefore put

$$P(\, F \mid n(t) = n\,) = \frac{1 - e^{-2sn}}{1 - e^{-2sN}}$$

$$P(\, F\,) \; = \frac{1 - e^{-2s}}{1 - e^{-2sN}}$$

The probability of fixation $P(\, F\,)$ independent of the clone size is simply the probability that a clone of size $n = 1$ will eventually fix. Substituting both probabilities back into the sum and cancelling terms gives us

$$E_N[\, n(t) \mid F\,] \; = \frac{1}{1 - e^{-2s}} \sum_{n\,=\,0}^{N} \quad n\, P(\, n(t) = n\,)\,(1 - e^{-2sn})$$

Next we make a key biologically motivated assumption: we assume that t is sufficiently large so that the population of mutant cells is:

1) either large enough to exhibit deterministic dynamics, or
2) has gone extinct.

Therefore, $P(\, n(t) = n\,)$ vanishes except for when $n$ is large, or $n = 0$. Since the only nonzero terms in the expectation are those for large $n$, and since $1 - e^{-2sn} \sim 1$ when $n$ is large, then we may replace $1 - e^{-2sn}$ with $1$ in the expectation as an approximation. After replacing $1 - e^{-2sn}$ with 1 and observing that the sum is now the unconditional expectation, we obtain

$$E_N[\, n(t) \mid F\,] \; = \frac{1}{1 - e^{-2s}}\, E_N[\, n(t)\,]$$

If we let $N >> ns$ and drop the subscripts, the expectation on the right hand side becomes the mean trajectory $E[n(t)] = (1+s)^t$ that we derived in the previous section. Substituting then gives us

$$E[\, n(t) \mid F \,] = \frac{1}{1-e^{-2s}} (1+s)^t$$

Note that the above approximation is not valid for small values of $t$, for example, evaluating at $t = 0$ does not give an average population size of 1, because we assumed that $t$ must be sufficiently large.

Importantly, the above approximation has an interesting biological interpretation. The growth dynamics excluding extinction events is functionally equivalent to the growth dynamics of unconditional trajectories that begin with a clone size $\frac{1}{1-e^{-2s}}$ . Later, we will use this observation to show that s could be inferred without prior knowledge of mutation rate.

To validate the approximation derived above, we simulated the average growth dynamics of the mutant cells. Figures below show the simulated mean trajectory (blue dots), the simulated mean trajectories conditioned on no stochastic extinction (orange dots), and then our approximation of the mean trajectories conditioned on no stochastic extinction (blue curve) for different values of s. The simulated mean trajectories were collected by taking the average number of mutant cells at each time slice over a large number of simulated clonal expansions. The simulated mean trajectories conditioned on no stochastic extinction were generated similarly by first letting the simulated clones expand until they either went extinct or fixed, discarding the clones that went extinct, and then taking the average number of mutant cells at each time slice over the remaining clonal expansions. All simulated expansions were run with $N = 10^6$ and are shown for $g = 28$ generations.

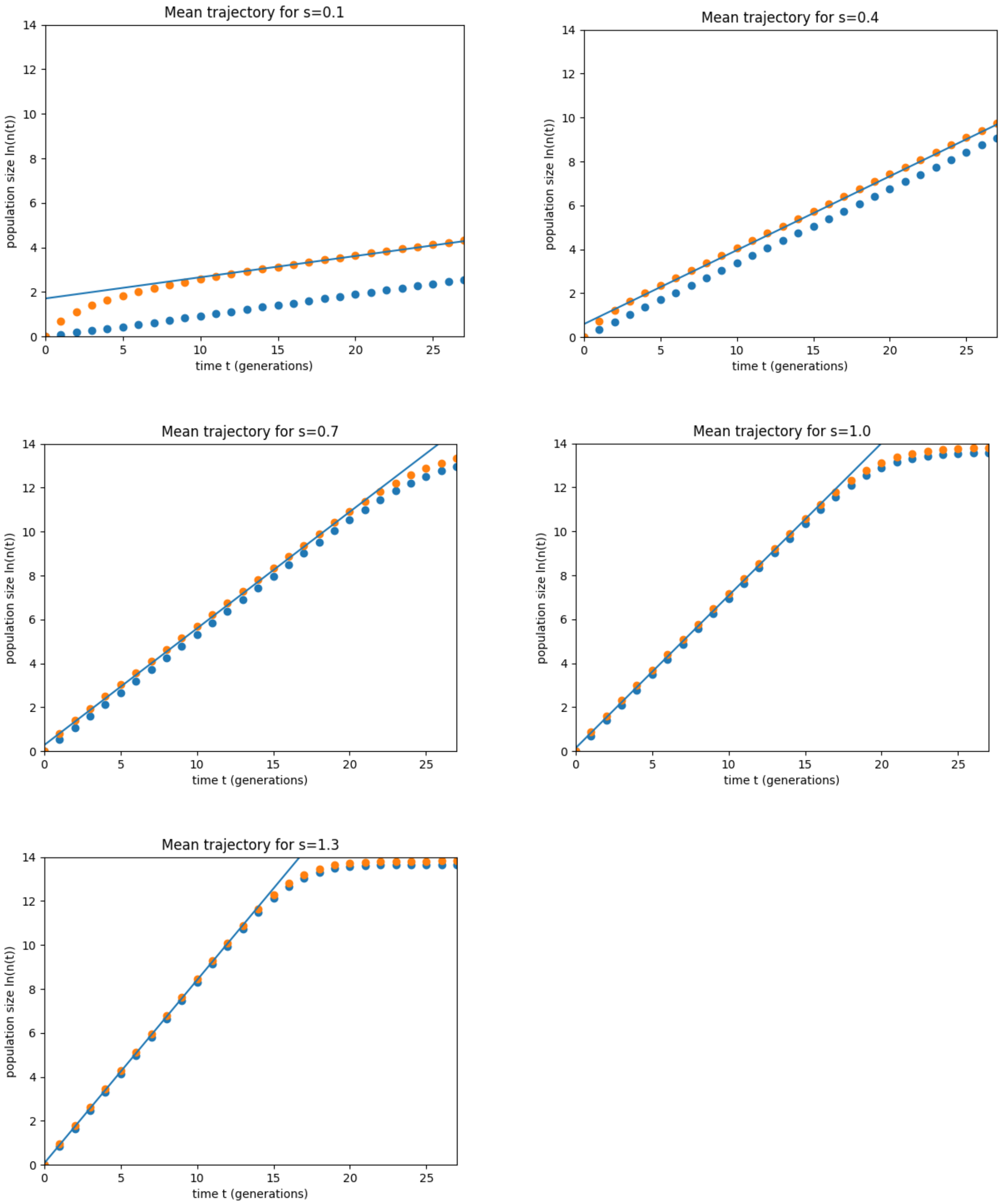


### Decoupling of fitness and mutation rate for weakly expanding clones

Under the neutral Wright-Fisher model, it is not possible to separately infer population size without knowledge of the mutation rate per generation. If we underestimate the mutation rate, we will overestimate the number of generations between coalescent events in the tree, and thereby overestimate the population size. Conversely, if we overestimate the mutation rate, we will underestimate the number of generations between coalescent events in the tree, and thereby underestimate the population size. It is therefore necessary to have a priori information about one parameter, for example mutation rate, to extract any information about the other, for example population size, from the tree.

Fitness, or growth rate per year $s_y$, can be inferred from the shape of the reconstructed lineage tree of a small number of cells randomly sampled from the population at the final time point. If $s_y$ is large, the population of mutated stem cells grows rapidly, therefore the coalescent events on the lineage tree will be confined to the first few generations, when the population size was small. Conversely, if $s_y$ is small, coalescent events are more likely to occur in the last few generations. Critically, unlike population size, $s_y$ can be inferred without any knowledge of the mutation rate, or equivalently, the total number of generations along the lineage tree. To intuitively understand this, note that rescaling the number of generations by a given factor scales the inferred population size by the same factor. Because at the onset of disease there is only one mutant cell, it might be expected that the growth rate, or s, must also be changed to achieve the scaled population size at the final time point. However, rescaling the number of generations also scales the minimum population size required before the mutated cells can escape stochastic extinction and grow exponentially. This is because more generations implies that the fitness advantage per generation is smaller and therefore the population is more susceptible to going extinct from random birth and death events. Taken together, these two competing effects precisely cancel and thus $s_y$ can be inferred directly from the observed lineage trees without knowledge of the mutation rate or the number of generations.

We will make the above intuition precise by deriving the analytical expression for coalescent statistics as a function of $s_y$ and showing that when expressed as growth per unit time, $s_y$ does not vary with the mutation rate.

**Mathematical analysis of mutation rate per generation and rate of population growth per unit time**

Here, we will show that it is possible to infer the population growth rate per unit time without knowledge of the mutation rate per generation.

First, we will define the population growth rate per unit time. Then, we will derive an expression for the expected coalescent times of a random sample of mutant cells, and use it to estimate the impact of mutation rate per generation on the inferred population growth rate.

Define L  to be the total number of divisions that an HSC would have undergone averaged across all HSCs, or equivalently the total number of generations in our trees. Note that knowing $L$ is equivalent to knowing the mutation rate per generation, since we know the number of mutations accrued throughout the patient's life.

**Definition of population growth rate per unit time**

Note that $1 + s$, where $s$ is the selection parameter, is the average growth per generation. We can also define a related quantity $s_y$ as the average percent growth per unit time (for example percent growth rate per year). If we let $a$ be the age of the patient, the number of generations per unit time is $\frac{L}{a}$ . Hence, per unit time, the mutant clone is expected to grow by a factor of $(1 + s)^{\frac{L}{a}}$, and so we arrive at the expression

$$s_y(s, L) = (1 + s)^{\frac{L}{a}} - 1$$

**Estimating the coalescent times**

We now derive an expression for the expected time for coalescence of k randomly sampled mutant cells given that the clone has expanded for g generations.

Let $t$ denote time in number of generations measured from the leaves of the tree towards the root. If there are $n(t)$ mutants at generation $t$, then the amount of coalescence time that passes from generation $t$ to $t + 1$is $\frac{1}{n(t)}$, where by coalescence time we are referring to the time-scale in the Kingman Coalescent model [ Kingman 1982a; Kingman 1982b; Kingman 1982c ]. To understand what we are doing intuitively, note that for the standard Wright-Fisher model without selection, where the population size $N$ is constant over time, the average time for coalescence of $k$ randomly sampled lineages is $\frac{N}{\binom{k}{2}}$generations. This is generally computed by scaling time so that $N$ generations correspond to 1 unit of time, and then letting $N$ become large. In doing so, the times of coalescence of $k$ randomly sampled lineages converge to the Kingman Coalescent where the coalescence times are known to be $\frac{1}{\binom{k}{2}}$. The coalescence times in generations can then be recovered through an inverse time-scale transformation. Note that this is equivalent to scaling time so that the time between two neighboring generations is $\frac{1}{N}$. To account for a variable population size, we let the time between two neighboring generations $t$ and $t+1$ be the inverse of the population size at $t$ and assume the population size is always large. In doing so, the times until coalescence also converge to the Kingman Coalescent model. The statistics of coalescence times are then recovered by transforming back to time in generations.

Since the expected coalescence time of k lineages is $\frac{1}{\binom{k}{2}}$ in units of coalescence, and since the population size in our simulations grows approximately as $n(t) = \frac{1}{1-e^{-2s}}(1+s)^t$, the expected coalescence time of k lineages in units of generations is the t satisfying:

$$\frac{1}{\binom{k}{2}} = \sum_{k=0}^{t-1} \frac{1}{\frac{1}{1-e^{-2s}}(1+s)^{g-k}}$$

We then notice the R.H.S. is a geometric sum and rewrite to obtain:

$$1 + \frac{s}{1-e^{-2s}} \frac{(1+s)^g}{\binom{k}{2}} = (1+s)^t$$

Solving for t we obtain:

$$t = \frac{log\left(1 + \frac{s}{1-e^{-2s}} \frac{(1+s)^g}{\binom{k}{2}}\right)}{log\,(1+s)}$$

We use $\frac{L}{a}$ to convert 3) from generational time to real time (such as years), thereby obtaining

$$t_k(s,g,L) = \frac{a}{L}\frac{log\left(1+\frac{s}{1-e^{-2s}}\frac{(1+s)^g}{\binom{k}{2}}\right)}{log\,(1+s)}$$

**Invariance theorem for weakly expanding clones**

We now show that for a weakly expanding clone, we always infer the correct percent growth rate per unit time (for example per year) independent of our assumption of $L$.

We begin by assuming that $s_p, g_p,$ and $L_p$ are the parameter values associated with our patient's tree, and that $|s_p| \ll 1$ so that selection is weak.

We then assume that we have incorrectly estimated our $g$ and $L$ parameters (i.e. our mutation rate) so that we erroneously believe they are $g_c = cg_p$ and $L_c = cL_p$ respectively. Note that we have kept the ratio $\frac{g_p}{L_p} = \frac{g_c}{L_c}$, and so we have treated the arrival time of the first mutated cell in real time as known.

Then we show that if we incorrectly assume our parameter values to be $g_c$ and $L_c$, we then infer $s_c = (1+s_p)^{\frac{1}{c}} - 1$ for our $s$ parameter, where

$$s_y(s_c, L_c) = s_y(s_p, L_p)$$

That is, we always infer the same percent growth per year. The way we show our inferred $s$ is $s_c$ is by plugging in $g_c$, $L_c$ and $s_c$ into the expected coalescent time expression we derived, and then showing that the coalescent times are identical to having plugged in $s_p, g_p,$ and $L_p$. In other words, we show that when we erroneously assume $g_c$ and $L_c$ are our parameter values, the $s$ value that generates trees that match our patient's is $s_c$, and that $s_c$ we infer combined with the $L_c$ we've assumed give us the same inference for yearly percent growth as the correct parameter values. We first show that $s_c$ is our inferred $s$:

Begin by recalling that our coalescent time expression is given by

$$t_k(s,g,L) = \frac{a}{L} \frac{log\left(1 + \frac{s}{1-e^{-2s}} \frac{(1+s)^g}{\binom{k}{2}}\right)}{log\,(1+s)}$$

Since $|s| \ll 1$, a Taylor expansion lets us make the following approximation:

$$1 - e^{-2s} \sim 2s$$

where we have let the 2nd order terms vanish. Substituting above and cancelling gives us

$$t_k(s,g,L) = \frac{a}{L} \frac{log\left(1 + \frac{1}{2} \frac{(1+s)^g}{\binom{k}{2}}\right)}{log\,(1+s)}$$

But then using $s_c = (1+s_p)^{\frac{1}{c}} - 1$ , $g_c = cg_p$ and $L_c = cL_p$ we can show:

$$t_k(s_c,g_c,L_c) = \frac{a}{L_c} \frac{log\left(1 + \frac{1}{2} \frac{(1+s_c)^{g_c}}{\binom{k}{2}}\right)}{log\,(1+s_c)} = \frac{a}{cL_p} \frac{log\left(1 + \frac{1}{2} \frac{\left(1 + \left[[(1+\ s_p)^{\frac{1}{c}} - 1\right]\right)^{cg_p}}{\binom{k}{2}}\right)}{log\,\left(1 +\ [(1+\ s_p)^{\frac{1}{c}} - 1]\right)}$$

$$= \frac{a}{L_p} \frac{log\left(1 + \frac{1}{2}\frac{\left(1+\ s_p\right)^{g_p}}{\left(\frac{k}{2}\right)}\right)}{log\left(1+\ s_p\right)} = t_k\left(s_p, g_p, L_p\right)$$

so that $s_c$ is our inferred $s$.

We then show our inference of percent growth is identical using $s_c = \left(1+s_p\right)^{\frac{1}{c}} - 1$ and $L_c = cL_p$:

$$s_y(s_c, L_c)\ = (1+s_c)^{\frac{L_c}{a}} - 1 = \left(1 + \left[\left[\left(1+s_p\right)^{\frac{1}{c}} - 1\right]\right)^{\frac{cL_p}{a}} - 1$$

$$= \left(1+s_p\right)^{\frac{L_p}{a}} - 1 =\ s_y\left(s_p, L_p\right)$$
