## Supplementary material for "Reconstructing the lineage histories and differentiation trajectories of individual cancer cells in *JAK2*-mutant myeloproliferative neoplasms": Methods

### Sample collection

Prospective MPN patients were identified through manual chart review of new outpatient clinic consults and clinician referral within our hospital network. Patients were required to have a confirmed diagnosis of PV or ET according to WHO criteria and next-generation sequencing (RapidHEME panel or SnapShot) documenting the presence of a missense variant at codon 617 of *JAK2*. Use of anti-platelet agents was permitted but disease-modifying treatments (e.g. hydroxyurea, interferon-alpha, ruxolitinib) were an exclusion criterion for the study. Consequently, our cohort consisted of newly diagnosed, treatment-naive *JAK2*-mutated MPN patients. Bone marrow aspirate samples were uniformly collected at the time of diagnostic bone marrow biopsy under tissue banking protocols at the participating centers. The study was approved by and conducted in accordance with the Declaration of Helsinki protocol (Dana-Farber Cancer Institute IRB protocol no. 01-206: Tissue and Data Collection for Research Studies in Patients with Hematologic Malignancies, Bone Marrow Disorders, and Normal Donors, and Massachusetts General Hospital protocol 13-583). All patients provided informed consent. The healthy donor samples were purchased as de-identified samples from the Boston Children's Hospital. Healthy donor 1 was a 22-year-old female and healthy donor 2 was a 29-year-old female. Bone marrow biopsies were performed on all donors. RapidHEME panel screen on peripheral blood from the same patients was performed as a part of the clinical diagnosis to screen for *JAK2* mutations and potential secondary mutations. Approximately 10-20mL of bone marrow aspirate from each donor was collected in EDTA-coated tubes. The syringes and tubes used were sterile with no preservative-free heparin coating. The bone marrow aspirates (BMA) were kept at room temperature until use.

### Mononuclear cell isolation

Mononuclear cells (MNCs) were isolated from the BMA via a density gradient centrifugation protocol using Stemcell Technologies's SepMate system. BMA, phosphate-buffered saline (PBS) with 2% fetal bovine serum (PBS + 2% FBS; Stemcell Technologies #07905), Lymphoprep (Stemcell Technologies #07801), and the centrifuge (Eppendorf #5810R) were all acclimated at room temperature. Approximately 20-22mL Lymphoprep was added to the 50mL SepMate tube (Stemcell Technologies #15450) by carefully pipetting it through the central hole of the SepMate insert, ensuring that as few air bubbles as possible were present. BMA was diluted with an equal volume of PBS + 2% FBS and mixed gently with wide-bore pipette. Keeping the SepMate tube vertical, the diluted sample was added by slowly pipetting it down the side of the tube. The diluted BMW was centrifuged at 1200 x g for 20 minutes at room temperature, with the brake off. For BMA rich in platelets/plasma, the top layer of platelets/plasma was pipetted off. The remaining volume down to the SepMate central hole (containing all the enriched MNCs) was poured into a new 50mL tube. The MNCs were washed by topping up until 45mL with PBS + 2% FBS and mixing well with a wide-bore pipette. The MNCs were then centrifuged at 300 x g for 12 minutes at room temperature, with the brake low, and the supernatant was removed. The MNC pellet was topped up again with PBS + 2% FBS, and the volume was mixed well with a wide-bore pipette. Centrifuge at 120 x g for 12 minutes at room temperature, with the brake off. The supernatant was then removed. For BMA rich in platelets, an additional wash and centrifugation at 12 minutes at room temperature, with the break off was performed. The MNC pellet was resuspended in 1mL of EasySep buffer (Stemcell Technologies #20144) and mixed with a wide-bore pipette. Cell concentration was counted with a hemocytometer (Reichert) and a Tali Image Cytometer system (ThermoFisher #T10796) using Tali Image Analysis Slides (ThermoFisher #T10794). Cells were placed on ice until further use.

**CD34+ enrichment**

CD34+ MNCs were isolated using the protocol for EasySep Human CD34 Positive Selection Kit II (Stemcell Technologies #17856). MNCs (at concentration of >$10^8$ cells/mL EasySep buffer) were added to 5mL (12 x 75 mm) polystyrene round-bottom tube (Stemcell Technologies

#38007). EasySep Human CD34 Positive Selection Cocktail (Stemcell Technologies #17856C) was added at a concentration of 100 µL per 1mL of sample. The sample was then mixed and incubated at room temperature for 10 minutes. EasySep Dextran RapidSpheres (Stemcell Technologies #50100) were vortexed for 30 seconds. RapidSpheres were added at a concentration of 75 µL per 1 mL of sample. The sample was mixed and incubated at room temperature for 5 minutes. The tube was topped up to 2.5mL with EasySep buffer and gently mixed. The tube was placed in EasySep magnet (Stemcell Technologies #18000) and incubated at room temperature for 3 minutes. The supernatant was discarded by inverting the magnet with the tube inside. This process was repeated 4 more times for a total of 5 rounds of enrichment. Cells were resuspended in PBS+2% FBS after the last round of enrichment. Cell concentration was counted with a hemocytometer (Reichert #1492) and Tali Image Cytometer. Cells were placed on ice until further use. CD34+ MNCs were used to create single-cell cDNA, single-cell RNA-Seq (scRNA-seq) and *JAK2* amplicon libraries as described below.

**Single-cell cDNA libraries**

The isolated CD34+ MNC suspensions were used to generate single-cell gel bead emulsions (GEMs) using a 10x Genomics Chromium controller (10x Genomics #120223). Following steps 1 and 2 of the protocol "Chromium Single Cell 3' Reagent Kit v3" (10x Genomics, CG000183 Rev A), single-cell cDNA libraries were constructed using the Chromium Single Cell 3' Library & Gel Bead Kit v3 (10x Genomics #1000075). The protocol yields 40µL of cDNA per sample after step 2.4. scRNA-Seq libraries were generated from 10µL of single-cell cDNA libraries using Step 3 of the Chromium Single Cell 3' Reagent Kit v3 user guide (10x Genomics, CG000183 Rev A).

**Locus-specific single-cell amplicon libraries**

We developed the following protocol to preferentially amplify transcripts containing loci-of-interest from single-cell cDNA libraries (ex. *JAK2*-V617F), thereby generating locus-specific single-cell amplicon libraries (SI Fig. 1a-c).

A triple-nested PCR approach was used to amplify the transcripts carrying the loci-of-interest from single-cell cDNA libraries with high sensitivity and specificity (Fig. 6a, SI Table 1). The approach used locus-specific reverse primers that flank the mutation site combined with generic forward primers that preserves the single-cell barcoding structure. In total, there were 5 total PCR steps (3 nested steps that increasingly filtered for a specific transcript, 1 step that added a Read2 sequence, and 1 step that added an Illumina P7 adaptor sequence). In Step 1, a PCR was conducted using a forward primer containing both the Illumina P5 sequence and part of the Read 1 sequence and a reverse primer containing a locus-specific sequence approximately ~300bp upstream of the mutation site. In Step 2, a PCR was conducted using a shortened version of the forward primer in Step 1 and a reverse primer containing a locus-specific sequence approximately ~150bp upstream of the mutation site. In Step 3, a PCR was conducted using the forward primer from Step 2 and a reverse primer containing a locus-specific sequence approximately ~50bp upstream of the mutation site. In Step 4, a PCR was conducted using the forward primer from Step 2 and a reverse primer containing part of the locus-specific sequence used in Step 3 combined with a Read2 sequence. In Step 5, a PCR was conducted using the forward primer used in Step 1 and a reverse primer containing the Read2 sequence combined with the Illumina P7 sequence.

The following was the specific protocol used for constructing the *JAK2*-V617F amplicon libraries. All PCR's were conducted in TempAssure PCR tubes (USA Scientific #14024700) on a Bio-Rad C1000 Touch Thermal Cycler.

In Step 1, a 25uL PCR mixture was made containing 12.5uL of Amplification Master Mix (10x Genomics #220125), 1.25uL of cDNA additive (10x Genomics #220067), 1.25uL of forward primer (P5-Partial Read 1, AATGATACGGCGACCACCGAGATCTACACTCTTTCCCTACACGACGCTC) at 20uM, 1.25ul of reverse primer (Reverse Ext 1, ACCAACCTCACCAACATTACAGAGGCCT) at 10uM, 3ng of cDNA library material, and remaining volume with nuclease-free water. With the thermal cycler lid set to 105℃, the following thermal cycling protocol is used: initial denaturation at 98℃ for 45 seconds, 10 cycles of denaturing at 98℃ for 20 seconds, annealing at 67℃ for 30 seconds, extension at 72℃ for 180 seconds, and a final extension at 72℃ for 60 seconds.

The PCR reaction mixture was purified using SPRIselect (Beckman Coulter #B23318) as follows. 20uL (i.e. 0.8x) of SPRIselect reagent was added to the reaction mixture, pipette mixed, and incubated at room temperature for 5 minutes. The PCR tube was placed on the 10x Magnetic Separator (10x Genomics #230003) on High until solution clears (usually about a minute). Supernatant was removed and discarded. 200uL of 80% ethanol in nuclease-free water was added to the pellet and allowed to sit for 30 seconds. The ethanol wash was removed and repeated once more. The PCR tube was briefly centrifuged and put back into the magnet at Low setting. Any remaining ethanol wash was removed, and the pellet was allowed to air dry for 1 minute. The DNA was then eluted by removing the PCR tube from the magnet, pipetting 20uL Buffer EB (Qiagen #19086) onto the pellet, pipette mixing, allowing the mixture to equilibrate for 2 minutes, placing the tube on the magnet on Low, and then eluting the liquid into a new tube.

In Step 2, a 25uL PCR mixture was made containing 12.5uL of Amplification Master Mix (10x Genomics #220125), 1.25uL of cDNA additive (10x Genomics #220067), 1.25uL of forward primer (Partial P5, AATGATACGGCGACCACCGAGATCT) at 20uM, 1.25ul of reverse primer (Reverse Ext 2, AGGAGACTACGGTCAACTGCATGAAACAGA) at 10uM, 5uL of the DNA product from

Step 1, and 3.75uL of nuclease-free water. With the thermal cycler lid set to 105℃, the following thermal cycling protocol is used: initial denaturation at 98℃ for 45 seconds, 10 cycles of denaturing at 98℃ for 20 seconds, annealing at 67℃ for 30 seconds, extension at 72℃ for 180 seconds, and a final extension at 72℃ for 60 seconds. The PCR reaction mixture was purified using SPRIselect (0.8x) as described previously.

In Step 3, a 25uL PCR mixture was made containing 12.5uL of Hot Start Taq 2x Master Mix (NEB #M0496S), 1.25uL of cDNA additive (10x Genomics #220067), 1.25uL of forward primer (Partial P5, AATGATACGGCGACCACCGAGATCT) at 20uM, 1.25ul of reverse primer (Reverse Ext 3, GCAGCAAGTATGATGAGCAAGCTTTCTCACA) at 10uM, 5uL of the DNA product from Step 2, and 3.75uL of nuclease-free water. With the thermal cycler lid set to 105℃, the following thermal cycling protocol is used: initial denaturation at 95℃ for 30 seconds, 10 cycles of denaturing at 95℃ for 30 seconds, annealing at 60℃ for 60 seconds, extension at 68℃ for 210 seconds, and a final extension at 68℃ for 300 seconds. The PCR reaction mixture was purified using SPRIselect (0.8x) as described previously.

In Step 4, a 25uL PCR mixture was made containing 12.5uL of Amplification Master Mix (10x Genomics #220125), 1.25uL of cDNA additive (10x Genomics #220067), 1.25uL of forward primer (Partial P5, AATGATACGGCGACCACCGAGATCT) at 20uM, 1.25ul of reverse primer (Partial Reverse Ext 3-Read 2, GTGACTGGAGTTCAGACGTGTGCTCTTCCGATCTAGCAGCAAGTATGATGAGCA) at 10uM, 2uL of the DNA product from Step 3, and 6.75uL of nuclease-free water. With the thermal cycler lid set to 105℃, the following thermal cycling protocol is used: initial denaturation at 98℃ for 45 seconds, 10 cycles of denaturing at 98℃ for 20 seconds, annealing at 67℃ for 30 seconds, extension at 72℃ for 195 seconds, and a final extension at 72℃ for 60 seconds. The PCR reaction mixture was purified using SPRIselect (0.8x) as described previously.

In Step 5, a 25uL PCR mixture was made containing 12.5uL of Amplification Master Mix (10x Genomics #220125), 0.5uL of SI-PCR primer (10x Genomics #220111), 2.5ul of from one well (recorded for future reference) from the Chromium i7 Sample Index Plate (10x Genomics #220103), 2uL of the DNA product from Step 4, and 7.5uL of nuclease-free water. With the thermal cycler lid set to 105℃, the following thermal cycling protocol is used: initial denaturation at 98℃ for 45 seconds, 10 cycles of denaturing at 98℃ for 20 seconds, annealing at 54℃ for 30 seconds, extension at 72℃ for 195 seconds, and a final extension at 72℃ for 60 seconds. The PCR reaction mixture was purified using SPRIselect (0.8x) as described previously.

Quality control of each step was verified using High Sensitivity D5000 ScreenTape (Agilent #5067-5592) and High Sensitivity D5000 ScreenTape (Agilent #5067-5593) on an Agilent 2200 Tapestation system (SI Fig. 1e). Amplicon libraries display multiple peaks in the TapeStation trace (SI Fig. 1e), which is believed to be due to promiscuous binding of primers to poly-A like regions. Successful enrichment with correct DNA product were typically associated with “sawtooth”-like traces. After using different indexing primers in Step 5 for different libraries, several libraries were pooled and subsequently sequenced together on an Illumina Novaseq 6000 sequencing machine. The sequencing cycle settings were as follows: 28 cycles for Read 1, 8 cycles for the i7 index, and 91 cycles for Read 2.

cDNA material generated in the single-cell cDNA library construction step described in the previous section was generally adequate for making the amplicon libraries described here. If additional cDNA was needed, the single-cell cDNA library was amplified by repeating Step 2.2 of the Chromium Single Cell 3’ Reagent Kit v3 user guide (10x Genomics, CG000183 Rev A).

Depending on the mutation being targeted, optimization of the location of the locus-specific primers and their biochemical properties were needed to minimize non-specific primer binding. For the *JAK2*-V617F mutation, the three locus-specific primers used were ~30bp and have melting temperatures of around 68 ℃. Since the PCR cycle number is dependent on the expression level of the gene harboring the targeted mutations, the number of cycles was optimized experimentally for other mutation targets (ex. *TET2*).

**Stem cell genotyping and preparation for WGS**

Several types of cells were genotyped and prepared for WGS: (1) HSCs, MPPs, and stromal cells from bone marrow biopsies, and (2) fibroblasts from a skin biopsy.

1. HSCs, MPPs, and stromal cells

10mL of bone marrow aspirates were used to isolate HSCs, MPPs, and stromal cells. Erythrocytes were removed from 9mL of bone marrow aspirate samples using red blood cell lysis buffer. CD34-enrichment was performed using magnetic-assisted cell sorting with anti-CD34 magnetic beads (Miltenyi Biotech #130-046-703). Different cell populations were purified through using FACSAria (Becton Dickinson). The following combinations of cell surface markers were used to define cell populations. HSC: CD34+CD38-CD45RA-CD90+CD49f+; MPP: CD34+CD38-CD45RA-CD90-CD49f-.

Cells were first sorted into a collection tube, and a second index sorting step was performed to seed single-cells into round bottom 384-well plates, with ~90 µL of growth medium. Colonies were grown in StemSpan SFEM medium (Stemcell Technologies #09650) supplemented with: SCF (100 ng/ml), Flt3-L (100 ng/mL), TPO (50 ng/mL), IL3 (10 ng/mL), Tpo (50 ng/mL), Epo (10 ng/mL), and GM-CSF (10 ng/mL). Colonies were grown at 37°C in 5% $CO_2$ for 4-6 weeks before collection, with partial media exchange and fresh cytokine repletion every 2 weeks.

Polyclonal mesenchymal stem cells (MSCs) cultures were established from 1mL of whole bone marrow aspirate samples after red blood cell lysis, cells were plated in tissue culture treated dishes in DMEM-F12 medium (GIBCO #11320082), supplemented with 10% fetal bovine serum (VWR # 89510-186). MSCs were kept in culture for a week and medium was replaced each day to remove non-adherent cells. Stromal cells were ready for collection after reaching 80-100% confluency.

2. Skin biopsy

One skin biopsy sample was obtained from the *JAK2* V617L-mutant patient. The tissue was dissociated using collagenase I (Stemcell Technologies #07415) and genomic DNA was extracted as described below.

For both samples, genomic DNA was extracted from cells using QIAmp UCP DNA Micro Kits (Qiagen #56204) and eluted into a final volume of 30 µL. The DNA concentration was quantified using a Qubit fluorometer (Invitrogen #Q32866) and Qubit dsDNA HS Assay Kit (Invitrogen #32854). Some of the genomic DNA (1 ng) was amplified using *JAK2*-V617F specific primers and screened for the mutation using Sanger sequencing.

To generate PCR products for the *JAK2* target loci for Sanger sequencing, we performed three rounds of nested PCR with locus-specific reverse primers and generic forward primers. First, a 611bp fragment of *JAK2* was amplified to obtain a sufficient amount of DNA containing the mutation site. PCR components included 1ng of gDNA template, Phusion Hot Start Flex 2x Master Mix (M0536L), forward and reverse primers ACTCTTGCTCTCTCTCACTTTG and ACCTGCCATAATCTCTTTTGCT (DNA oligos synthesized by IDT), respectively, and nuclease-free water. The amplification protocol was as follows: (1) initial denaturation at 98°C for 30 seconds; (2) 40 cycles of denaturation at 98°C for 10 seconds; (3) annealing at 63°C for 30 seconds; (4) extension at 72°C for 30 seconds; and (5) final extension at 72°C for 10 minutes.

PCR reaction mixtures were slowly run on a 1.5% agarose gel in EDTA. PCR products were extracted from the gel using a Monarch DNA Gel Extraction Kit (NEB T1020L).

Second, the PCR products were Sanger sequenced in four separate reactions (each with one of four primers) through the Psomagen's gDNA sequencing service. The four primers used were flanking both directions of the mutation site (DNA oligos synthesized by IDT):

1. TGGCAGAGAGAATTTTCTGAAC (147bp upstream of the mutation)
2. ACTCTTGCTCTCTCTCACTTTG (304bp upstream of the mutation)
3. GTCCTACAGTGTTTTCAGTTTCA (166bp downstream of the mutation)
4. ACCTGCCATAATCTCTTTTGCT (306bp downstream of the mutation)

Electropherograms from Sanger sequencing were annotated. A sample of cells was marked as likely having the mutation if all of the corresponding electropherograms for that sample contained the mutation. Samples were then selected and their gDNA submitted for WGS sequencing (Broad Genomics). The WGS sequencing results were always consistent with the Sanger sequencing, that is the *JAK2* mutation was detected in the whole-genome sequencing data for the colonies that were designated as mutated using Sanger sequencing.

Of the ~600 MPPs and ~600 HSCs cultured from ET 1, we recovered sufficient genomic DNA from 62 MPP colonies and 22 HSC colonies after expansion. Of these, 6 MPP colonies and 16 HSC colonies had the *JAK2*-V617F mutation, indicating that mutated HSCs proliferated more in our culture conditions compared with WT HSCs. A similar bias was also observed in cells cultured from the patient ET 2. The total sequencing depth was 1.6 billion reads for ET 1 and 480 million reads for ET 2.

**scRNA-seq preprocessing and cell type identification**

scRNAseq libraries and the amplicon libraries were sequenced on the NovaSeq platform. The resulting bcl files were run through the Cell Ranger 4.0.0 pipeline to generate the fastq files and the count matrices. The fastq files for the amplicon libraries were analyzed as described below. Count matrices from each patient were loaded into Scanpy[1]. Genes expressed in < 3 cells and cells with < 2,000 total UMIs or > 20% mitochondrial transcripts were excluded from further analysis. Total count normalization was performed so that each cell had 100,000 total transcripts. Log-transformed expression values were used for UMAP visualization and clustering after regressing out % of mitochondrial transcripts and total counts. UMAP coordinates were calculated using Scanpy default parameter values. To assign an HSPC cell type to each cell, the scRNA-seq data from all patients were merged and batch corrected using Seurat's data integration workflow with the default parameter values[2]. Louvain clustering was performed on the merged and batch-corrected dataset in Scanpy and each cluster was assigned an HSPC cell type identity by manually reviewing the expression levels of marker genes in that cluster. Identification of monocyte subsets was performed similarly on CD14+ cells from all donors.

**Differential gene expression analysis between HSCs from different patient groups**

Scanpy's implementation of the Student's *t*-test was used to compare gene expression between ET, PV, and healthy HSCs using total count normalized gene expression values without batch correction. To limit the impact of batch effects, we performed all pairwise comparisons between each patient in both groups (e.g., for the ET vs PV comparison, we separately compared ET 1 and PV 1, ET 1 and PV 2, etc) and identified genes that were differentially expressed in all comparisons for gene set enrichment analysis using GSEApy (https://pypi.org/project/gseapy/).

**Identification of *JAK2*-mutant cells in the scRNA-seq data**

To identify individual cells in the scRNAseq library as either WT or *JAK2*-mutated, we separately analyzed the fastq files of the amplicon libraries derived from the same cells. First, the reads in the fastq files were discarded if the average Illumina base quality value was less than 30. Next, only reads were retained whose single-cell barcode uniquely matched (up to at most 2 bps differences) a barcode from the list of single-cell barcodes in the scRNAseq library of the same cells. A threshold for the number of reads was determined by inspecting the plot of the number of reads from each molecule (unique cell-barcode and UMI) after rank ordering the molecules by their number of reads, corresponding to the knee in the plot, usually around 100 to 1,000 reads depending on the sequencing depth (SI Fig. 1f). Molecules that had fewer reads than the threshold were discarded. To correct for sequencing errors in the UMIs, those molecules that shared the same barcode but had the same UMI sequence up to 2 mismatches were merged. Next, the mutation site was inspected in the remaining reads. Only reads were retained that had the expected WT nucleotide, or the expected mutated version of the nucleotide, and the where the 10 bps upstream and downstream of the mutation site matched the reference genome. A molecule was designated as "mutated" if more than half of its reads carried the mutated nucleotide, and WT otherwise. The above analysis pipeline was implemented in Matlab R2018.

Cells with at least one WT amplicon call were marked as "WT" for differential expression analysis. It is important to note that in cells with a heterozygous *JAK2* mutation, the presence of a WT transcript does not guarantee that the cell is homozygous WT. However, cells with at least one *JAK2*-mutant transcript were definitively classified as *JAK2*-mutant cells. Therefore, to correct for *JAK2* mutation heterozygosity in patients with < 50% peripheral blood *JAK2* mutation VAF, the fraction of *JAK2*-mutant cells in a cell population was estimated as the fraction of *JAK2*-mutant molecules in the cell population multiplied by 2. This correction factor comes from the observation that, since most cells only have one *JAK2* transcript call, cells with a heterozygous *JAK2* mutation have approximately a 50% chance of having a *JAK2*-mutant transcript sequenced so approximately half of true *JAK2*-mutant cells have a mutant *JAK2* transcript sequenced.

**Differential gene expression analysis between *JAK2*-mutant and WT cells**

Scanpy's implementation of the Wilcoxon rank-sum test was used to compare gene expression between *JAK2*-mutant and *JAK2*-WT cells within individual patients using total count normalized gene expression values. Different normalization strategies, including total count normalization without highly expressed genes and scTransform normalization[3] (ref) yielded similar results (not shown). Comparisons were done between *JAK2*-WT and *JAK2*-mutant cells in each cell type separately for each patient. A cell was classified as *JAK2*-mutant if it had at least one *JAK2*-mutant transcript. In ET 1 or ET 2, additional cells were classified as JAK2-mutant if they had at least one mutant transcript with a mutation specific to the *JAK2*-mutant HSCs. To find disease subtype specific differentially expressed genes, Benjamini-Hochberg-corrected *P* values for each gene from all patients with ET or PV were combined using Fisher's method. All genes identified as significant in the combined ET or PV comparisons are listed in SI Tables 3-4. Gene set enrichment analysis was done using GSEApy to find enriched KEGG biological processes and ChEA/ENCODE transcription factor target groups.

Gene module scores were computed by summing total-count normalized expression levels for all genes in a gene set for each cell and taking the $\log_2$ of this sum. Gene set lists for each module are found in SI Table 5. Two-tailed Student's t-tests were performed to compare module scores between *JAK2*-mutant and WT cells in each patient and corrected for multiple hypotheses using the Benjamini-Hochberg procedure.

**Population balance analysis of erythroid differentiation**

PBA is an algorithm to infer single-cell differentiation dynamics from static scRNA-seq datasets[4]. We applied it to our CD34+ HSPC scRNA-seq data to compare the rate of erythroid fate commitment in *JAK2*-mutant and *JAK2*-WT cells. HSCs, MEPs, and erythroid progenitors identified as *JAK2*-mutant or *JAK2*-WT using either amplicon-sequenced mutations or scRNA-seq somatic mutation calls were selected. In patients with heterozygous *JAK2* mutations, some cells with only *JAK2*-WT transcripts sequenced are actually *JAK2*-mutant cells which by chance did not have any observed *JAK2*-mutant transcripts. Therefore, to correct for *JAK2* mutation heterozygosity, in patients with < 50% peripheral blood *JAK2* mutation VAF some cells with only *JAK2*-WT transcripts observed were added to the *JAK2*-mutant population with probability

$p(x_i \text{ is } JAK2\text{-mutant} \mid w) = ((0.5)^w * z)/ ((1 - z) + z * (0.5)^w)$.

This is the Bayesian probability that a cell $x_j$ of cell type j with *w* WT transcripts observed is in truth a heterozygous *JAK2*-mutant cell, with the prior probability *z* equal to the estimated fraction of *JAK2*-mutant cells of cell type j.

Separate kNN (*k*=10) graphs were constructed for *JAK2*-mutant and WT cells for each dataset. The kNN graph and the net growth rate for each cell (*R*) were used as inputs to the PBA algorithm. R values for all cells in each dataset must sum to 0 for the population to be at steady state. R for each cell was defined as follows:

1. Cells with *HBB* expression in the 90th percentile or higher out of all HSCs, MEPs, and erythroid progenitors were assigned *R* = -1. This population represents the most differentiated erythroid progenitors.
2. The remaining MEPs and erythroid progenitors were assigned *R* = 0.9 * (number of terminal erythroid progenitors) * (number of intermediate MEPs and erythroid progenitors). Therefore, the total net growth of MEPs and intermediate erythroid progenitors balances out 90% of the total differentiation flux out of the terminal erythroid compartment.
3. Each HSC was assigned *R* = (number of terminal erythroid progenitors)/(number of HSCs) * 0.1 to balance differentiation of the terminal erythroid progenitors.

The net growth rate of the entire population (HSCs, MEPs, and erythroid progenitors combined) is therefore 0.

Using these kNN and *R* values as input, we used PBA to compute the differentiation potential of each cell. As in Weinreb *et al*[4], this potential was used to estimate transition rates between every pair of sampled cell states in the dataset, using *D* = 1 as the PBA diffusion parameter value. The discrete time Markov process defined by these transition rates describing HSPC differentiation was simulated, with the cell with the highest differentiation potential (least differentiated) as the starting state. At each timestep of the simulation, the probability of the initial cell occupying each sampled cell state $x_i$ is given by $x = (A^t)y$, where $y_i$ is 1 at the initial cell index, 0 elsewhere. This probability distribution over cell states at each timestep is used to compute the average *GATA1* expression level at each timestep as $x \bullet g$, where $g$ is the *GATA1* expression level for each cell. Erythroid progenitors with $R = -1$ were allowed to transition into a terminally differentiated erythroid state with *GATA1* expression equal to the mean erythroid progenitor *GATA1* level, representing differentiation out of the sampled population.

**Whole-genome sequencing data analysis**

Raw sequencing reads were mapped to the GRCh38 build of the human reference genome using BWA-MEM[5] version 0.7.17-r1188. Aligned reads in BAM format were processed following the Genome Analysis Toolkit (GATK, version 4.1.2.0) Best Practices workflow to remove duplicates and recalibrate base quality scores[6].

**Detection of somatic single-nucleotide variants and INDELs**

The germline short variant discovery workflow from GATK version 4.1.2.0 was used to detect somatic single-nucleotide variants (SNVs) and small insertions and deletions (INDELs) in the single-cell-derived WGS data. In brief, intermediate GVCF files were generated for each colony and chromosome using HaplotypeCaller in GVCF mode. Default parameter values were used except for the output-mode argument, which was set to "EMIT_ALL_SITES". Next, GVCF files for all colonies from each patient were consolidated into a single GVCF file using the GATK functionality CombineGVCFs using default options. Finally, colonies were jointly genotyped across all sites using GenotypeGVCFs with the "--include-non-variant-site" parameter set to true.

In order to identify somatically acquired point mutations and indels in the colonies the following steps were followed.

- All sites with a genotype quality of at least 50 and showing variation in at least one colony were selected.
- Variants mapping less than 10bp upstream or downstream of a simple repeat reported in the RepeatMasker track from the UCSC Genome Browser were discarded.
- Variants mapping less than 100bp apart from each other were removed, as in our experience these are likely artefacts.
- Variants that could not be genotyped in 10 or more colonies in each patient were discarded.
- To remove subclonal mutations acquired during *in vitro* culture the mean variant allele frequency (VAF) value across all mutated colonies was required to be between 0.3 and 0.7 for patient ET 2 (female) and for the autosomes in the case of patient ET 1 (male). Additionally, we required a minimum coverage of at least 6 sequencing reads. Variants mapping to chromosomes X and Y in the case of patient ET 1 and chromosome X in colony MPP-73 from patient ET 2, which harbors only one copy of this chromosome (SI Fig. 12), were required to show a VAF value of at least 0.9 and the coverage threshold was set to 3 sequencing reads.
- Sites supporting more than 4 genotypes across all colonies were removed, as after manual inspection of a number of such cases we concluded that these were likely artefacts.
- We required the genotype quality in the bulk sequencing data from stromal cells to be at least 80 in order to remove variants in low-quality mapping regions.
- Only variants with a homozygous reference genotype in the bulk sample were kept in order to filter out germline heterozygous polymorphisms.
- Given that all cancer cells share 220 and 398 mutations in patients ET 1 and ET 2, respectively, we reasoned that any mutations occurring early in development and giving rise to both the cancer and wild type cells should be present in all cancer colonies and in a subset of the normal colonies, but not in normal colonies and just a subset of the cancer colonies. Therefore, all mutations detected in just a subset of the cancer cells and one or more wild-type colonies were

discarded, as these are likely germline polymorphisms or artefacts. We did not find any mutation present in all cancer cells and one or more normal cells. In this analysis we only focused on the discovery of heterozygous variants given that all colonies, with the exception of the loss of chromosome X in colony MPP-73 from ET 2, show diploid karyotypes with no copy number alterations.

All variants remaining after applying the filters described above were visually inspected using BAMsnap (https://github.com/parklab/bamsnap), and those deemed to be false positives were removed. The remaining variants were deemed to be somatic and were considered for further analysis. Annovar (version 2018Apr16) was used to annotate variants. Missense variants predicted to be deleterious by MetaLR and MetaSVM were considered pathogenic[7].

**Detection of microsatellite mutations**

Somatic mutations at microsatellite loci were detected using HipSTR version 0.6.2[8] using *de novo* stutter estimation and allele generation, and the reference set of microsatellite loci provided by the authors. Subsequently, microsatellite calls were filtered and only calls satisfying the following criteria were considered for further analysis: (1) Posterior probability for the genotype higher than 0.95; (2) the fraction of indels in the reads mapping to the flanking regions of the microsatellite under consideration smaller than 0.15; (3) the fraction of reads estimated to contain a stutter artifact smaller than 0.15; (4) at least 3 sequencing reads spanning each of the supported alleles for the microsatellite under consideration; (5) $\log_{10} P$ value for the allele bias test implemented in HipSTR higher than 2; (6) $\log_{10} P$ value for the Fisher strand bias test higher than 2; (7) the ratio of the number of reads supporting each allele higher than 0.7. This filter served to remove low-VAF mutations likely arising during *in vitro* culture or PCR noise; and (8) a depth of at least 10 sequencing reads. Finally, only microsatellite loci with a reliable call in at least 30 samples and with at least 2 different genotypes across all colonies were considered for further analysis. All mutations satisfying the criteria listed above were further validated through visual inspection of raw sequencing reads.

#### Detection of somatic structural variants

Structural variants were called in each colony using Manta (version 1.6.0), LUMPY (version 0.2.13), SvABA (version 1.1.3), and Delly (version 0.8.3)[9–12]. Each algorithm was run independently on each colony using the bulk sequencing data for stromal cells from the corresponding patient as control, and in a second run using a randomly selected *JAK2*-WT colony as control. The calls generated by each algorithm were merged using the Python library mergevcf (https://github.com/ljdursi/mergevcf) and only calls generated by at least two algorithms were kept for further analysis.

#### Somatic copy number calling

The software package ascatNGS[13] was used to detect somatic copy number alterations in each colony and to estimate their purity and ploidy. The bulk sequencing data from bone marrow stromal cells from the same patient was used as the normal sample in all cases.

#### Mutational signature analysis

Mutational signature analysis was performed using the R package *MutationalPatterns*[14]. To quantify the contribution of mutational processes known to be operative in MPNs[15] (namely SBS1, SBS2, SBS5, SBS19, SBS23, and SBS32) to the observed spectrum of somatic point mutations in each colony, we used the function *fit_to_signatures* using default parameter options. The goodness of fit was determined by computing the cosine similarity between the observed mutational pattern and the reconstructed one using the estimated signature contributions. In all cases we obtained cosine similarity values >0.95, suggesting that our analysis explained most of the variance related to the contribution of different mutational processes to the observed mutational spectra.

**Comparing the mutation rate between *JAK2*-mutant and *JAK2*-WT colonies**

To assess whether the mutation rate in *JAK2*-mutant and *JAK2*-WT colonies is statistically significant, we had to account for the fact that *JAK2*-mutant colonies are clonally related, as they share hundreds of mutations in both ET 1 and ET 2. To account for this shared ancestry, we computed the difference between the mean number of mutations in *JAK2*-mutant and *JAK2*-WT colonies. Next, we computed the expected variance by accounting for the clonal relatedness of *JAK2*-mutant colonies. Specifically, we scaled the variance of the number of mutations in *JAK2*-WT colonies by the number of years at which the clonal expansion started (that is, 9/34 and 19/63 in the case of ET 1 and ET 2, respectively), and computed the square root. We scaled the number of mutations by the variance rather than by the standard deviation given that we assume that the accumulation of mutations in HSPCs can be modelled as a Poisson process. If we then consider the distribution of mean differences to be Gaussian with mean zero, we can compute a *z* score by computing the mean difference divided by the estimated standard deviation, and then estimate the corresponding one-sided *P* value.

**Inference and validation of phylogenetic trees**

The somatic mutations detected across all colonies in a given patient were used to reconstruct phylogenetic trees using the software package PHYLIP version 3.695 (https://evolution.genetics.washington.edu/phylip.html). For each patient and mutation type, namely, SNVs, INDELs, and microsatellite mutations, as well as for these three combined, we constructed a binary matrix with rows indexed by somatic mutations and columns by colonies such that the *i,j* entry in each matrix was set to one if mutation *i* is present in colony *j*, and to zero otherwise. Only mutations detected in at least two colonies were considered to build lineage trees, as private mutations are uninformative to establish the phylogeny of the colonies. We detected a total of 21,699 SNVs (935 present in at least two colonies), 1,396 (60) indels, and 482 (31) microsatellite mutations across the single-cell-colonies derived from patient ET1 (SI Table 2). In the case of ET 2, we detected a total of 33,994 SNVs (1,245), 2,464 (94) indels, and 891 (70) microsatellite mutations (SI Table 2).

For each input mutation matrix, we generated 100 bootstrap replicates by sampling with replacement using the *Seqboot* method. Lineage trees were then estimated for each resample using the Wagner parsimony algorithm as implemented in the *Mix* method using the bulk data from stromal cells as the outgroup. The consensus tree across all bootstrap samples was generated using the extended majority rule method as implemented in the programme *Consense*. Once the consensus tree was determined, we assigned to each branch those mutations that were present in all the descendant colonies of that branch and in none of the other colonies. Lineage tree representations were generated using the R package *ggtree*[16].

As expected given the high number of mutations shared across cancer colonies, the clonal architecture of cancer cells was largely consistent across bootstrap resamples irrespective of the type of mutations considered for lineage tree inference (SI Figs. 18-19). In fact, the clonal architecture for the cancer colonies was the same across all resamples when using somatic SNVs as input. More variability was observed when the lineage trees were constructed using indels or microsatellite mutations as input, although the majority of splits in the tree were consistent across more than 90% of resamples. This is expected given that variant callers generally show lower sensitivity and specificity for the detection of small insertions and deletions as compared to point mutations[17]. This is also consistent with the fact that the highest rates of private INDELs and microsatellite are detected for those colonies with the lowest sequencing quality in our cohort (*e.g.*, HSC-49 from ET 1; SI Fig. 18-19). The clonal architecture of WT colonies varied across resamples, as indicated by the low bootstrap values we obtained for nodes splitting clades of WT colonies. The low concordance observed for node splits across resamples is likely due to the low number of somatic mutations detected in more than one WT colony, consistent with previous lineage tree analyses of human HSPCs using somatic mutations[18]. Overall, the reliability of the consensus trees we have generated is supported by the following: (1) the clonal architecture, in particular for cancer colonies, observed across lineage trees inferred using different types of somatic mutations is overall consistent, (2) the nodes in the trees are largely concordant across bootstrapping resamples for cancer colonies, (3) 96% and 99% of the SNVs detected in at least 2 colonies from ET 1 and ET 2, respectively, could be unambiguously assigned to the consensus lineage tree generated using SNVs, and (4) the single cells in which mutations detected in all *JAK2*-mutant but not in wild-type colonies were identified showed a marked bias towards the megakaryocyte-erythrocyte fate, as opposed to single cells in which these mutations were not detected (Fig. 6a).

**Detection of somatic mutations in the single-cell RNAseq data**

To map single cells to the clades identified in the phylogenetic analysis of WGS data from single-cell derived colonies from patients ET 1 and ET 2, we looked for sequencing reads supporting the mutated allele in the single-cell RNAseq data for all somatic point mutations detected in the WGS data. For each patient and single cell, we extracted all sequencing reads mapping to each position mutated in at least one colony using the Python module Pysam (https://github.com/pysam-developers/pysam)[19]. Only reads with unambiguous cellular barcode ('CB') and molecular barcode ('UB'; i.e., UMI) sequences were further considered. Subsequently, we classified reads as mutant or wild type depending on whether the supported allele, requiring a minimum base quality of 30, was identical to either the alternate or reference alleles, respectively, as determined by the WGS data analysis. All reads with the same molecular barcode were aggregated and the most supported base was assigned to that molecular barcode and considered for further analysis.

To remove mutation calls likely originating from sequencing or library preparation errors we sought evidence for the mutant allele for each mutation across 36 single-cell RNAseq data sets from bone marrow and peripheral blood samples also generated using the 10X Chromium platform. These control data sets include the single-cell RNAseq data we generated as part of this study for MPN patients and healthy controls (Fig. 1b), as well as the following public data sets from the Sequence Read Archive (SRA): SRR6192408, SRR6192409, SRR7244582, SRR7881400, SRR7881401, SRR7881402, SRR7881403, SRR7881404, SRR7881405, SRR7881406, SRR7881407, SRR7881408, SRR7881409, SRR7881410, SRR7881411, SRR7881413, SRR7881415, SRR7881417, SRR7881418, SRR7881419, SRR7881421, SRR7881422, and SRR7881423. All control data sets were uniformly processed using Cell Ranger version 4.0.0 as described above. A total of 1,127,668 UBs mapped to somatic point mutations detected in the WGS data from ET 1 across all controls, of which 1,048,518 (93.0%) supported the reference allele, 10,103 (0.9%) the mutant allele, and 69,046 (6.1%) an allele not observed in the WGS data. In the scRNAseq data from ET 1 128,141 UBs mapped to mutation sites, of which 126,214 (98.5%) supported the reference allele, 1,827 (1.4%) the mutant allele, and 99 (0.15%) an allele not observed in the reference data. In the case of ET 2, a total of 1,712,466 UBs mapped to mutation sites across all controls, of which 1,632,118 (95.3%) supported the reference allele, 10,006 (0.6%) the mutant allele, and 70,341 (4.1%) an allele not observed in the WGS data. In the scRNAseq data from ET 2, we found 73,430 UBs mapping to mutation sites, of which 72978 (99.4%) supported the reference allele, 317 (0.4%) the mutant allele, and 135 (0.2%) an allele not observed in the reference data.

Consistent with previous reports[20], we observed variable error rates across the transcriptome (SI Fig. 27). To eliminate false positives, we discarded all sites with a false positive error rate, defined as the number of molecular barcodes in the controls supporting the mutated allele over the total number of molecular barcodes mapped across all controls, higher than 0.001. For the remaining sites, Fisher's exact test was used to assess the significance of the enrichment for somatic mutation in the single-cell libraries from ET 1 or ET 2 as compared to the controls. The significance level was set to 0.05. A total of 103 somatic SNVs could be reliably detected in the scRNAseq data from patient ET 1, and 96 in the case of ET 2. The greater number of SNVs detected in ET 1 is consistent with the greater sequencing depth used for that patient. These sites were further validated through visual inspection of raw reads, and by confirming consistent phasing with a single allele when the mutations could be phased with nearby heterozygous SNPs (SI Fig. 32).

**Phylodynamic inference**

To infer the clonal expansion of mutant HSCs, we first used BNPR[21], an algorithm that infers population size multiplied by a constant factor from lineage trees, where the constant factor is the time between generations. BNPR assumes a Gaussian process prior on the clonal expansion and infers the marginal posterior distributions of the population size (multiplied by a constant factor) at different time points from the coalescent times of a tree. To infer population dynamics with BNPR, we used the Phylodyn package[22].

For the 34-year-old patient tree, we used an averaging algorithm (the averaging algorithm is described in the ABC sections) to make the length from any leaf to the root of the tree the same. We then converted the branch lengths from mutations to generations by assuming 1 cell division per year[23,24], so that we could infer population size without the constant factor. The coalescent times of the tree were given to BNPR as input. The only parameter we set was lengthout = 28, which determines the number of time slices at which the population size is estimated, and the remaining were default parameters. The BNPR inference on the 63-year-old patient tree was done in an identical manner.

The BNPR inference was not sensitive to the priors we chose. In particular, changing the covariance associated with the Gaussian process did not change the interpretation of the results.

We also tested BNPR on simulated clonal expansions under various scenarios, including simple exponential growth and population bottlenecks, and reliably inferred the population size over time.

**Description of ABC**

Approximate Bayesian Computation, or ABC, is an algorithm used to infer the parameter values of a stochastic model. ABC works by simulating data with the model using parameter values drawn from a prior distribution, and then computing a metric distance between the simulated data and the observed data. If their distance is smaller than a predetermined threshold, the parameter values are retained, otherwise, they are discarded. This procedure is iterated until a sufficient number of parameter values are retained to construct the posterior distribution.

**ABC implementation**

To perform ABC, it is first necessary to define a model. We briefly describe our model (see Supplemental Text for more details), and then give a detailed description of our ABC inference algorithm.

The model we used to infer the population dynamics of mutant cells is a variation of the Wright-Fisher model with selection[25,26]. Briefly, we consider a population of $N$ stem cells that exists in discrete generations. There are $L$ generations in total. At each generation, each cell chooses a parent cell at random from the previous generation. After $t'$ generations, a cancerous mutation is acquired by one of the cells. Critically, the mutant cells are 1 + $s$ times as likely to be chosen as a parent than the wild type cells. As a result, the number of mutant cells grows as ~ $(1 + s)^i$ , for $i$ = 1, … , $g$, where $g = L – t'$ corresponds to the disease duration. For convenience we use g as opposed to t' as a parameter in the following sections. However, provided $L$ is given, if we know the value of g, we also know the value of $t'$ and *vice versa*, and so the two parameters are equivalent. To summarize, the parameters of our model are:

$N$ = saturation parameter (the total number of stem cells)

$L$ = total number of generations

$g$ = disease duration ($g <= L$)

$s$ = selection parameter

We also define $n$ as the number of mutant cells at the final time-point.

We now outline the steps of the ABC algorithm, and then elaborate on the details.

1. Draw $s$ from its prior distribution.
2. Draw $N$, $L$, and $g$ from their prior distributions.
3. Simulate a clonal expansion with our model for $g$ generations.
4. If the final number of mutant cells is $n < k$, where $k$ is the number of mutant cells we sample from the final population, back to 2). Else, move on to 5).
5. Sample $k$ mutant cells from the final population and simulate their lineage history.
6. Simulate the number of mutations along the branches of the tree with the given mutation rate.
7. Perform the averaging algorithm on the tree so that the number of mutations from any leaf to the root of the tree is the same.
8. Convert the tree to an LTT plot.
9. If the area between the LTT plot of our simulated tree and the LTT plot of our patient tree is smaller than epsilon, retain the parameter values, otherwise discard them.
10. If a sufficient number of parameters to construct a distribution has been retained, finish. Else, back to 1).

Note that to perform the ABC, we must first specify the prior distributions on $s$, $N$, $L$ and $g$ (to test the robustness of our simulation *in silico*, we will sometimes fix parameter values rather than drawing from a distribution), the number of cells we will sample from the final population $k$, the mutation rate, and the epsilon threshold for retaining or discarding parameter values.

We now elaborate on the details of each step. We begin with 3) since 1) and 2) simply involve assigning a distribution, which will be specified when the simulations are described below.

After drawing the parameter values in 1) and 2), we simulate a clonal expansion for $g$ generations. By a clonal expansion, we mean the number of mutant cells as a function of time. This can be attained in linear time complexity through a series of binomial draws. We begin with an initial condition of one mutant cell, since the number of mutant cells is always one when the mutation first arises. Then, assuming there are $n(i)$ cells in the $i$th generation, the number of mutant cells in the $(i + 1)$th generation is drawn from a binomial distribution with parameters $N$ and $p = n(i)*(1+s)/(N + n(i)*s)$ (see Supplemental Text for the derivation). After iterating the binomial draw $g$ times beginning with the initial condition, we recover the number of mutant cells as a function of time (SI Fig. 22a). If the mutant clone does not grow to at least $k$ cells, we redraw the parameters in 2) and re-simulate 3), iterating until we have acquired an expansion that does.

It is important to note that the clonal expansion is conditioned on $n >= k$, which is equivalent to conditioning on the mutant cells escaping stochastic extinction. In general, if the clonal expansion is simulated for a sufficient number of generations, that is for a sufficiently large $g$, then the mutant clone will either go extinct or grow to a large size and exhibit deterministic dynamics. Therefore, when g is sufficiently large, if the mutant clone has grown to more than $k$ cells, its size will be much larger than $k$ and will have escaped stochastic extinction. Conditioning on escaping stochastic extinction has the important consequence of allowing us to infer the fitness in percent growth per year from the lineage trees (see Supplemental Text for details).

After obtaining a clonal expansion, we randomly sample $k$ cells from the population of mutant cells at the final timepoint and simulate the lineage history of only the random sample, while ignoring the lineage history of all other cells. The lineage history is constructed by letting each sampled cell choose a mutant cell at random from the previous generation to be its parent. Each mutant cell chosen from the previous generation then chooses its parent at random from the generation of mutants before, etc. This is repeated until all lineages have coalesced (SI Fig. 22b). It is worth noting that simulating the number of mutant cells as a function of time and thus initially ignoring all genealogical relationships, and then simulating only the genealogical history of the random sample backwards in time is statistically equivalent to simulating the genealogical process forward in time, and then producing a tree by following the lineages of the random sample back to common ancestry. This equivalence follows from the fact that each mutant cell can be descended from any of the mutant cells in the previous generation with equal probability. Simulating the lineage history for only the subset of the k chosen cells significantly increases the speed of the simulations without the loss of any information.

Once we have simulated the lineage tree, we then simulate the mutational process. To each edge of the tree (edge refers to a single line connecting two nodes on the tree (SI Fig. 22c), we assign the number of mutations drawn from a Poisson distribution, where the mean is equal to the mutation rate (in units of mutations per generation). The mutation rate is generally computed empirically from the patient tree by dividing the total length of the patient tree in mutations by the value of $L$ - 1, where $L$ was drawn in step 2). It is very important to note that in this case, when we refer to the length of the tree, we mean the total number of mutations from the very bottom of the tree, which corresponds to the present time, to the very top of the tree which corresponds to the birth of the patient (not to the common ancestor of the mutant cells).

Before computing the mutation rate empirically from the data tree, we need to rescale the branches of the tree so that the distance (in mutations) from any leaf to the root of the tree is the same. If we don't do this, the number of mutations from each leaf to the root of the tree would not be the same, resulting in a tree length and mutation rate that is not well-defined. We accomplished this by applying an averaging algorithm described below. The same averaging algorithm is also applied to simulated trees immediately after they are constructed before computing the metric distance between the simulated tree and the data tree. Therefore, any information loss from the algorithm will be expressed as uncertainty in the error bars of our inference.

The averaging algorithm we designed is based on the principle that the best estimate of time to common ancestry between two lineages is the average number of mutations between the two. In particular, let's define a tree as well-averaged if the distance from any leaf to the root is the same. In pseudocode, the algorithm works by calling the following function on the parent of any two sisters:

```
        Average( currentNode )
        {
If the left subtree of currentNode is not well-averaged:
    Average(left child of currentNode)
If the right subtree of currentNode is not well-averaged:
Average(right child of currentNode)
If both the left and right subtrees are well-averaged:
Compute the average length of the left and right subtrees. Then, for both the left and right subtree, rescale the branches of the subtree proportionally so that the length of the subtree equals the average.
    if currentNode != root:
        Average( parent of currentNode)
    else:
        break
}
```

We begin at the parent of two sisters, where the subtrees are single branches connecting a parent node to two leaf nodes. Note that we may start at the parent of any two sisters (or even more generally, at any node) and produce the same averaged tree since averaging the two subtrees of any node produces a unique value. For implementation of this algorithm refer to our GitHub repository. See SI Fig. 22d for a schematic of the averaging algorithm.

After producing a well-averaged tree, we construct its LTT (Lineages Through Time) plot. The LTT plot of a tree shows the number of lineages as a function of time in mutations (SI Fig. 22e). The LTT plot of a tree loses all information about its topology (the way the branches are connected). However, since the mutant cells in each generation pick their parents at random from the mutant cells in the previous generation, any topology on the tree is equally likely, and thus the tree topology contains no information about the parameter values that gave rise to the tree. Therefore, LTT contains all possible information about the parameter values.

After converting the simulated tree to an LTT plot, we compute the distance between the LTT plot of the simulated tree to the LTT plot of the data tree, defined as the area between the two LTT curves. The LTT plot of the data tree is always constructed before the ABC begins by first applying the averaging algorithm we previously described so that the leaf nodes line up side by side, and then converting it to an LTT plot. If the area between the two plots is smaller than the epsilon threshold, we retain the parameter values $s$, $N$, $L$, and $g$ drawn from the priors, as well as the cancer trajectories $n(t)$ and the LTT curve produced, and if the area is >= epsilon we discard them. This process is iterated until a sufficient number of parameter values (along with the trajectories and LTT plots) to construct a convergent posterior distribution is retained.

The LTT curves start at zero but may end at different values because of different tree lengths. The area between two LTT curves that do not end at the same point on the x axis is undefined. To address this, we extend the end points of LTT curves, which corresponds to a value of 1, to infinity.

Since the lengths of LTT curves tended to vary, we decided to divide the area by *k**(the length of the data tree) before checking if epsilon was smaller than the threshold. This allowed us to run ABC without having to choose a new epsilon for each tree, since a smaller epsilon would be required for a tree of smaller length, and a larger epsilon for a tree with a larger length. Intuitively, this is equivalent to taking the percent difference between the data tree and the simulated trees.

**Simulating data**

To test the robustness of our ABC inference, we were interested in inferring the model parameters from simulated data where the ground truth is known. The way we simulated data as follows:

1. Draw $s$ from its prior distribution.
2. Draw $N$, $L$, and $g$ from their prior distributions.
3. Simulate a clonal expansion for g generations.
4. If the final number of mutant cells $n < k$, where $k$ is the number of mutant cells, we sample from the final population, back to 2). Else, move on to 5).
5. Sample $k$ mutant cells from the final population and simulate their lineage history.
6. Simulate the mutational process on the tree with the given mutation rate.
7. Perform the averaging algorithm on the tree.

The resulting tree is used as the data tree in ABC, and its parameters are inferred. Note that these steps are simply the first 7 steps of ABC.

***In silico* validation of the inference algorithm**

To validate our inference, we decided to test the inference on simulated data over a wide range of parameter values. We began by simulating 30 trees as data with our model, where the underlying parameter values were known. Each tree was constructed using the following specifications:

1. $s$ was drawn from a uniform distribution on (0, 1.2).

2. $N$ was drawn from $10^X$, where $X$ is uniformly distributed on (1, 9).
3. $L$ was drawn from round($Y$), where $Y$ is a Gaussian with mean 35 and std 5. If we drew $L$ < 2, we redrew $L$ until L >= 2 since at least 2 generations are necessary to produce a tree.
4. $g$ was drawn uniformly on 2, … , $L$.
5. $k$ = 22 mutant cells were randomly sampled. (22 is the number of mutant stem cells sampled for ET 1 patient data)
6. The mutation rate was 723/($L$ - 1). 723 was the number of mutations observed in ET 1 patient data.

We then inferred the parameters for each tree using ABC. For the ABC, we used the exact same specifications as the data to generate trees for comparison, except that we instead drew $s$ from a uniform distribution on (0, 5) in Step 1, and the mutation rate was instead estimated empirically as (total length of tree in data in # of mutations)/($L$ - 1) in Step 6. Epsilon was set to 0.03, since this threshold was sufficient to obtain an inferred distribution that converges to the posterior distribution for most inferences, while also allowing a large number of points to be retained for the inferred distribution. The simulations were run until they accrued ~10,000 or more points for the posterior.

In SI Fig. 23, we show a representative set of 10 inferences out of the 30 inferences we ran.

To quantify the accuracy of our ABC inference, we then simulated 200 trees in precisely the same way as above, except that the value of $s$ for the tree data was drawn uniformly on (0, 2) instead to obtain data across a much wider range of fitness values. We then carried out ABC inferences on each tree with epsilon = 0.0225 until most of the posterior distributions had accrued ~400 or more points. Tree data where the inference accrued less than 30 points for the posterior were excluded.

We then applied the following filters to the data:

1. We excluded data trees where the ratio of the standard deviation to the mean of the posterior of $s$ (in percent growth per year) was greater than 0.425

2. We excluded data trees where the std of *n* was larger than 1.15

We arrived at the first filtering criterion by noting that inferences for small s tended to have large error bars relative to their inferred means (or large coefficient of variation), and their inferred means were generally inaccurate and much larger than the true values. ABC inference cannot determine whether a small number of cells at the final time-point is due to a small growth rate s or small saturation limit (see the inferred trajectories in SI Fig. 23 row five column one and SI Fig. 24 row two column 2). In both scenarios the population size is small and coalescence events occur rapidly, producing similar trees. We reasoned that if expansions produced by small s produce ABC inferences characterized by large coefficient of variation, then by eliminating ABC inferences exhibiting this characteristic we could exclude inaccurate inferences without any knowledge of the ground truth.

Similarly, we arrived at the second filtering criterion because simulations that expanded to sufficiently large population sizes generated inferred log(*n*) distributions with large standard deviations and mean values distributed around $10^6$. This suggested that ABC was extracting little information from the data and that the log(*n*) distributions were almost identical to the prior. We reasoned that by filtering out inferences with large log(*n*) standard deviations we could exclude inaccurate inferences without any knowledge of the ground truth value. We emphasize that this filtering procedure does not use the ground truth values in any way. The inference is deemed inaccurate if the posterior distribution width is too large regardless of the ground truth value. Therefore, the filtering procedure can also be applied to actual data where the ground truth is not known. Finally, when devising the filtering criteria we were conservative with our choices. As such, the interpretation of these data was not sensitively dependent on the filters we chose.

The inferred vs true values of the inferences is plotted in SI Fig. 25a.

Next, we quantified the accuracy of our ABC inference for a 63 year old patient in a similar manner by simulating 200 trees, applying filters to the data, and then plotting the inferred vs true values (SI Fig. 25c). The data were produced in a similar manner as for the 34 year old patient using the following criteria:

We simulated 200 trees as data for a 63 year old patient using the following specifications:

1. $s$ was drawn from a uniform distribution on (0, 2).
2. $N$ was drawn from $10^X$, where $X$ is uniformly distributed on (1, 9).

$L$ was drawn from round($Y$), where $Y$ is a Gaussian with mean 64 and std 10. If we drew $L < 2$, we redrew $L$ until L >= 2 since at least 2 generations are necessary to produce a tree.

3. $g$ was drawn uniformly on 2, … , $L$.
4. we sampled $k$ = 13 cancer cells, which is the number of sampled cells in the ET 2 patient data.
5. The mutation rate was 1205/$L$. 1205 was the number of mutations observed in ET 2 patient data.
6. No feedback was included.

We then carried out ABC inferences on each tree using the same specifications as the data, except that we drew s from (0, 5) uniformly in Step 1, and the mutation rate was estimated empirically as (total length of tree in data)/($L$-1) for 6). Epsilon was set to 0.0125. The inferences were left running until about half of them (many ABC inferences accrued little to no points for the inferred distributions) had accrued ~100 or more points for the posterior distribution. Many simulations accrued little or no points, and so we excluded trees with less than 30 points.

We then applied the following filters to the data:

1. We excluded data trees where the mean of the posterior of s (in percent growth per generation) was larger than 1.5.
2. We excluded data trees where the ratio of the std to the mean of the posterior of s (in percent growth per year) was greater than 1.5.

Similar reasoning was applied to devise the above filtering criteria. Mainly, values outside of above criteria contain little information beyond the prior distributions.

**ABC on simulated data with feedback**

Our model of growth dynamics of the mutant cells only approximates the actual growth dynamics. In particular, the population of mutant cells seems to saturate when it has expanded to a certain fraction of the total population of stem cells. Therefore, it is conceivable that the mutants lose their fitness advantage as their population size increases. Here, we set out to test whether the inference of the parameters of the simple model of growth dynamics remains accurate if the actual dynamics is simulated using a different model. To do so, we constructed a model with feedback, whereby the fitness advantage of the mutant cells decreases compared to wild-type cells as their population size increases. The model with feedback is described in detail in the Supplemental Text.

We then repeated the simulations carried out for 34-year-old and 63-year-old patients, except that we used feedback with $x = 30$ when generating the simulated data (see Supplemental Text for definition of x parameter). The ABC did not incorporate feedback in the model, since we were interested in how well we could infer the parameter values if the ground truth incorporated feedback. For the 10 example trees shown in SI Fig. 24 we ran ABC until most of the inferences had accrued ~5,000 or more points for the posterior. For the 200 trees for the 34 year old (SI Fig. 25b), we ran ABC until most of the inferences had accrued ~300 or more points for the posterior. For the 200 trees for the 63 year old (SI Fig. 25d), we ran ABC until about half of the inferences had accrued ~50 or more points for the posterior. Inferences that accrued less than 30 points were always excluded. The filters were applied in an identical fashion as for the inferences without feedback.

Taken together, the inferences suggest that the ABC inference can infer model parameters over a wide range of parameter values, regardless of whether or not feedback is incorporated in the underlying model. In particular, *s* in percent per year and the age of onset of the disease can be inferred from lineage trees, even if feedback is incorporated. However, it appears n can only be inferred for the 34-year-old patient. The inferred *n* vs true *n* plots for the 63-year-old patient indicate that the trees have no information about the number of mutant cells at the final time point. This is likely due to the fact that we have sampled only 13 lineages (as opposed to 22 lineages for the 34 year old), and that the clone has expanded for much longer. Because of this, most coalescent events occur in the early history of the expansion, and information about the dynamics of the later history are lost.

**Fitness can be inferred without knowing the number of generations**

So far, we have shown that our inference is robust to feedback but have not shown how well we can infer the parameter values if our assumption about the total number of generations is incorrect. Surprisingly, fitness, when converted to percent growth per year, can always be inferred without knowing the number of generations (see Supplemental Text). To validate this, we simulated ~10 data trees for a 34 year old patient, and carried out ABC on them. We then selected a data tree where the ABC inference precisely inferred the parameter values. The data tree had been simulated with the following specifications:

1. Parameter values were fixed at $s = 0.264911$, $N = 10^9$, $g = 50$, $L = 70$, arbitrary values for which the inference was accurate.
2. $k = 22$ cells were sampled
3. The mutation rate was 723/69 per generation

We then inferred *s*, *n*, and the age of onset of the disease having kept all other parameter values fixed, but assuming that the number of generations *L* was *c* times 70 (the ground truth *L*) in the ABC model. More precisely, for each $c = 0.5, 1, 2, 4, 8, 16, 32, 64$ we ran an ABC inference on the data tree with the following specifications:

1. $s$ was drawn uniformly on (0, 10/$c$)
2. We fixed $N = 10^9$ and $L = c*70$
3. $g$ was drawn uniformly on 2, … , $L$
4. A mutation rate of 723/(70*$c$ - 1) was used
5. An epsilon distance of 0.02 was used

The inferences were run until the ABC had accrued ~15,000 points for the posterior distribution. We then plotted inferred joint distributions for $s$ in growth per generation, $s$ in growth per year, age of onset of the disease, and $n$ (SI Fig. 21).

As expected, when increasing the number of generations assumed by the model for ABC inference, the inferred $s$ in percent growth per generation decreased while the inferred percent growth per year remained invariant. In theory, the decrease in growth per generation will increase the rate of stochastic extinction, and so the number of mutant cells must fluctuate to a larger population size to escape stochastic extinction. As expected, the number of mutant cells increased at the final time point.

Taken together, our simulation results are consistent with our theoretical calculations (Supplemental Text) in that fitness, in percent growth per year, and the age of onset in years can be inferred from lineage trees without prior knowledge of $L$, while prior knowledge of $L$ is necessary to infer $n$.

**The analytical calculation of coalescent times matches the simulations results**

In the Supplemental Text, we provide an analytical calculation of the average coalescent times of our model, and show that the average coalescence times do not change if we scale the number of generations while keeping the percent growth the same (suggesting that fitness can be inferred without prior knowledge of the number of generations).

To validate our analytical calculation of coalescent times, we performed the following simulations. For each $s$ = 0.1, 0.3, … 1.5 (growth per generation), we constructed thousands of data trees using the following specifications:

1. $g$ = 25, $L$ = 35, $N = 10^9$ (with the corresponding $s$)
2. $k$ = 22 cells were randomly sampled
3. A mutation rate of 723/34 was used

We then converted the branches of each data tree to years, assuming the tree was for a 34 year old patient, by multiplying the branch lengths by 34 / 723. Then, for each value of s separately, we constructed a distribution for each of the i = 1, … , 21 coalescence times using the corresponding trees. We computed the means and standard deviations of those distributions and plotted them (SI Fig. 20a).

The analytical derivation of coalescence times therefore matches the simulated coalescence times within a standard deviation. Our derivations also predict that the coalescence times of a tree should not change when scaling the number of generations while keeping the percent growth per year fixed. To verify this occurs in our simulated trees, we did the following:

For each $s'$ = 0.1, 0.6, 1.1 and for each $c$ = 0.5,1, 2, 10, 100, we simulated thousands of data trees using the following specifications:

1. $s = (1 + s')^{1/c} - 1$ (this $s$, in growth per generation, along with the $L$ in specification 2), keep the percent growth per year invariant (See supplemental).
2. $L = c*35$, $g = c*25$, $N = 10^9$,
3. 22 cells were randomly sampled
4. A mutation rate of 723/($c$*35 - 1) was used

We then converted the branches of each data tree to years, assuming the tree was for a 34 year old patient, by multiplying the branch lengths by ($c$*35 - 1)/723. For each combination of $s$ and $c$, we constructed distributions for each of the $i = 1, \ldots, 21$ coalescence times of the corresponding trees. We then computed the means and standard deviations of the distributions and plotted them (SI Fig. 20b).

Consistent with our theoretical predictions, scaling the number of generations while keeping the percent growth per year invariant appears to not significantly change the average times until coalescence. This implies that trees are indistinguishable when the percent growth per year is the same, even if the number of generations is different, showing that percent growth per year can be inferred from lineage trees without knowing $L$.

**Inference on patient data**

On each patient tree, we ran our ABC algorithm and inferred the model parameters.

For the patient with age 34, we used the following specifications for ABC:

1. $s$ was drawn from a uniform distribution on (0, 2).
2. $N$ was drawn from $10^X$, where $X$ is uniformly distributed on (1, 9).
3. $L$ was drawn from round($Y$), where $Y$ is a Gaussian with mean 35 and std 5. If $L < 2$, we redrew $L$ until $L >= 2$ since at least 2 generations are necessary to produce a tree.
4. $g$ was drawn uniformly on $2, \ldots, L$.
5. $k = 22$ cells were sampled
6. the mutation rate was (total length of patient tree in mutations)/($L$ - 1). In this case, the total length of the patient tree in mutations was 723.
7. An epsilon threshold of 0.0225 was used.

For the patient with age 63, we used the following specifications for ABC:

1. $s$ was drawn from a uniform distribution on (0, 2).
2. $N$ was drawn from $10^X$, where $X$ is uniformly distributed on (1, 9).

3. $L$ was drawn from round($Y$), where $Y$ is a Gaussian with mean 65 and std 10. If $L < 2$, we redrew $L$ until $L >= 2$ since at least 2 generations are necessary to produce a tree.
4. $g$ was drawn uniformly on 2, … , $L$.
5. $k$ = 13 cells were sampled
6. The mutation rate was (total length of patient tree in mutations)/($L$ - 1). In this case, the total length of the patient tree in mutations was 1205.
7. An epsilon threshold of 0.0125 was used.

Note that we drew $s$ from (0, 2) instead of from (0, 5). This was done to speed up the simulations and is justified because preliminary runs showed the distribution of s converging to a much smaller value.

For each patient we ran 400 parallel simulations. For the 34-year-old we collected 1,038,712 data points from the posterior, and for the 63-year-old we collected 8,816,199 data points from the posterior. The posterior joint distributions are plotted in Fig. 4d and SI Fig. 26.

As indicated by our analysis, fitness $s$ could be inferred from the patient trees. Our analysis also suggests that if we assume a division rate of one per year[18], $n$ can be inferred for the 34 year old patient as 4.74 +/- 0.68 of the posterior distribution. $n$, however, cannot be inferred for the 63-year-old patient, and the inferred distribution of $n$ is just the prior information. This is due to the fact that the coalescent events occur in the very early history of the disease, and the information about the population size is lost. We can, however, put bounds on the point of saturation if we assume one division per year. As seen before, when the number of mutant cells approaches $N$, the growth of the mutant population slows down and starts to exhibit neutral dynamics. If $N$ is sufficiently small, it changes the coalescent structure, and ABC then assumes the saturation point is $n = N$. Trajectories generated by ABC if $N$ is below a certain threshold value produced coalescent structures that did not match that of the patient data. Any value of $N$ larger than this threshold had no effect on the coalescent structure and therefore was retained as a possible inferred value. Therefore, $N$ could not be precisely determined.

**Code availability**

We developed a C++ object called StemCellSim for simulating clonal expansions and inferring the parameters of our model. StemCellSim has the ability to generate simulated data under various models, and to infer model parameters from either simulated or real data with ABC. The code can be found on GitLab (https://gitlab.com/hormozlab).

**Data availability**

Raw scRNA-seq and whole-genome sequencing data will be deposited in dbGAP prior to publication.
